# Interkingdom glycine conjugates of indole-3-carboxylates are Ah receptor ligands

**DOI:** 10.64898/2026.02.13.705761

**Authors:** Ethan W. Morgan, Andrew J. Annalora, Denise M. Coslo, Krishne Gowda, Dhimant Desai, Fangcong Dong, Ethan W. Davis, Iain A. Murray, Fuhua Hao, Imhoi Koo, Kristina S. Petersen, Penny M. Kris-Etherton, Trenton Wolfe, Reece Erickson, Seth T. Walk, Jordon E. Bisanz, Shantu G. Amin, Craig B. Marcus, Andrew D. Patterson, Gary H. Perdew

## Abstract

Substituted indoles are conserved metabolites across all kingdoms of life and may function as a mediators of inter- and intra-species communication. Indole-3-carboxylates (indole-3-acetic acid (IAA) and indole-3-propionic acid (IPA)) represent abundant tryptophan-derived AHR agonists in human serum, potentially influencing AHR-dependent physiology. LC-MS analysis of mouse serum, urine and cecal/fecal contents reveals that both IAA and IPA undergo host and microbial mediated glycine conjugation to facilitate urinary elimination. Notably, at physiologically detectable human serum concentrations (μM), IAA-Glycine retains human AHR activation potential. Comparative *in silico* docking simulations corroborate IAA-Glycine as a direct ligand for the human AHR. Data suggest, in contrast to xenobiotic ligands, AHR activation by endogenous tryptophan metabolites is greater in humans than in mice. These results underscore the role of microbial and host-derived amino acid conjugation in generating bioactive metabolites. Thus, positioning interkingdom auxin chemistry within human physiology and revealing an unexpected link between plants, microbes, and humans.

## Introduction

The aryl hydrocarbon receptor (AHR) is the only member of the basic helix-loop-helix Per-Arnt-Sim family of transcription factors that is activated by a wide array of small molecule ligands^1^. In its unliganded state, the AHR resides in the cytoplasm as a complex with heat shock protein 90 (HSP90) and X-associated protein 2 (XAP2) that undergoes dynamic nucleocytoplasmic shuttling^2,3^. The structure of the AHR/HSP90/XAP2 complex was recently confirmed using cryogenic electron microscopy^4^. Upon agonist binding, the AHR translocates into the nucleus, where it heterodimerizes with the aryl hydrocarbon receptor nuclear translocator (ARNT). This heterodimer binds to dioxin response elements and regulates a diverse array of genes, offering insights into the physiological functions of this enigmatic receptor. In addition to canonical signaling, the AHR also modulates signaling through non-canonical pathways, underscoring the complexity of AHR-mediated functions^5^.

The AHR is regarded as an environmental sensor that mediates responses critical to barrier tissue homeostasis^6,7^. For example, AHR activation by dietary ligands in the gut ameliorates *Clostridioides difficile* infection^8^, while AHR activation in keratinocytes promotes differentiation and stratum corneum development, thereby enhancing skin barrier function^9,10^. In the intestinal tract, AHR directly regulates the cytochrome P450 1A1 (CYP1A1), a metabolic enzyme that limits absorption of dietary AHR ligands, acting like a metabolic barrier^11^. We have previously established that AHR expression and activity are highest in the duodenum and decline along the gastrointestinal tract toward the colon.^12^ Duodenal AHR activation promotes epithelial differentiation and suppresses proliferation. Additionally, AHR activation in the small intestine induces production of antimicrobial peptides and enhanced goblet cell production, further supporting barrier function^12–14^. Collectively, these findings illustrate that the AHR is a vital regulator of both normal barrier tissue physiology and responses to chemical or microbial insults.

The AHR is a promiscuous receptor capable of binding structurally diverse ligands originating from dietary sources, microbial metabolism, or host metabolic pathways^15,16^. Early studies focused on relatively high-affinity xenobiotics such as 2,3,7,8-tetrachlorodibenzo-*p*-dioxin (TCDD), planar chlorinated biphenyls, and polycyclic aromatic hydrocarbons^1,16^. More recently, attention has shifted to Trp (tryptophan) metabolites produced by the host and microbiota^17^. Various Trp metabolites are made by the gut microbiota at concentrations sufficient to activate the AHR and have demonstrated therapeutic potential^18–20^. Notably, many of these metabolites, including indole, indole-3-carboxylic acid, kynurenic acid (KA), are more potent activators of the human AHR than the mouse receptor^17,18,21^. In mammals, systemic abundance of the essential amino acid tryptophan is regulated primarily through the kynurenine (KYN) pathway, which is initiated by hepatic 2,3-tryptophan dioxygenase (TDO) activity and results in multiple downstream metabolites, including NAD^+22,23^. KYN, KA and xanthurenic acid—products of the KYN pathway—are AHR ligands^24^. Indole dioxygenase (IDO), expressed in immune cells, can contribute to local tryptophan depletion through metabolism to KYN and the circulating KYN concentration.

Recently, we have demonstrated significant concentrations of tryptophan-derived AHR ligands circulating in mice and humans, which are produced by both the host and its microbiota^15^. A comparison of serum from germ-free and conventional mice revealed that the presence of KYN, KA, indole-3-lactic acid, indole-3-carboxaldehyde, indole-3-acetic acid (IAA) and 5-hydroxindole-3-acetic acid do not depend on the presence of gut microbiota. Indeed, IAA is known to be made by both host and gut microbial metabolism^15,18^. The source of host generated IAA has not been explored in detail but is likely an enzymatic or spontaneous product of indole-3-pyruvic acid, formed through the metabolic deamination of tryptophan by IL4I1 amino acid oxidase or transaminases (e.g. glutamic-oxaloacetic transaminase 1/GOT1)^25,26^. IAA was shown to attenuate various parameters associated with metabolic dysfunction-associated steatotic liver disease in mouse models^27,28^. In addition, IAA production in the intestinal tract has been linked to an AHR-PARP1 axis to enhance DNA repair in aging mice^29^. Other microbiota-derived Trp metabolites found in serum, such as indole-3-propionic acid (IPA), 2-oxindole and indole-3-acrylic acid, activate the AHR. The mean serum concentrations of IPA are 3 μM in humans, and 5 μM in mice, suggesting that IPA may exhibit pharmacologic potential^15^. IPA is a potent antioxidant exhibiting a variety of positive health effects such as attenuating various metabolic disease endpoints and neuroprotective properties^30,31^. Additionally, IPA can activate the pregnane X receptor and is associated with improved outcomes in sepsis, intestinal inflammation, and barrier function^32–34^.

Plants, microbes and humans use a common set of signaling molecules that shape physiology across species. Despite growing interest in these Trp metabolites, the host and microbial metabolic fate and clearance mechanisms remain poorly understood. Therefore, in this study, we investigated the metabolism of IAA and IPA via glycine conjugation and assessed how this modification modulates AHR activation potential. Notably, we found that glycine-conjugated IAA retains the human AHR activation potential in its unconjugated form. *In silico* docking revealed that IAA and IAA-glycine adopt an orientation in the ligand binding pocket similar to that of indirubin, a high-affinity AHR ligand. In contrast, glycine conjugation of IPA markedly reduces the activation potential of the human AHR. These results suggest that tryptophan deamination products influence basal systemic AHR activity, not only through the production of IAA, but also through the presence of its glycine-conjugated form. The identification of functional glycine conjugated indolic AHR ligands represents a previously unrecognized addition to the AHR ‘exposome’. Such interkingdom (host, microbe and plant) availability of AHR active conjugates, arising through interdependent host-microbe metabolism or through dietary absorption of indolic conjugates abundantly stored in plants, increases the complexity of the AHR ligand reservoir and thus may modulate physiological AHR activity. Furthermore, these findings indicate that plants, microbes and humans share a common chemical language that shape physiology across species.

## Results

### IPA and IPA-Glycine presence in mice and humans

IAA and IPA are two notable Trp metabolites that can increase AHR activation potential (Fig. 1A). A previous report has shown the presence of IPA conjugated with glycine through the acyl-CoA synthetase medium chain family member 2A/glycine N-acyltransferase (ACSM2A/GLYAT) pathway^35^. We used LC-MS with recovery and matrix-effect adjustments to quantify circulating IPA and IPA-glycine (IPA-Gly) (Fig. 1B). Consistent with being an obligate microbial metabolite, serum IPA levels were below the limit of detection in germ-free mice, thus confirming their status. In contrast, conventional mice on a standard laboratory chow diet exhibited a mean serum IPA concentration of 7.2 μM compared to mice on a semi-purified AIN-93G (AIN) diet, essentially phytochemical-free, which exhibited a significantly reduced mean serum concentration (0.27 µM). Consistent with the low abundance observed for IPA, the mean serum IPA-Gly followed a similar pattern: 2.8 µM in chow-fed mice compared to a significantly reduced to 0.7 μM in mice when on the AIN diet (Fig. 1B). Analysis of mouse cecal contents revealed an absence of IPA in germ-free mice, again supporting the microbial-derived origin of IPA (Fig 1C). In conventional mice, cecal IPA levels mirrored those observed in serum, with a significantly elevated mean abundance in chow fed mice relative to those on the AIN diet (Fig. 1C). Analysis of cecal IPA-Gly abundance across germ-free and conventional chow-fed mice as well as conventional mice on an AIN diet, revealed a similar pattern to that obtained in serum (Fig. 1C). Analysis of feces and urine in conventional, chow-fed mice identified opposing metabolite quantification. While IPA is quantifiable in feces, IPA-Gly is not detectable, and the converse is apparent in mouse urine (Fig. 1C-D). These data are consistent with urinary excretion, rather than fecal, as the primary mode of IPA elimination through glycine conjugation. Examination of IPA and IPA-Gly levels in renal tissue indicates the presence of both metabolites in conventional chow-fed but not germ-free mice (Fig. 1E). While in liver tissue only IPA was detected (Fig. 1E).

**Figure 1.**
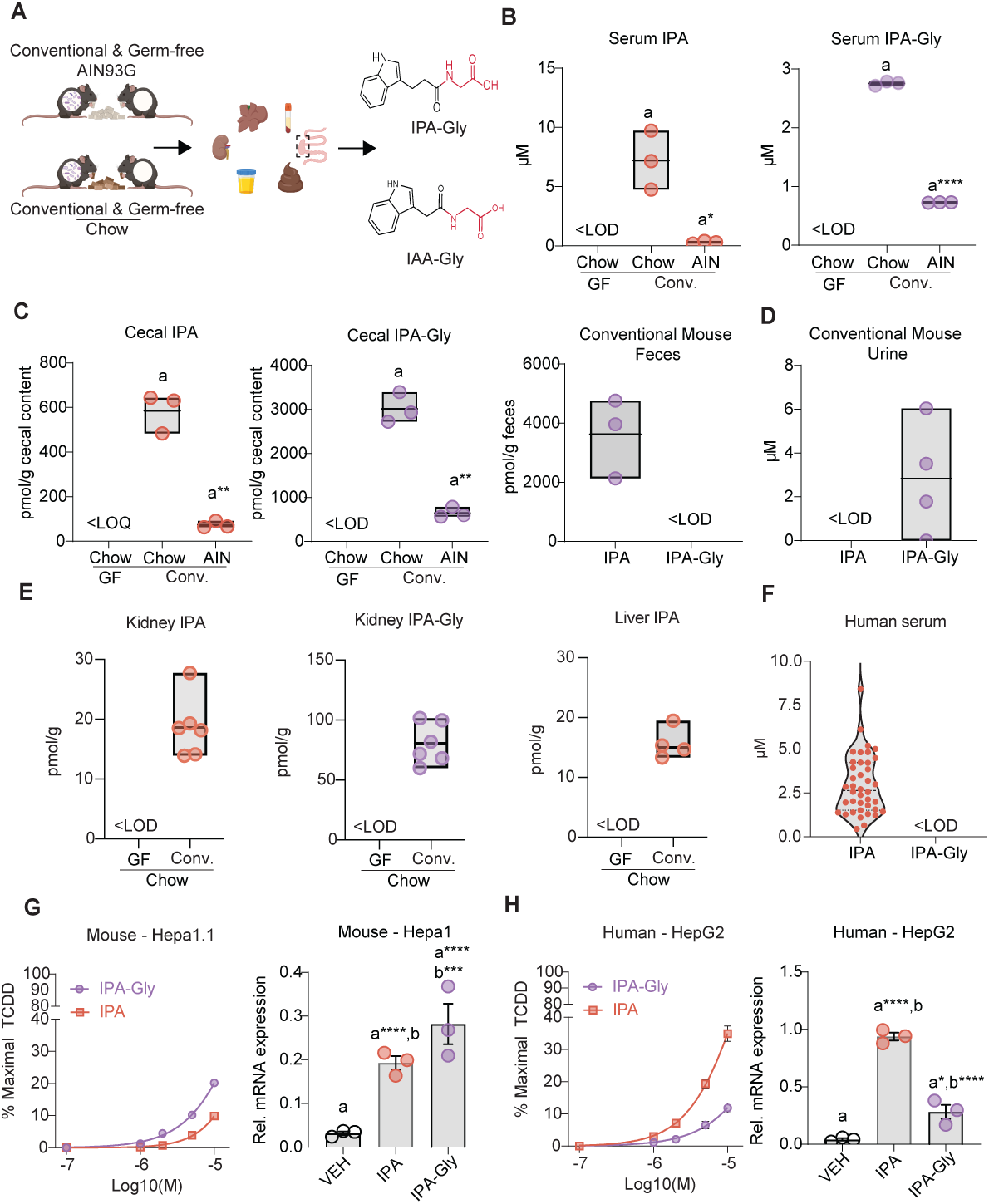
Indole-3-propionylglycine is present in mouse bio samples and exhibits increased AHR activation potential. **A.** Chemical structures of IPA, IAA and their respective glycine conjugates (IPA-Gly and IAA-Gly). **B.** Quantification of IPA and IPA-Gly in serum collected from germ-free (GF) mice (n=5) and conventional (Conv.) mice fed a chow diet (Chow., n=3) or the phytochemical-depleted AIN93G diet (AIN, n=3). **C.** Quantification of IPA and IPA-Gly in metabolite extracts from cecal contents of GF mice fed chow, and Conv mice fed Chow or AIN diet, and feces from Conv mice (n=3). **D.** Quantification of IPA and IPA-Gly extracted from Conv mouse urine (n=4). **E.** Quantification of IPA and IPA-Gly in kidney and liver samples collected from GF (n=5) and Conv. mice (n=6). IPA-Gly was below the limit of detection (<LOD) in all liver samples. **F.** Serum from humans fed a defined diet contained quantifiable concentrations of IPA, but IPA-Gly was not detected (n=40). **G.** AHR activity in mouse cell lines was assessed by luciferase activity normalized to the response produced by a saturating dose of TCDD (10 nM) in Hepa 1.1 cells and *CYP1A1* expression upon exposure to 20 μM of each ligand in Hepa 1 cells (n=3). **H.** AHR activity in human cell lines was assessed by luciferase activity normalized to the response produced by a saturating dose of TCDD (10 nM) in HepG2 40/6 cells and *CYP1A1* expression upon exposure to 20 μM of each ligand in HepG2 cells (n=3). Data are represented by mean ± S.E. or box and whisker plots. Cells were exposed to treatments for 4 h in serum-free media. Statistical comparisons were made using a *t-*test or one-way ANOVA. *: p < 0.05, **: p < 0.01, ***: p < 0.001, ****: p < 0.0001. Alphabetical characters indicate statistical comparisons between 2 groups.

To determine if glycine conjugation of IPA occurs in humans, we examined overnight-fasted serum from a cohort of subjects maintained on a defined diet (Fig.1F). LC-MS analysis across 40 subjects reveals IPA concentrations ranging from 400 nM to 8.4 μM (%CV = 57%) with a mean serum concentration of 2.9 ± 1.7 μM (Fig. 1F). Despite the variability in serum IPA, IPA-Gly levels across the cohort were below the limit of detection (Fig. 1F).

### IPA and IPA-Gly AHR activation potential

We have previously established that, relative to other Trp metabolites (i.e., KYN, indole-3-acetic acid), IPA is a relatively weak human AHR agonist^15^. To determine whether glycine conjugation of IPA impacts its capacity to activate AHR transcriptional activity, mouse AHR-dependent reporter assays were performed using Hepa1.1 cells. Consistent with previous studies, IPA activated mouse AHR in a dose-dependent manner, achieving 10% maximal TCDD-mediated induction at the highest concentration examined (10 μM) (Fig.1G). IPA-Gly also elicited dose-dependent mouse AHR activation, achieving 20% maximal TCDD-mediated induction at the highest concentration examined (10 μM) (Fig. 1G). The elevated capacity of IPA-Gly over IPA to induce mouse AHR activity was corroborated by assessing *Cyp1a1* inducibility in Hepa1 cells. (Fig. 1G). Although not detected in human serum samples, IPA-Gly human AHR activity was assessed using HepG2(40/6) reporter cells. In contrast to mouse AHR, data reveal IPA-Gly has reduced capacity to activate human AHR relative to IPA (Fig. 1H). The reduced potential of IPA-Gly was confirmed by assessing *CYP1A1* induction in HepG2 cells (Fig. 1H). Given that IPA-Gly manifests as a weak human AHR agonist, even in comparison to IPA, coupled with the observation that IPA-Gly was not detected in human serum, we did not further characterize this metabolite.

### IAA metabolism and conjugation in mice

We next explored the more potent (relative to IPA) AHR ligand, IAA, which also undergoes glycine conjugation in vivo (Fig. 2). LC-MS analysis of serum from germ-free mice quantified the mean serum concentration of IAA and IAA-Gly at 1.0 and 1.5 μM respectively (Fig. 2A). These data suggest that the presence of IAA and its conjugation with glycine to IAA-Gly are not dependent upon the microbiota and can be host facilitated. Comparison between germ-free and conventional mice on either chow or AIN diet revealed mean serum concentrations of IAA to be similar (Fig. 2A). In contrast, the mean serum concentrations of IAA-Gly across these groups were significantly different, with conventional chow-fed mice exhibiting elevated IAA-Gly (2.7 μM) compared to both germ-free mice (1.5 μM) and mice on the AIN diet-fed counterparts (0.7 μM) (Fig.2A). These data suggest that, while microbial status and diet do not appear to influence serum IAA, they individually, or in combination, affect IAA conjugation with glycine. Although not absolutely required, the microbiota likely contributes to the circulating levels of IAA-Gly. Analysis of cecal contents showed significantly elevated levels of IAA in conventional mice (161.6 pmol/g) compared to germ-free mice (9.9 pmol/g) (Fig. 2B). Additionally, the concentration of IAA in cecal contents in conventional mice fed an AIN diet was determined to be significantly diminished (21 pmol/g). Analysis in the cecal contents of germ-free and conventional mice in addition to mice on an AIN diet revealed IAA-Gly levels to be <LOD, 552.8 and 192 pmol/g, respectively (Fig. 2B). These results indicate that there is 3.4-fold more IAA in the cecum conjugated with glycine compared in IAA in conventional mice on a chow diet. In conventional mice on chow diet, both IAA and IAA-Gly were also present in urine and feces with urinary IAA-Gly levels exceeding that of IAA by > 9-fold (Fig. 2C).

**Figure 2.**
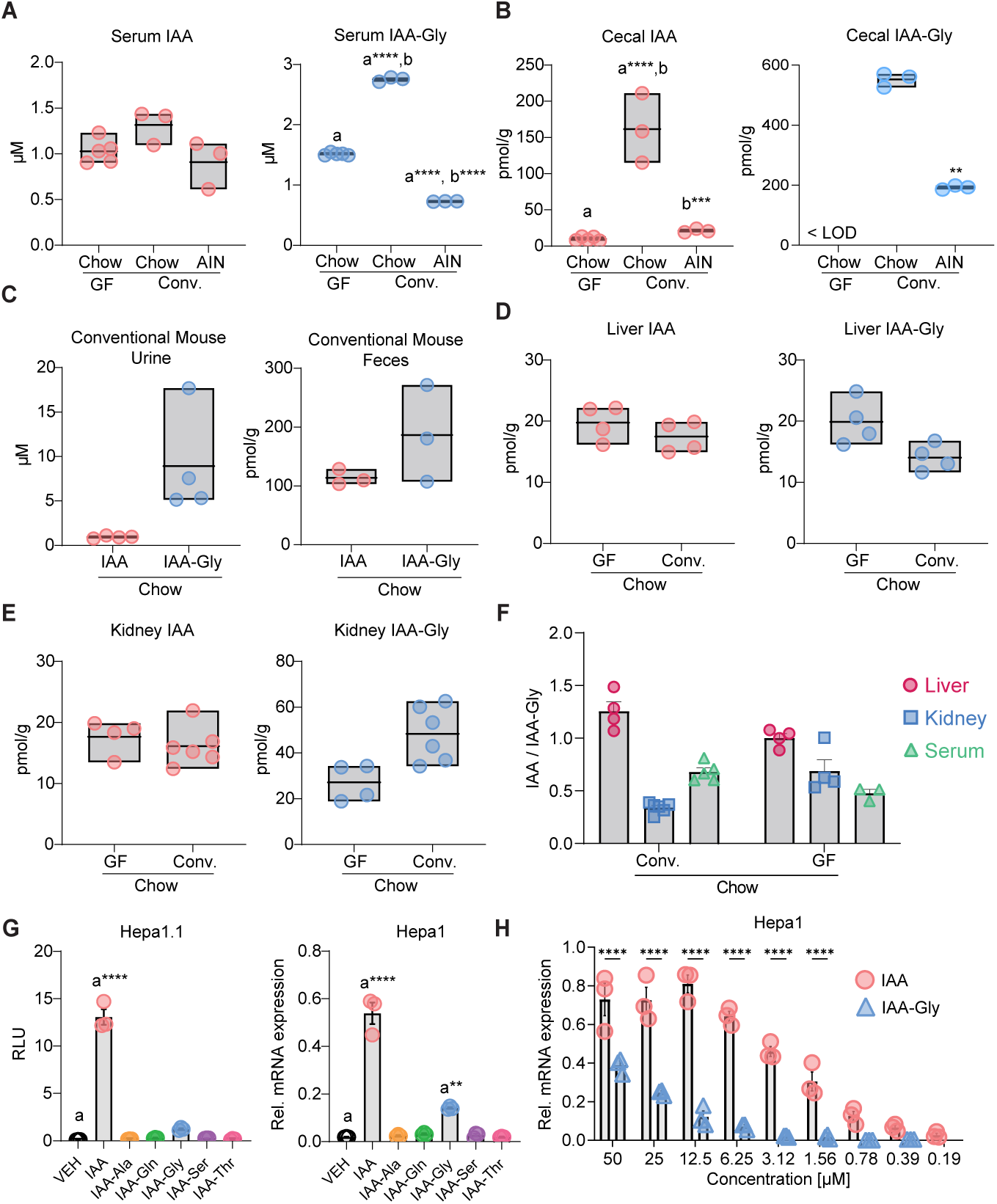
Indole-3-acetylglycine was detected in mouse bio samples and exhibits reduced AHR activation potential. **A-B.** Quantification of IAA and IAA-Gly in serum and cecal contents collected from germ-free (GF, n=5) and conventional (Conv., n=3) mice fed a standard chow diet or a phytochemical-depleted AIN-93G diet (AIN, n=3). **C.** Quantification of IAA and IAA-Gly in urine (n=3) and feces (n=3) samples from Conv. mice fed a chow diet. **D-E.** IAA and IAA-Gly concentrations in liver and kidney samples from Conv mice fed a chow diet and from GF mice fed either a chow (n = 4) or AIN diet (n=5), as measured by LC-MS. **F.** Ratio of IAA/IAA-Gly in liver, kidney and serum in Conv. and GF mice fed a chow diet. **G.** Five IAA–amino acid conjugates (20 μM) were evaluated using serum-free media for AHR activation potential by measuring luciferase activity in Hepa1.1 cells and *Cyp1a1* gene expression in Hepa1 cells (n=3). **H.** Hepa1 cells were treated with IAA and IAA-Gly at a range of concentrations (0.19-50 μM) using reduced serum media to evaluate dose-dependency (n=3). Data are represented by mean ± S.E. or box and whisker plots. Cells were exposed to treatments for 4 h. Statistical comparisons were made using a *t-*test, one-way ANOVA or two-way ANOVA with Sidak multiple comparison post-test; *: p < 0.05, **: p < 0.01, ***: p < 0.001, ****: p < 0.0001. Alphabetical characters indicate statistical comparisons between 2 groups.

These data clearly indicate that IAA and IPA can undergo glycine conjugation; therefore, we examined whether such ligands are conjugated with other amino acids. A series of IAA-X conjugates, where X represents alanine (Ala), glutamine (Gln), serine (Ser), and threonine (Thr), were synthesized for use as LC-MS standards. LC-MS analysis of cecal contents from conventional chow-fed mice failed to identify their presence and were below the limit of detection (Table S1), for all the IAA-X conjugates examined. This finding suggests that glycine is preferentially utilized as a co-substrate for IAA conjugation.

To further investigate the human metabolism of the indolic AHR ligands IAA and IPA, we examined the possibility that these ligands may generate Indole-3-carboxylic acid and/or its glycine conjugate through medium-chain acylCoA dehydrogenase mediated β-oxidation, similar to that observed with the metabolism of phenylpropionic acid^36^. LC-MS analysis from our cohort of human serum samples failed to detect these potential metabolites at the LOQ (Table S1). The presence of Indole-3-acrylic acid and indole-3-acryloylglycine were also examined in human serum and were similarly below the threshold for qualitative detection.

### Fecal IAA-Glycine/IAA conjugation and deconjugation

The presence of both IAA and its conjugate IAA-Gly in both cecal and fecal contents suggests that IAA is a substrate for reversible microbial glycine conjugation/deconjugation. To examine this concept further, fresh feces were collected from conventional chow-fed mice and incubated, under anaerobic conditions, with either indole-ring deuterated d5-IAA or d5-IAA-Gly. Fecal suspensions were sampled over time and analyzed by LC-MS for deuterated IAA or IAA-Gly. Fecal samples incubated with d5-IAA revealed the presence of conjugated d5-IAA-Gly (Fig. S1). Similarly, fecal samples incubated with d5-IAA-Gly revealed the presence of deconjugated IAA (Fig. S1). These data demonstrate the capacity of mouse fecal microbiota to reversibly conjugate and deconjugate IAA with glycine.

### Source of host IAA conjugation

Having established that the presence of IAA and its glycine conjugate in mice are not absolutely microbiota dependent, we attempted to identify the source of host IAA conjugation. Previous observations suggest that the liver and kidney are the predominant sources of acylCoA-amino acid conjugation. To investigate this hypothesis, we performed LC-MS quantification of IAA and IAA-Gly in mouse hepatic and renal tissue. Data indicates the hepatic levels of IAA and IAA-Gly to be 17-20 and 14-20 pmol/g respectively, in both germ-free and conventional chow-fed mice (Fig. 2D). Renal quantification of IAA and IAA-Gly indicate concentrations of 15-18 and 24-45 pmol/g respectively, across both groups of mice (Fig. 2E). The observed elevation in renal IAA-Gly, relative to hepatic, suggests the kidney as a likely source of IAA conjugation prior to urinary elimination (Fig. 2C). In support, the renal IAA/IAA-Gly ratio in conventional mice is 1:3 which significantly exceeds the hepatic 1:1 ratio (Fig. 2F). To directly demonstrate the capacity of host tissues to conjugate IAA, in vitro conjugation reactions were performed using both hepatic and renal mitochondrial lysates incubated with IAA and glycine. Both renal and hepatic lysates were permissive for IAA conjugation with glycine, as assessed using LC-MS analysis (Fig. S2). On an IAA-Gly/g protein basis, renal lysate conjugation was greater than hepatic conjugation, further suggesting the kidney as a dominant site of IAA conjugation.

### Mouse AHR activation potential of IAA and IAA-Gly

Previous data established IAA as an endogenous AHR ligand, however, its glycine conjugate IAA-Gly, has not previously been investigated in the context of AHR activation. Using the AHR-sensitive mouse Hepa1.1 reporter cell line we examined the activation potential of 20 μM IAA-Gly (Fig.2G). Consistent with its status as an AHR ligand, IAA elicited a significant and robust induction in AHR activity relative to the vehicle treated. Based on Individual statistical comparison with control, it appears that IAA-Gly has weak but significant activation potential in the context of mouse AHR (Fig. 2G). Although not detected in vivo, we decided to utilize our panel of IAA-X (non-glycine) conjugates to assess the structural specificity of mouse AHR activation by IAA-Gly in mouse cells. In the context of the mouse AHR, none of the IAA-X (IAA-Ala, IAA-Gln, IAA-Ser or IAA-Thr) elicited significant reporter cell line induction at 20 μM (Fig. 2G). To corroborate these observations, we assessed the AHR activation potential of IAA, IAA-Gly and IAA-X by qPCR using mouse Hepa1 cells in the context of *Cyp1a1* induction (Fig. 2G). Relative to control, IAA elicited a significantly robust induction of *Cyp1a1*. IAA-Gly also significantly induced *Cyp1a1* expression, albeit to a lesser degree than IAA. In contrast, none of the other IAA-X conjugates demonstrated significant induction at 20 μM (Fig. 2G). These data suggest that glycine conjugation of IAA retains a diminished capacity to activate mouse AHR, relative to IAA. Additionally, such activation appears to be structurally specific for glycine, since none of the IAA-X conjugates display any mouse AHR activation potential.

The establishment of IAA-Gly, at 20 μM, as an agonist for mouse AHR led us to perform a dose-response study to determine if the levels of IAA-Gly quantified in mouse serum (3 μM) are sufficient to promote physiological mouse AHR activation. Mouse Hepa1 cells were incubated with a concentration range (0.19-50 μM) of either IAA or IAA-Gly and *Cyp1a1* expression was assessed (Fig. 2H). Consistent with AHR agonist activity, both IAA and IAA-Gly elicited dose-dependent *Cyp1a1* induction with IAA exhibiting enhanced potency relative to IAA-Gly. Although dose-dependent activity is observed, significant induction by IAA-Gly was only detected above 6 μM (Fig. 2H and S3), which is above normal physiological concentrations in mouse serum.

### IAA and IAA-Gly presence in humans

The detection of IAA-Gly, as a potential AHR agonist, in mouse serum and tissues led us to question whether IAA-Gly is also present in human serum. LC-MS analysis of serum from a cohort of 40 individuals revealed the presence of both IAA and its glycine conjugate (Fig. 3A). IAA concentrations ranging from 700 nM to 3.1 μM (%CV = 33%) and a mean serum concentration of 1.62 ± 0.54 μM were detected. The levels of IAA-Gly were strikingly similar to those of IAA, with serum concentrations ranging from 600 nM to 2.6 μM (%CV = 33%) with a mean serum concentration of 1.39 ± 0.46 μM. Subsequent analysis indicates a strong (R = 1.0) positive correlation between serum levels of IAA and IAA-Gly (Table S2 and Fig. S4). In contrast, there is no correlation between serum levels of IAA and IPA (Fig. S4). LC-MS analysis of human fecal samples from an independent cohort revealed similarly variable levels of IAA in the low nmol/g range, whereas IAA-Gly levels however were below the limit of detection (Fig. 3A).

**Figure 3.**
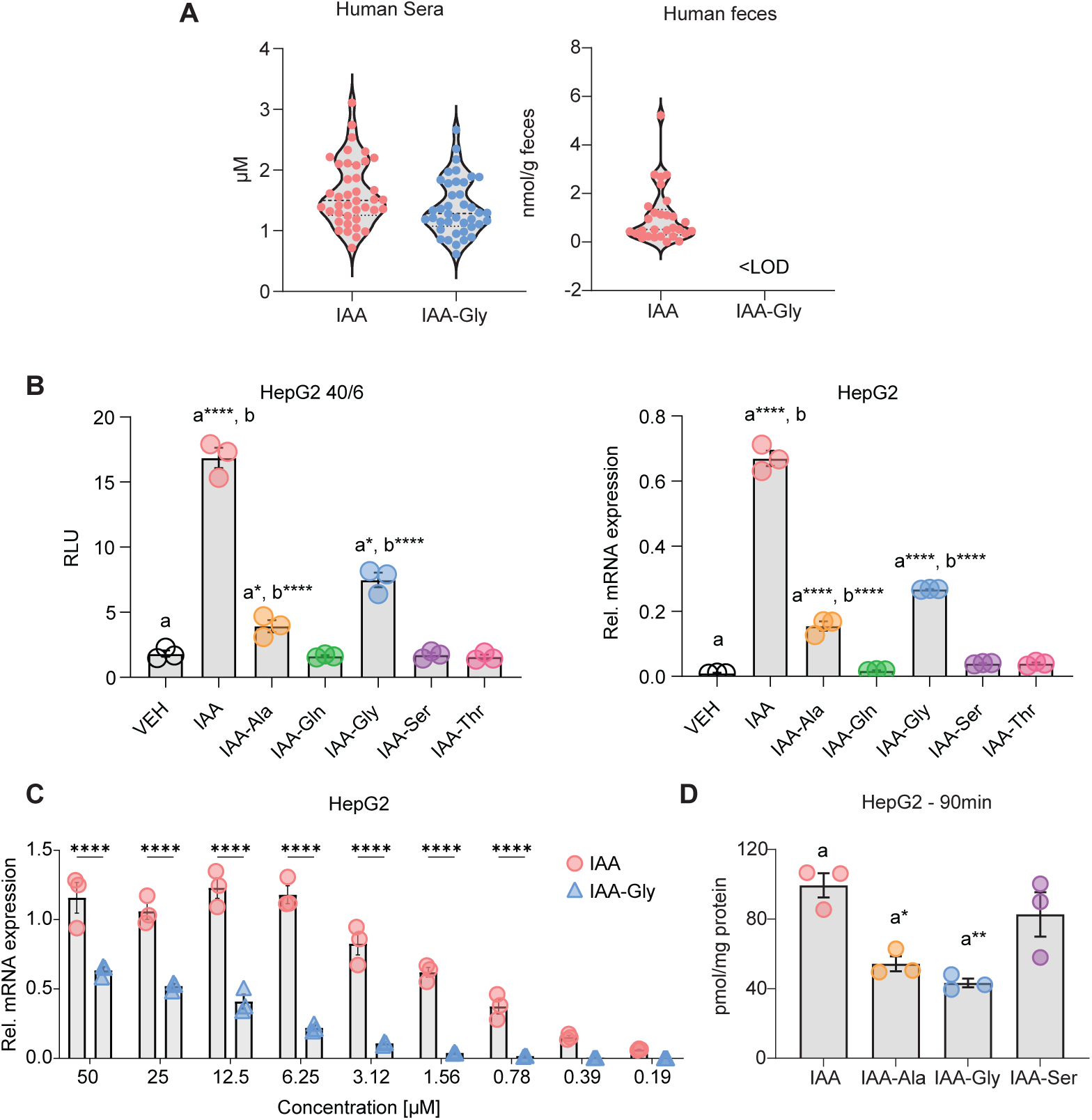
Indole-3-acetylglycine induces AHR activity. **A.** LC-MS analysis of human serum and fecal samples was used to quantify IAA and IAA-Gly. Samples were collected from individuals fed a defined diet (n=40). **B.** IAA conjugates with alanine, glutamine, glycine, serine, and threonine were evaluated for AHR activation potential at 20 μM in serum-free media by measuring luciferase activity and *CYP1A1* gene expression (n=3). **C.** *CYP1A1* gene expression was measured in response to increasing doses of IAA and IAA-Gly in reduced serum media (n=3). **D.** HepG2 cells in Hanks balanced salt solution were treated with 100 μM IAA, IAA-Ala, IAA-Gly, or IAA-Ser for 90 min. Cellular concentrations of each compound were quantified by LC-MS (n=3). Total protein was measured by UV absorbance at 280 nm and used for normalization. Data are represented by mean ± S.E. Cells were exposed to treatments for 4 h. Statistical comparisons were made using a *t-*test, one-way ANOVA or two-way ANOVA with Sidak multiple comparison post-test. *: p < 0.05, **: p < 0.01, ***: p < 0.001, ****: p < 0.0001. Alphabetical characters indicate statistical comparisons between 2 groups.

### Activation potential of IAA and IAA-Gly with human AHR

The presence of IAA-Gly in human serum at concentrations exceeding that found in mouse serum led us to explore its agonist potential, relative to IAA, in the context of human AHR. Using human HepG2(40/6) reporter cells, exposure to 20 μM IAA or IAA-Gly prompted significant induction associated with both IAA (9-fold) and IAA-Gly (4-fold), relative to vehicle (Fig. 3B). However, as with mouse serum, LC-MS analysis failed to detect the presence of the IAA-X conjugate (IAA-Ala, IAA-Gln, IAA-Ser or IAA-Thr) panel in human serum. Thus, this panel was utilized to screen for the structural dependency of IAA-Gly-mediated human AHR induction (Fig. 3B). As with mouse AHR, human AHR was insensitive to IAA-Gln, IAA-Ser and IAA-Thr, at the levels tested (Fig. 3B). However, unlike mouse, human AHR was significantly activated (2-fold) by 20 μM IAA-Ala (Fig. 3B). Corroboration of these observations was established by qPCR analysis of *CYP1A1* induction in HepG2 cells. Exposure to IAA elicited a significant 67-fold induction, while IAA-Gly demonstrated a significant but diminished (relative to IAA) 27-fold *CYP1A1* induction (Fig. 3C). Consistent with the reporter analysis, IAA-Ala elicited a significant 16-fold induction; however, *CYP1A1* expression was not influenced by the remaining IAA-X panel (Fig. 3C). Confirmation of IAA-Gly as an agonist for human AHR led us to determine if the individual levels of IAA and IAA-Gly quantified in human serum (1.3 μM) are sufficient to promote physiological human AHR activation. Human HepG2 cells were incubated with a concentration range (0.19-50 μM) of either IAA or IAA-Gly and *CYP1A1* expression was assessed by qPCR (Fig. 3C). Consistent with AHR agonist activity, both IAA and IAA-Gly elicited dose-dependent *CYP1A1* induction, with IAA exhibiting enhanced potency relative to IAA-Gly. Although dose dependent activity is observed, significant induction by IAA-Gly was detected above 3 μM (Fig. 3C and S3). The dose dependent *CYP1A1* expression was similarly assessed in human Huh7 cells (Fig. S3). As with HepG2 cells, IAA exhibited enhanced potency relative to IAA-Gly, however in Huh7 cells, significant induction by IAA-Gly was evident at 1.56 μM (Fig. S3 and S5).

To determine if the observed difference in AHR activity elicited by IAA, IAA-Gly, IAA-Ala and IAA-Ser are attributable to differential levels in cellular accumulation, metabolite uptake assays were performed. HepG2 cells were incubated for 90 min with each IAA metabolite and intracellular accumulation was quantitatively assessed by LC-MS. Data indicate similar levels of accumulation between IAA and IAA-Ser, suggesting that the lack of *CYP1A1* induction by IAA-Ser is not due to restricted uptake (Fig. 3D). Comparison of intracellular IAA and IAA-Gly levels indicate reduced accumulation of IAA-Gly (Fig. 3D). IAA-Ala exhibited an intracellular centration similar to IAA-Gly, indicating its reduced CYP1A1 inducibility is not attributable to differential uptake (Fig. 3D).

To examine whether the *CYP1A1* inducibility by IAA-Gly may be a consequence of deconjugation to the known AHR ligand IAA, similar to that established in fecal microbial cultures, HepG2 cells were incubated with 100 μM IAA-Gly for 24 h and the level of IAA was determined by LC-MS. The level of IAA, following incubation with IAA-Gly, was below the limit of detection (Table S1), indicating the absence of a deconjugation mechanism to yield IAA and subsequent IAA-mediated AHR activation (Fig. S6). Similar incubations performed with IAA demonstrate IAA-Gly to be below the limit of detection, indicating the absence of IAA conjugation with glycine (Fig. S6).

### Trp metabolites and IL1B mediated combinatorial regulation of *PAI-2*

Fibroblast-like synoviocytes are important mediators of inflammation within the joints of a rheumatoid arthritis patient. Plasminogen activator inhibitor-2 (PAI2, SERPINB2) levels correlate with inflammatory status, and its expression is regulated by cytokines and AHR activity^37,38^. To determine if IAA or IAA-Gly can modulate inflammatory gene expression, primary human synoviocytes (isolated from a rheumatoid arthritis patient) were incubated with 10 μM IAA or IAA-Gly in isolation or in combination with 0.5 ng/ml IL1B and *PAI-2* expression was assessed by qPCR. Neither IAA, IAA-Gly nor IL1B in isolation elicited a significant induction of *PAI-2* (Fig. 4A). However, combinatorial exposure to either IAA or IAA-Gly with IL1B prompted significant *PAI-2* induction (Fig. 4A). To determine if the combinatorial effect of IAA or IAA-Gly with IL1B is AHR dependent, cells were co-incubated with the established AHR antagonist GNF351. Antagonism of AHR ablated the combinatorial activity of both IAA and IAA-Gly (Fig. 4A). To expand on this observation, we examined whether IAA-Gly, in the context of a previously established pool of AHR-sensitive Trp metabolites^15^, can modulate IL1B-dependent *PAI-2* expression. Synoviocytes were treated with a Trp metabolite pool comprising human serum concentrations of IAA, IAA-Gly, IPA, ILA, IAld, KA and KYN derived from a single healthy individual (29DP1) with or without IL1B. Exposure to IL1B and the 29DP1 Trp metabolite pool synergistically induced the expression of *PAI-2* over that observed with IL1B or 29DP1 metabolites alone (Fig. 4B). Furthermore, co-incubation with the AHR antagonist GNF351 significantly suppressed *PAI-2* expression to the level of that obtained with IL1B alone (Fig. 4B). These data indicate that by independent activity assays that IAA-Gly exhibit AHR agonist at physiologically relevant concentrations.

**Figure 4.**
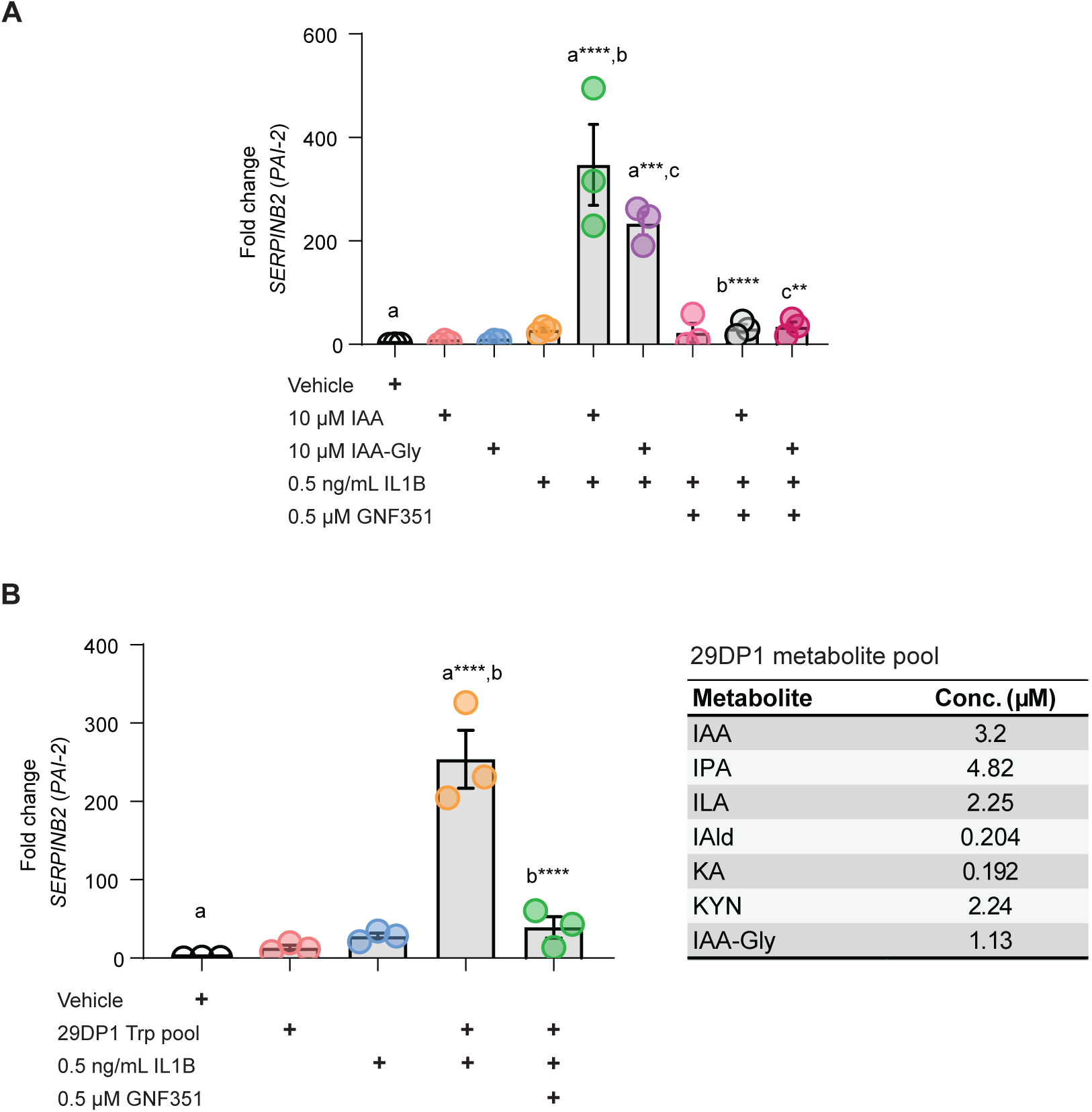
Trp metabolite activation of AHR enhances IL1B mediated *SERPINB2* (*PAI-2)* induction in human synovial fibroblasts. **A.** Human synovial fibroblasts were exposed to indicated concentrations of IAA or IAA-Gly (n=3) just prior to 6 h incubation with 0.5 ng/ml human IL1B. Where indicated, synoviocytes were pre-exposed to the AHR antagonist GNF351 (500 nM) for 30 min prior to the addition of IAA, IAA-Gly and IL1B. Subsequently, *SERPINB2* (*PAI-2*) gene expression was then analyzed by RT-qPCR. **B.** Human synovial fibroblasts were exposed to a Trp metabolite pool (29DP1) comprising IAA, IPA, ILA, IAld, KA, KYN and IAA-Gly (n=3) at concentrations previously quantified in serum from a human subject. Concentrations of individual Trp metabolites are listed in the adjoining table. Immediately following the addition of 29DP1, synoviocytes were treated with 0.5 ng/ml human IL1B for 6 h. Where indicated, synoviocytes were pre-exposed to the AHR antagonist GNF351 (500 nM) for 30 min prior to addition of 29DP1 and IL1B. *SERPINB2* (*PAI-2*) gene expression was then analyzed by RT-qPCR. Data are represented by fold induction ± SEM (n=3) as determined. Statistical significance between treatment groups is indicated and assessed utilizing one-way ANOVA with multiple comparisons. *: p < 0.05, **: p < 0.01, ***: p < 0.001, ****: p < 0.0001. Alphabetical characters indicate statistical comparisons between 2 groups.

### Insights into agonist interactions with the AHR ligand binding pocket

In this study, we performed computational docking of IAA, IAA-Gly, and IAA-Ser within the human AHR PAS-B domain, modeled from the indirubin-bound cryo-EM human AHR structure (Table S3). The docking results for IAA, IAA-Gly, and IAA-Ser, superimposed on the structural coordinates of indirubin (shown in transparent green stick) are illustrated (Fig. 5A-C). Overlaying these ligands with indirubin (transparent green; Fig. 5A–C) revealed that all three interact with key PAS-B residues (F295, I325, C333) in the primary binding site (Fα/Gβ) near the pocket’s opening. This site facilitates key hydrophobic interactions and plays a critical role in ligand recognition and receptor activation. IAA and IAA-Gly retained strong hydrophobic interactions, while IAA-Ser engaged in a distinct hydrogen bonding network deeper in the pocket (Y322, S336, Q383), suggesting an alternative binding mode. Additionally, IAA-Ser demonstrates binding interactions that extend into the secondary pocket toward the Cα helix, providing further insights into the structural dynamics of ligand-receptor interaction. PoseView-generated 2D diagrams of these same results, illustrating the nature of their key interactions in the PAS-B domain. IAA (Fig. 5D) and IAA-Gly (Fig. 5E) engage similar hydrophobic residues at the primary binding site, while IAA-Ser (Fig. 5F) shows a shift toward hydrogen bonding in the secondary site, limiting its hydrophobic interaction with F295.

**Figure 5.**
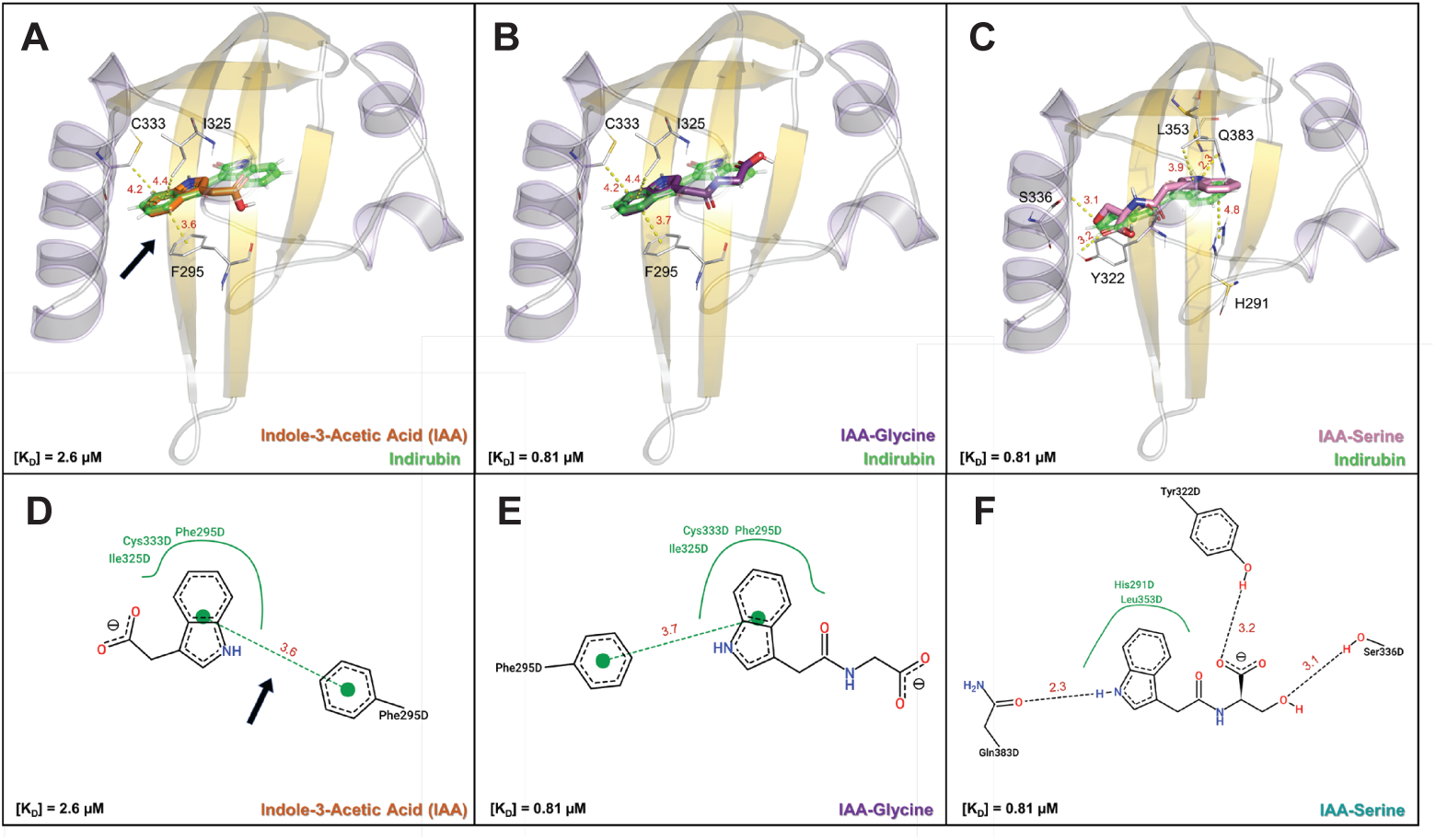
Comparative docking study of IAA and IAA-amino acid conjugates in the human AHR PAS B domain. Ligand binding properties of IAA (orange stick) and its amino acid conjugates IAA-Gly (purple stick) and IAA-Ser (pink stick) were explored with Autodock Vina using a computational model of the PAS-B domain derived from the cryo-EM structure of the human AHR (PDB: 7ZUB). **A.** IAA (orange stick; [KD] = 2.60 µM; see Supplemental Table 1) is shown docked in the ligand binding pocket of the PAS B domain, superimposed on the structural coordinates of its high affinity ligand, indirubin (shown as a transparent green stick; [KD] = 0.44 nM). Important active site residues (F295, I325 and C333) are shown. Important hydrophobic and electrostatic ligand contact distances to key amino acids are shown (in angstroms; Å; yellow dash). The black arrow indicates the key hydrophobic contact between the indole ring and highly conserved active site residue Phe-295 (F295). **B.** IAA-Gly (purple stick; [KD] = 0.81 µM) and **C.** IAA-Ser (pink stick; [KD] = 0.81 µM) dock the PAS-B domain with nearly identical affinity and are also shown superimposed on the structural coordinates of the higher affinity ligand, indirubin. Despite having similar maximum binding energies (-8.3 kcal/mol; see Supplemental Table 1), IAA-Gly and IAA-Ser dock the PAS-B domain in distinct conformations, with IAA-Gly forming more contacts with conserved hydrophobic residues (F295, I325, C333) **D.** The ligand-protein interactions for IAA were also visualized using PoseView-generated two-dimensional diagrams, where key binding interactions are highlighted with dashed lines for directed bonds and spline sections for hydrophobic contacts. Here, the black arrow highlights the same key interaction between the ligand and F295, as shown in Figure 5A, for reference. **E.** The 2D PoseView diagram for IAA-Gly is nearly identical to that of IAA, highlighting a highly conserved set of hydrophobic interactions between these two ligands near the putative substrate entry site. **F.** In contrast, for IAA-Ser, the 2D PoseView diagram highlights a distinct, high affinity binding configuration that is dominated by hydrogen bonding (with residues Y322, S336, and Q383) and unique hydrophobic interactions with L353 and H291, rather than F295, I325 and C333.

Analysis of both ligand planarity and the superposition of indole structures relative to indirubin, highlight these as key determinants of AHR activation (Fig. 6). Single docking poses for IAA, IAA-Gly, and IAA-Ser, superimposed with indirubin from a side-view highlights their planar orientation with respect to F295 (Fig. 6A-C). Both IAA and IAA-Gly adopt planar configurations when bound at high affinity, aligning well with indirubin, and making similar contacts with key residues like F295, I325 and C333. In contrast, IAA-Ser only binds the receptor in a planar orientation at lower affinity (K_D_=1.13 μM; Fig. 6C), exhibiting a kinked conformation when bound at high affinity (K_D_=0.81μM; Fig. 4C and Fig. S1C). Figure 6D-F expands this analysis by highlighting the superposition of the indole of IAA and its conjugates (IAA-Gly and IAA-Ser) onto the bi-indole structure of indirubin. The indole of IAA and IAA-Gly superimpose closely with indirubin’s structure in the primary binding pocket (Fig. 6D and 5E). In contrast, the indole ring of IAA-Ser fails to achieve the prototypical alignment in a planar configuration, suggesting that even ligands with significant binding affinity may have reduced transactivation potential if misaligned within the primary or secondary binding pocket, as compared to indirubin (Fig. 6F). Overlays of multiple docking poses for each ligand, confirm their consistent engagement with both the primary and secondary regions of the pocket (Fig. S7). IAA docking poses maintain consistent planarity across binding energies, whereas IAA-Gly and IAA-Ser exhibit greater variability (Fig. S7A-C). Notably, IAA-Ser consistently adopts kinked conformations at relatively high affinities, which likely explains its diminished transactivation potential compared to IAA and IAA-Gly. Figures S7D-F further highlight the differences in ligand binding among the three AHR agonists by comparing their superposition on indirubin’s bi-indole structure. Three relatively high affinity docking poses of IAA show that, while IAA consistently docks in a planar configuration, only the highest affinity pose replicates the prototypical indole ring positioning relative to F295 (Fig. S7D). IAA-Gly binds in two prototypical orientations, with its indole ring closely superimposing on indirubin’s bi-indole structure for both high and moderate affinity results (Fig. S7E). In contrast, IAA-Ser adopts a kinked conformation at high affinity and only displays a planar configuration at lower affinities. (Fig. S7F) However, none of the observed planar configurations of IAA-Ser properly position the indole ring relative to indirubin’s bi-indole structure.

**Figure 6.**
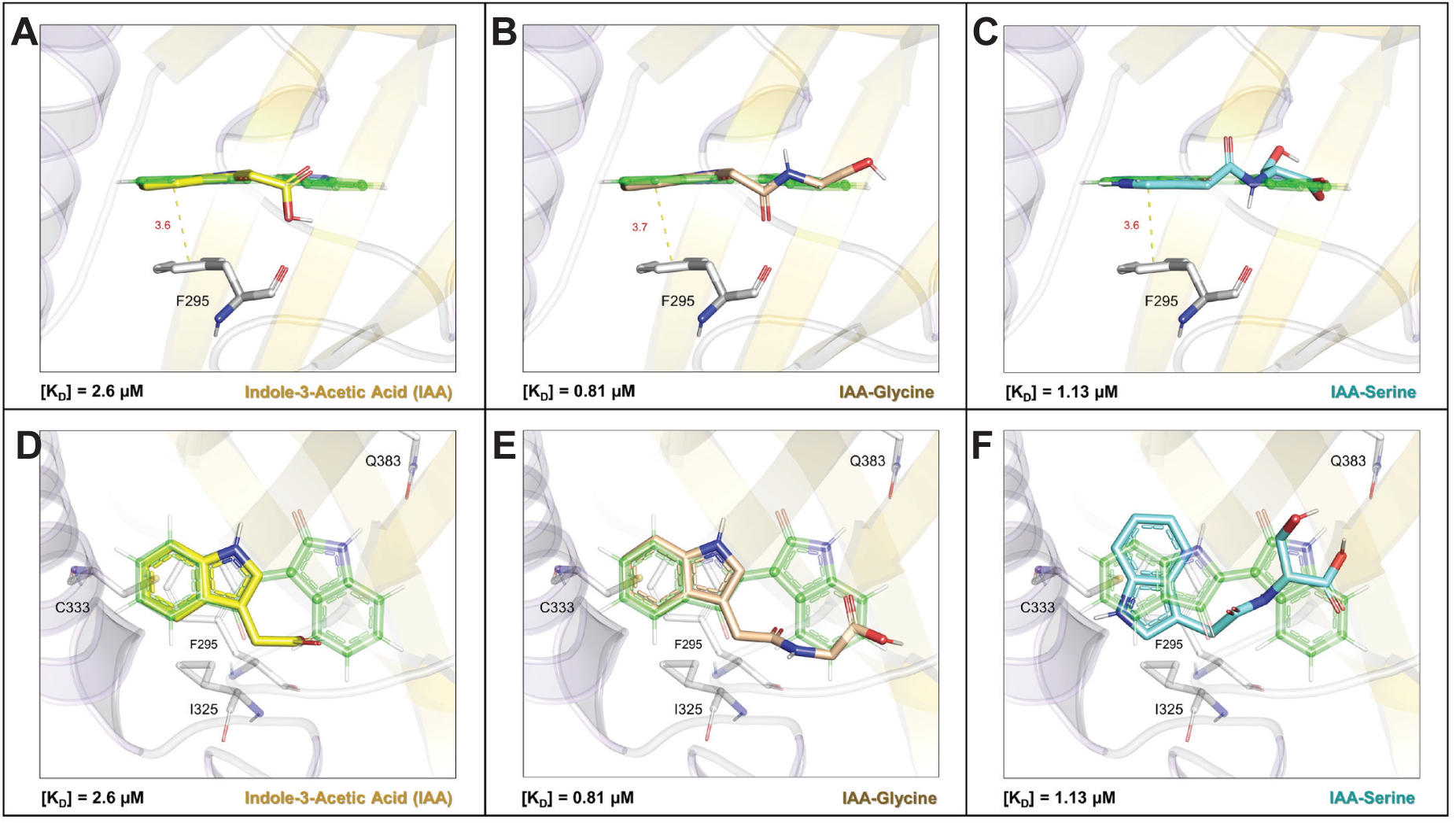
Distinguishing the structural basis for AHR agonism through planarity comparisons with indirubin in the PAS B domain. Ligand binding properties of IAA, IAA-Gly and IAA-Ser were further explored for correlations with experimental, AHR activation data generated *in vitro.* **A.** The highest affinity binding conformation for IAA (yellow stick; [KD] = 2.60 µM; see Supplemental Table 1) is shown superimposed on the structural coordinates of indirubin (transparent green stick) to highlight their similar planarity with respect to F295, in the center of the PAS B domain. **B.** IAA-Gly (tan stick; [KD] = 0.81 µM) displays similar planarity as IAA, when its highest affinity configuration is superimposed on indirubin. **C.** In contrast, IAA-Ser displays planar binding with respect to indirubin only at moderate affinity (cyan stick; [KD] = 1.13 µM) but adopts a more kinked configuration at higher affinity ([KD] = 0.81 µM; as shown in Figure 5C), suggesting a less flexible binding mode for the larger conjugate. Ligand binding properties of IAA, IAA-Gly and IAA-Ser were further assessed in relation to their superposition on the bi-indole structure of indirubin. **D.** The high affinity docking result for IAA (yellow stick; [KD] = 2.60 µM) closely aligns with the structural coordinates of one of the indole rings of indirubin (transparent green stick), specifically the ring located near the opening of the ligand binding pocket at the Fα/Gβ site. This alignment facilitates an identical hydrophobic contact with the key residue F295, central to the PAS-B domain. **E.** High affinity docking results for IAA-Gly (tan stick; [KD] = 0.81 µM) also displayed a near identical superposition with the indole ring of indirubin, proximal to the primary (Fα/Gβ) binding site. **F.** In contrast, IAA-Ser, which displays planar binding with respect to indirubin only at moderate affinity (cyan stick; [KD] = 1.13 µM; see Figure 2c), does not dock with its indole ring in the same configuration as IAA, IAA-Gly or indirubin, suggesting a distinct set of contacts in the active site that may limit transactivation of the receptor.

### Species comparison of IAA- and IAA-Gly-mediated AHR activation

If Trp metabolites are important endogenous mediators of the developmental and homeostatic biological activity facilitated by the AHR then their activation potential should be similar across mammalian species. To address this hypothesis, we compared the activation potential of two high-affinity polycyclic aromatic AHR agonists with endogenous bicyclic Trp metabolites in human and mouse hepatoma established cell lines. Dose-response studies indicated that TCDD activation potency, as determined by the EC_50_, was ∼22-fold lower in human HepG2 cells compared with mouse Hepa1 cells (Fig. 7A)—consistent with previous reports that TCDD has 10-fold lower affinity for the human AHR^39,40^. Next, we tested the endogenous, high-affinity ligand 6-formylindolo[3,2b]carbazole (FICZ), which similarly revealed an ∼8-fold decrease in human HepG2 EC_50_ compared to mouse Hepa1 cells (Fig. 7B), thus consistent with the species-dependent data obtained with TCDD. In contrast, IAA, IAA-Gly, and IAA-Ser each exhibited a 1.2-1.9-fold more potent activation potential in human HepG2 cells compared with mouse Hepa1 cells (Fig. 7C-E) while ILA exhibited a similar AHR agonist potency in both cell lines (Fig. 7F). Taken together, these data suggest IAA and IAA-Gly exhibited a modest increase in potency in the human cell line, the opposite of what was observed with TCDD and FICZ (Table 1).

**Figure 7.**
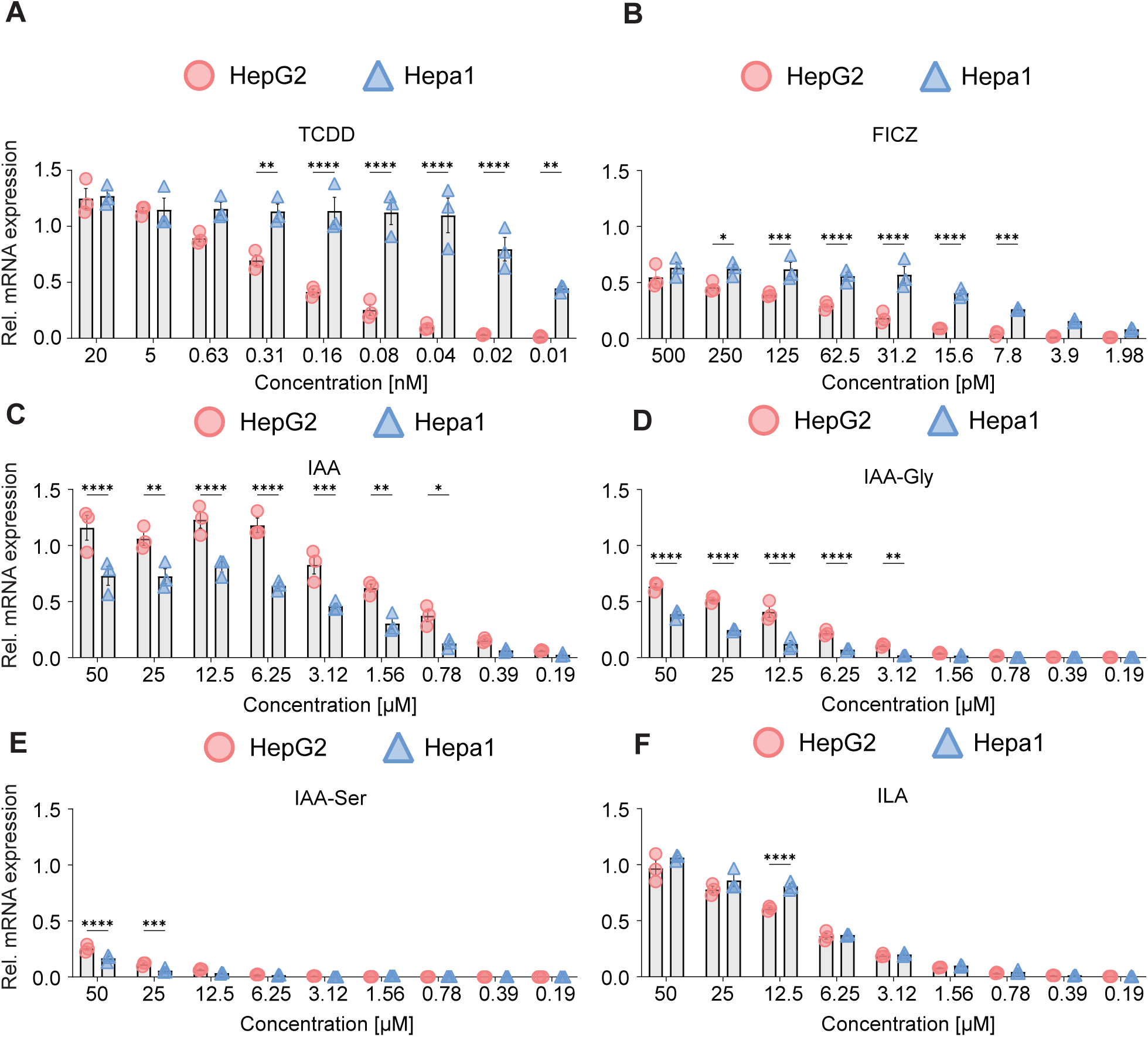
Species comparison of AHR mediated *CYP1A1* gene expression in HepG2 and Hepa1 cells after exposure to target compounds. Cells were treated for 4h with either VEH, TCDD (**A**), FICZ (**B**), IAA (**C**), IAA-Gly (**D**), IAA-Ser (**E**), or ILA (**F**) at the doses indicated. qPCR analysis for relative *CYP1A1* mRNA expression was performed. Data were normalized to the human or mouse housekeeping gene beta-actin, and the vehicle baseline was subtracted. Data are represented by mean ± SEM, n=3, and statistical analysis was performed using two-way ANOVA analysis followed by Sidak multiple-comparison test. *: p < 0.05, **: p < 0.01, ***: p < 0.001, ****: p < 0.0001.

**Table 1.** Comparative AHR ligand EC_50_ between human and mouse.

| LIGAND | HUMAN AHR EC<sub>50</sub> | MOUSE AHR EC<sub>50</sub> | ΔEC<sub>50</sub> |
| --- | --- | --- | --- |
| TCDD | 286 pM | 12.9 pM | ~ 22-fold > mouse AHR |
| FICZ | 77 pM | 10 pM | ~ 8-fold > mouse AHR |
| IAA | 1.3 μM | 1.9 μM | ~ 1.5-fold > human AHR |
| IAA-GLY | 8.9 μM | 17.3 μM | ~ 1.9-fold > human AHR |
| IAA-SER | 23.5 μM | 27.2 μM | ~1.2-fold > human AHR |
| ILA | 8.6 μM | 7.8 μM | ~ 1.1-fold > mouse AHR |

### Indole-3-carboxylates exhibit antioxidant potential

In addition to their capacity to induce AHR transcriptional activity, both IPA and IAA have previously been shown to exhibit significant antioxidant potential. We tested whether IAA, IPA, and IAA-Gly at typical concentrations in human serum, or IPA-Gly levels found in normal mouse serum can individually or collectively mediate significant antioxidant activity. Results revealed that, at physiologically relevant concentrations, only IPA exhibits significant antioxidant activity (Fig. 8). However, IAA-Gly/IAA and IPA/IAA-Gly/IAA in these combinations at concentrations common in human serum, exhibited significant activity. Thus, we conclude that mixtures of Trp metabolites exhibit enhanced antioxidant potential.

**Figure 8.**
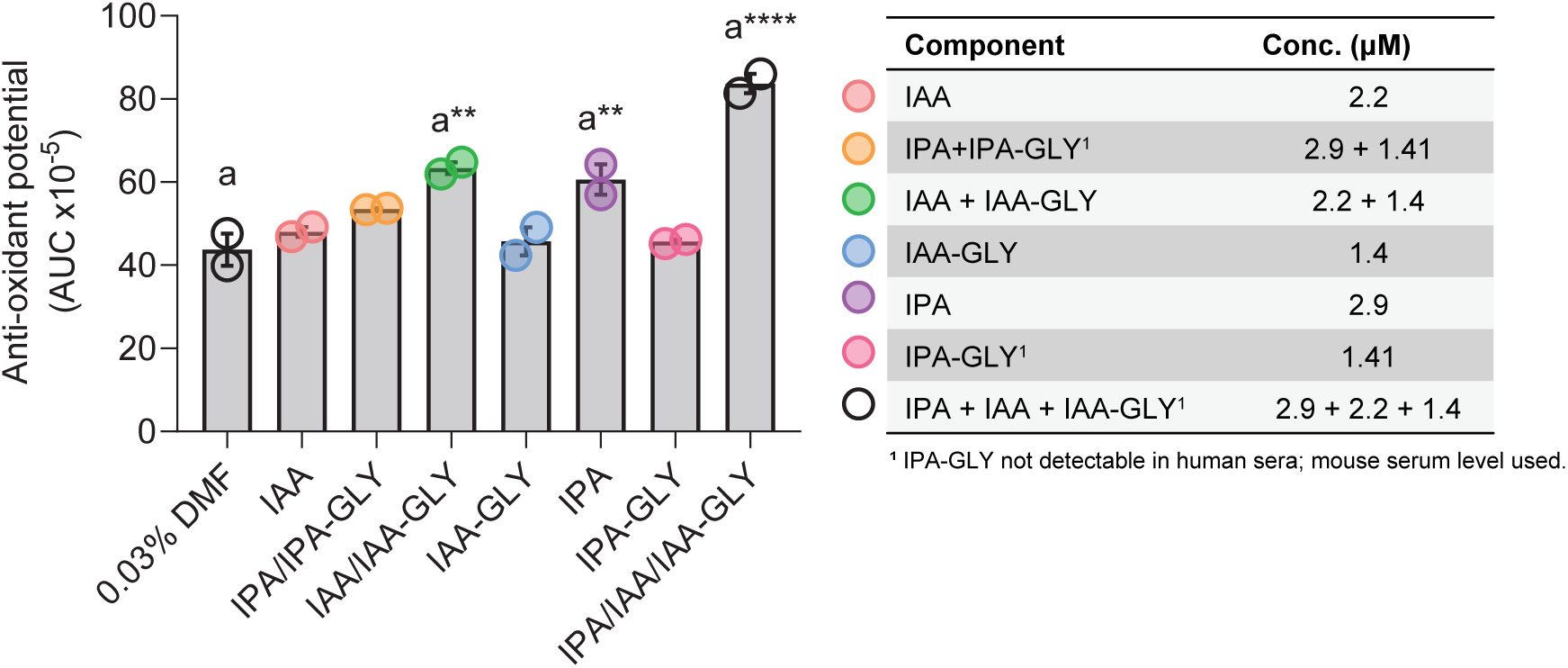
Intrinsic chemical antioxidant potential of IPA, IAA and their respective glycine conjugates. Chemical antioxidant potential of IPA, IAA, IAA-GLY, IPA-GLY and combinations were quantified, using previously determined serum concentrations, as indicated. Antioxidant assays were performed using the ORAC (AOX-2) antioxidant kit at a final DMF concentration of 0.03% (v/v). Data represents mean kinetic fluorescence AUC (arbitrary units) × 10^−5^ ± SEM, (n=2). Statistical analyses were performed using one-way ANOVA with multiple comparisons. *: p < 0.05, **: p < 0.01, ***: p < 0.001, ****: p < 0.0001. Alphabetical characters indicate statistical comparisons between 2 groups.

## Discussion

The AHR plays important roles in a myriad of physiological processes, such as embryogenesis, reproductive success, and immune homeostasis. Despite these diverse functions, the mechanisms underlying homeostatic or basal AHR activation are only beginning to be understood. Recently, we determined that a number of serum Trp metabolites are host-derived and circulate at concentrations sufficient to activate the AHR, even in fasted humans^15^. Importantly, these Trp metabolites are not CYP1A1 substrates and could maintain elevated or at least CYP1A1 constitutive levels without triggering a negative feedback loop^41^. Thus, the maintenance of significant basal CYP1A1 enzymatic activity would be primed to metabolize multi-ring exogenous substrates. Tryptophan is metabolized by the microbiota in the gastrointestinal tract to a number of metabolites including IPA and IAA, and each has been detected at μM levels in human serum^15^. The host is also capable of metabolizing tryptophan through amino acid transaminase or deaminase activity to indole-3-pyruvic acid, which undergoes further metabolism additional metabolites, likely including IAA^27^. The liver contains relatively high concentrations of amino acid transaminases, such as GOT1 and tyrosine aminotransferase (TAT), and the spleen expresses relatively high levels of the amino oxidase IL4I1, suggesting that these organs are major sources of indole-3-pyruvic acid production and consequently, IAA and ILA.

The fate of many polar endogenous or exogenous anions is either through excretion of the parent compound or direct conjugation reactions. Glycine conjugation is a common two-step enzymatic reaction for a variety of carboxylic acids. The first step is activation to an acyl-CoA intermediate by mitochondrial xenobiotic/medium-chain fatty acid CoA ligases (ACSM2A/B in humans or ACSM2 in mice). The expression pattern of ACSM2 in mice is found only in the proximal tubular cells of the kidney, while in humans ACSM2A/B is expressed in both liver and kidney^42,43^. The second step is conjugation with glycine, mediated by GLYAT. Examples of substrates include benzoate, salicylate, and medium-chain fatty acids^44^. GLYAT is expressed in the liver and kidney of both mice and humans. Results presented here indicate that IPA and IAA are conjugated with glycine by both the host and gut microbiome. In mice, glycine conjugation most likely occurs in the kidney, as the enzymes necessary for glycine conjugation are only present in the kidney. In addition, the concentration of IAA and IAA-Gly are higher in the kidney and urine than in serum or liver, further suggesting kidney specific glycine conjugation. Thus, the presence of IAA/IPA-glycine conjugates in mouse serum is most likely due to reabsorption from urine by proximal tubules back into the bloodstream, perhaps similar to a mechanism observed with dipeptides^45^. Glycine conjugation may serve as a mechanism to enhance clearance by the kidney. Support for this concept can be found in the expression of human genetic variants of ACSM2A that exhibit reduced conjugation potential, which has been associated with a 4.7-fold increase in plasma levels of IPA, suggesting that IPA conjugation leads to enhanced clearance^35,45^. Indeed, the data presented here reveal that most of the IAA present in urine is conjugated with glycine, further supporting this explanation.

IAA has been identified as a significant driver of human AHR activity at concentrations found in human serum, yet the metabolic fate of this compound has not been previously reported. The observation presented here, that IAA-Gly exhibits significant AHR agonist activity, was somewhat unanticipated, considering that AHR ligands are assumed to often lose activation potential following phase II metabolism^46^. Given that this generalized assumption does not hold true for IAA-Gly, this metabolite should also be assessed alongside IAA to determine their combined effect on AHR activation. In our previous assessment of human AHR activation potential of relatively abundant serum Trp metabolites, three ligands exhibited AHR activation potential at typical concentrations present: IAA, KYN, and KA, with the latter two in combination exhibiting the greatest AHR impact^15^. However, we have now reassessed this observation by considering the activation potential of IAA-Gly in combination with IAA. Combining IAA and IAA-Gly levels in human serum results in a total concentration of 3 μM. These results underscore the importance of products made by amino acid transaminases, such as GOT1 and TAT, to basal AHR receptor activation potential.

The AHR can regulate expression of several cytokine/chemokine (e.g., *IL6*, *CCL20*) and growth factor (e.g., *AREG*, *EREG*) genes through cytokine and AHR ligand combinatorial exposure^47–49^. The mechanism of IL1B and AHR agonist synergistic induction of IL6 has been determined^50^. There are several noncanonical dioxin-responsive elements at ∼3000 bp upstream within the *IL6* promoter that recruit the AHR/ARNT complex upon ligand treatment. IL1B leads to the recruitment of NFkB (p65/p50) to the proximal *IL6* promoter. The presence of the AHR/ARNT mediates the displacement of HDAC1 and the subsequent enhanced acetylation of p65, which facilitates to an increase in p65 mediated *IL6* gene expression. SERPINB2 is a highly pleotropic factor that exhibits diverse functions often linked to chronic diseases, such as arthritis, through a role in inflammatory responses^51^. SERPINB2 expression can be mediated independently by Ah receptor activation and inflammatory signals^52,53^. The mechanism of AHR activation of *SERPINB2* expression appears to be distinct from that observed with *IL6* and involves a pause-release and elongation mechanism^54,55^. In this report, IAA, IAA-Gly and a mixture of Trp metabolites in combination with IL1B induces *SERPINB2* expression in primary synovial fibroblasts. Illustrating the potential impact of physiological relevant circulating Trp metabolites potential to modulate chronic disease progression through AHR activation.

Docking studies determine the ability of a given compound to dock into a ligand binding pocket, which does not always correlate with AHR-mediated transcriptional activity. This discrepancy arises from the inherent static nature of docking models, which fail to capture the dynamic conformational transitions required for AHR activation, nuclear transport, HSP90 dissociation, and heterodimerization with ARNT. High-affinity antagonists exemplify this limitation, as they bind the receptor tightly yet fail to invoke transcriptional activity. Because the structural intermediates coupling ligand binding to transcriptional output remain incompletely resolved, docking results alone have limited predictive value.

However, docking studies can provide mechanistic insight when comparing weak versus potent agonists, especially among structurally related compounds. They can also offer insight into why minor changes in a series of closely related compounds can markedly affect agonist activity. For example, one recent study examined three structurally related polycyclic aromatic hydrocarbons that differed by ∼1,000-fold in AHR activation potential, yet their docking scores varied only modestly^56^. This finding indicates that transcriptional efficacy depends more on ligand orientation and the conformational changes it induces than on raw binding energy. Thus, the precise geometry of ligand positioning within the pocket, rather than the magnitude of the docking score, appears to determine productive receptor activation dynamics.

In our studies, we utilized the human AHR cryo-EM structure, which includes indirubin in the ligand binding pocket^4^. Indirubin is a potent human AHR ligand that essentially consists of two oxindole residues linked together. Docking results show that IAA and IAA-Gly align one indole ring similarly to indirubin’s proximal indole, while their aliphatic groups occupy the space normally filled by indirubin’s second ring. These results are consistent with the biological AHR agonist activity exhibited by IAA and IAA-Gly. The addition of progressively larger side chains relative to glycine results in decreased agonist activity, supporting the hypothesis that the AHR binding pocket favors planar substituents. To mechanistically understand these results, we compared the docking pose of IAA-Gly to that of IAA-Ser. Unlike IAA-Gly, the high-affinity binding pose of IAA-Ser does not align with the position or planarity of indirubin. Deviation of IAA-Ser from the prototypical binding mode, despite its significant affinity, leads to diminished receptor activation, highlighting the importance of precise structural alignment and interactions within the binding pocket for effective AHR activation. These findings underscore that the AHR has highly evolved to be activated by planar, aromatic ligands, particularly those with indole-based structures, revealing a finely tuned regulatory mechanism that selectively responds to ligands adopting the optimal planar configuration and position for productive agonist engagement. Furthermore, considering that many of these Trp metabolites (e.g., indole, indole-3-carboxylic acid) are more potent activators of the human versus mouse AHR, their impact on human-specific AHR signaling may be underappreciated and thus needs to be further explored.

There are several studies that have revealed that IPA exhibits potent antioxidant activity, attenuating neurological toxicity and positively correlating with beneficial health outcomes^30,57^. Similarly, the potential for IAA to mediate antioxidant activity has been observed but warrants further investigation^58^. Here, we demonstrate, at concentrations detectable in human serum, the combination of IAA and IAA-Gly significantly attenuates hydroxyl radical (•OH) dependent oxidation. Such antioxidant activity is further enhanced in the presence of IPA. These observations highlight the potential for the physiological milieu of tryptophan-derived indolic metabolites to suppress prooxidant activity associated with chronic diseases. Importantly, quantification of Trp metabolites in fasted human serum, from a controlled diet study, revealed a >3-fold inter-subject variation in the levels of IAA and IAA-Gly, suggesting that non-dietary factors impact circulating levels of these host-generated Trp metabolites. Serum IPA exhibited an 18-fold inter-subject variability, consistent with it being a microbially exclusive Trp metabolite and thus dependent upon differences between inter-subject microbiome composition and metabolic activity. Furthermore, diet directly influences microbial generation of IPA and likely impacts circulating levels^57^.These observations suggest that the antioxidant potential of circulating Trp metabolites are combinatorially dictated by host genetics, diet and the microbial community structure.

IAA is a broadly conserved metabolite across all domains of life, including plants, bacteria, and mammals, and may function as a mediator of inter- and intra-species communication. Indeed, research has demonstrated that bacteria in soil can signal to plants through production of IAA and alter plant growth^59^. IAA is also considered a microbial quorum-sensing molecule^60,61^. Research presented here suggests the ability of bacterially produced IAA in the gastrointestinal tract to signal to the host through activation of the AHR. In plants, IAA is an auxin that regulates plant growth, and the level of free, bioactive IAA is controlled through amino acid conjugation and storage. Glycine conjugation in animals is likely a mechanism to enhance transport into urine by the kidney, perhaps through recognition by a transporter(s) that remains to be identified or increased aqueous solubility. Notably, approximately two-thirds of IAA in mouse cecal contents is conjugated to glycine; however, the functional basis for this microbial conjugation remains unknown. Glycine conjugation may influence the activity of IAA or IPA within the gut, possibly serving as an inhibitor of microbial growth. For example, IPA has been shown to inhibit anthranilate synthase TrpE, leading to attenuated tryptophan synthesis in *Mycobacterium tuberculosis*; therefore, it is plausible that IAA may operate through a comparable mechanism in the gut^62^.

Examination of the evolution of the AHR between rodents and primates in terms of ligand activation potential may yield important insights into the identity of physiologically relevant endogenous AHR activators. In mice, residue 375 occupies a central position in the ligand binding pocket and partially determines the receptor’s affinity for TCDD and other polycyclic aromatic hydrocarbons. In high-affinity mouse alleles, an alanine residue is present, while low-affinity alleles encode a valine, in the human AHR this residue is 381^1^. Compared to other primates, including Neanderthals, only the modern human AHR carries a A381V substitution^63^. This universally conserved valine substitution in human AHR leads to a dramatic 140 to 5000-fold decrease in the *CYP1A1* induction potential of xenobiotic planar polycyclic aromatic hydrocarbons, such as benzo(a)pyrene, benzanthracene and 2,3,7,8-tetrachlorodibenzofuran. In contrast, the valine substitution has no effect on the *CYP1A1* induction potential by physiologically detectable AHR ligands, such as indirubin, indoxyl sulfate and indole. However, this study only examined primate AHR activation. Other studies have demonstrated that bicyclic endogenous compounds, such as KA and 2,8-quinolinediol, are more potent activators of the human AHR compared to the mouse AHR, using hepatoma cell lines^21,64^. The results presented here are consistent with these previous studies revealing that TCDD and FICZ exhibit 8-22-fold diminished AHR activation potential in human hepatoma cells relative to a mouse hepatoma cell line. In contrast, IAA and IAA-Gly exhibited greater human AHR activation potential, while ILA activated the mouse and human AHR to a similar extent. Such inter-species conservation of AHR activation potential by endogenous Trp metabolites, which is not apparent with xenobiotic AHR ligands, supports their role as important mediators of AHR dependent physiological processes. Conservation of endogenously mediated, but not xenoestrogen-dependent, activation is similarly reported for the estrogen receptor^65^. FICZ, despite being a tryptophan derivative, has high AHR potency relative to other Trp metabolites. However, the AHR activation potential of FICZ is species dependent and does not adhere to the paradigm of endogenous conservation—rendering it atypical. FICZ, like other tryptophan condensation products (e.g. β-carbolines), are formed non-enzymatically and, while they may be detectable in biological samples, their physiological contribution to AHR activation is ill-defined^66^.

In a previous report we examined the fasted human serum levels of IAA, IPA, indole-3-lactic acid, indole-3-carboxaldehyde, KYN and KA in a cohort of 40 individuals that were placed on a standardized isocaloric American diet^15^. These same samples were also analyzed for IAA-Gly content in this study and all of these metabolites measured vary from 4 to 25-fold. This suggests that tryptophan metabolism through the TDO/IDO and amino acid transaminase pathways exhibit considerable variability between individuals, which may be due to underlying genetics, exposure histories, health status, age, sex, etc. In addition, since IPA is produced exclusively by the gut microbiome the wide variation in IPA levels observed likely reflects difference in individual gut microbiomes. Most importantly, this would suggest that systemic AHR activation would likely vary widely, influencing both protective and detrimental effects of AHR activity. However, the significance of these observations would require extensive population-based studies to determine the importance of tryptophan metabolism and AHR activity to overall health consequences.

Despite decades of research into the ability of the AHR to bind host- and microbe-derived Trp metabolites, much remains unknown. Classical models suggest that even minor modifications to AHR ligands—such as hydroxylation, as seen in indolo[3,2-b]carbazole or 5-hydroxyindoleacetic acid—greatly reduce activity^18,46^. However, the results presented here challenge that paradigm, revealing that certain modifications, like glycine conjugation of IAA, can significantly retain AHR activation potential. The stark differences in activation potency between IAA, IPA, and their respective glycine conjugates underscore the complexity of AHR ligand recognition and point to gaps in our understanding of its structure-function relationship. Furthermore, microbial conjugation of indole compounds may represent a sophisticated strategy for modulating host signaling or microbial ecology within the gut. Whether this reflects a mechanism for immune modulation, inter-microbial competition, or another survival-related function remains an open question. These findings highlight the need for deeper investigation into how structural nuances in microbial and host metabolites influence AHR activation and, ultimately, host-microbe communication.

## Methods

### Chemicals

2-(1H-indol-3-yl)acetamido]acetic acid (indole-3-acetic acid, IAA) and 3-(1H-indol-3-yl)propanoic acid (indole-3-propionic acid, IPA) were purchased from Alfa Aesar (Heysham, Lancs. UK). d5-indole-3-acetic acid and indole-3-acrylic acid were purchased from Cayman Chemical Co., Ann Arbor, MI, USA). Indole-3-acryloylglycine was purchased from GOLDBIO (St. Louis, MO).

2-[2-(1H-indol-3-yl)acetamido]acetic acid (IAA-glycine), d5-2-[2-(1H-indol-3-yl)acetamido]acetic acid (IAA-glycine), 2-[2-(1H-indol-3-yl)acetamido]propanoic acid (IAA-alanine),), 3-hydroxy-2-[2--(1H-indol-3-yl)acetamido]propanoic acid (IAA-serine), 3-hydroxy-2-[2-(1H-indol-3-yl)acetamido]butanoic acid (IAA-threonine), 4-carbamoyl-2-[2-(1H-indol-3-yl)acetamido]butanoic acid (IAA-glutamine), 2-(1H-indole-2-carbonylamino)acetic acid (ICA-Glycine), 2-[3-(1H-indol-3-yl)propanamido]acetic acid (IPA-glycine) were synthesized as described in the supplement.

### Human Serum Samples

Human serum samples were obtained from a controlled feeding study^67^. Participants were fed a defined isocaloric average American diet. After 4 weeks, serum was collected following a 12 h fast. Forty of the individual serum samples stored at -80°C were chosen at random for this study and subjected to LC-MS analysis. The protocol was approved by the Institutional Review Board at Pennsylvania State University. Previously the concentrations of Indole-3-acetic acid, KA, indole-3-carboxaldehyde, indole-3-propionic acid, indole-3-lactic acid, and KYN in the individual human serum samples was published, including individual 29DP1^68^.

### Human fecal samples

A randomly selected collection of 29 human stool samples was obtained from BioIVT LLC (Hicksville, NY). All samples obtained were stored at −80°C.

### Sample collection from conventional and germ-free mice

C57BL/6J mice were obtained from Jackson Laboratories (Bar Harbor, ME) and bred in-house. Mice were maintained in autoclaved polypropylene cages with corncob bedding and a paper tube enrichment in a pathogen-free environment under a standard 12 h light/dark cycle. Mice had ad libitum access to PicoLab rodent diet 20 (LabDiet) and water. In certain experiments, mice were switched for 7 days to an AIN-93G (AIN) diet (Dyets, Bethleham, PA, USA). All experiments were carried out in accordance with approval from the Animal Care and Use Committee of the Pennsylvania State University. Female mice in the 8- to 12-week-old range were utilized in all experiments and were humanely euthanized by CO_2_-asphyxiation. Germ-free (GF) C57BL/6J mouse experiments were conducted at the AAALAC-accredited Animal Resource Center at Montana State University and at Pennsylvania State University. GF mice were reared in standard, sterile (autoclaved cages containing sterile (autoclaved) bedding inside a hermetically sealed isolator with HEPA-filtered airflow and maintained on sterile (autoclaved) water and sterile (autoclaved) food (LabDiet^®^ 5013, Land O’Lakes) ad libitum. Germ free surveillance was performed as previously described.^15^ Whole blood was collected by cardiac puncture using a 25 G needle and transferred to Microtainer^®^ SST^™^ blood collection tubes (Becton Dickinson, NJ). Tubes were incubated at room temperature for 30 min and then centrifuged (1,200 x *g*, min, 4°C). Serum supernatants were transferred to 2 ml O-ring screw cap tubes and immediately snap-frozen in liquid nitrogen. Frozen serum was stored at -80°C prior to LC-MS extraction and analysis. In addition, cecal contents, liver, kidney and urine samples were collected and flash frozen in liquid nitrogen.

### Metabolite extraction

Trp metabolites were quantified using LC-MS, following an established method with some modifications^15^. Serum and urine were extracted by mixing 25 µL of sample, thawed on ice, with 100 µL of ice-cold methanol extraction solvent containing indole-3-acetic acid-d4 and KA-d5 (Caymen Scientific). The mixture was vortexed and incubated at −20 °C for 30 min. After centrifugation at 12,000 × g for 15 min at 4°C, 90 µL of the supernatant was collected, evaporated to dryness using a VacuFuge (Eppendorf, Hamburg, Germany), and dissolved in 45 µL of 10% acetonitrile containing 1 µM chlorpropamide. Following another centrifugation at 17,000 × g for 15 min at 4°C, the supernatants were transferred to autosampler vials for LC-MS analysis.

Fecal and cecal content extracts were prepared by adding 50 μL of ice-cold 50% acetonitrile in water (v/v) extraction solvent containing indole-3-acetic acid-d4 and KA-d5 to 10 mg of sample. Samples were homogenized using 1.0 mm zirconia beads (Biospec Products, Bartlesville, OK, USA) and a Bead Blaster 24 (Benchmark Scientific, Sayreville, NJ, USA) prior to sonication for 20 min using a chilled Branson 3800 sonicating water bath (Danbury, CT, USA). Extracts were placed at −20 °C for 30 min for protein precipitation before centrifuging at 17,000 × g for 15 min at 4°C. Supernatant was transferred to a clean tube and diluted 5-fold with LC-MS grade water containing internal standards and chlorpropamide.

For tissue extractions, 50 mg of liver and kidney tissue was added to 1 mL of ice-cold methanol extraction solvent containing indole-3-acetic acid-d4 aKA-d5. Samples were homogenized as previously described before centrifugation at 17,000 × g for 15 min at 4°C. 400 μL of supernatant was transferred to a clean and stored at −20 °C for 30 min for protein precipitation. Samples were centrifuged at 17,000 × g for 15 min at 4°C, 350 μL was transferred to a clean tube, and a VacuFuge was used to completely dry the sample. Samples were solubilized in 10% acetonitrile/water (v/v) containing internal standards and chlorpropamide.

To extract metabolites, cells were thawed at room temperature, water was added to each well and a cell lifter was used to remove cells from the surface of the plate. Cell lysate was added to an equal volume of LC-MS grade methanol (Fisher Scientific) for a final concentration of 50% methanol in water (v/v). Samples were placed at -20°C for 30 min to precipitate proteins, centrifuged (17,000 x *g*, 15 min, 4°C), and a portion of supernatant was transferred to a clean tube. Samples were dried using an Eppendorf VacuFuge (Hamburg, Germany) to complete dryness and resuspended in 10% acetonitrile in water (v/v) with 1 μM chlorpropamide to serve as an instrument calibrant. LC-MS conditions were the same as previously described.

### Quantification of Trp metabolites

Metabolite quantification was performed using reverse-phase ultra-high-performance liquid chromatography (UHPLC) on a Nexera UHPLC system (Shimadzu, Columbia, MD), equipped with a Waters BEH C18 column (2.1 × 100 mm, 1.7 µm particle size) maintained at 55 °C. Separation was achieved using a 20-min gradient of aqueous acetonitrile at a flow rate of 250 µL/min. Mobile phase A was water with 0.1% formic acid, and mobile phase B was acetonitrile with 0.1% formic acid. The gradient program began at 97% A and 3% B, ramped to 45% B by 10 minutes, increased to 75% B at 12 min, held at 75% B until 17.5 minutes, and then returned to the initial conditions.

The UHPLC eluate was introduced into a ZenoTOF 7600 mass spectrometer (SCIEX, Framingham, MA) via a Turbo V ion source. Data acquisition was carried out in high-resolution multiple reaction monitoring (MRM) mode, with mass accuracy maintained through calibration using a SCIEX-provided calibration solution.

Analyte recovery and matrix effects for each sample type were assessed and used to correct quantification values.

Recovery was calculated as:

**Recovery = (Peak area of spiked analyte before extraction – Peak area of endogenous analyte) / Peak area of spiked analyte after extraction – Peak area of endogenous analyte)**.

Matrix effect was calculated as:

**Matrix effect = (Peak area of spiked analyte – Peak area of endogenous analyte) / Peak area of spiked analyte in water**.

Limits of quantification (LOQ) were determined based on signal-to-noise ratios from ten replicate injections, analyzed using MSDIAL v.5.4.241004. Quantification was based on standard curves generated by linear regression from six serial dilutions of a standard mixture. Internal standards included chlorpropamide, indole-3-acetic acid-d4, and KA-d5.

### Cell culture

HepG2 and Hepa1c1c7 cells were obtained from ATCC. The dioxin-response element driven reporter cell lines Hepa1.1 were a kind gift from Michael S. Denison (University of California, Davis, CA) and HepG2 40/6 were generated as previously described^69^. These cell lines were grown in αMEM (Sigma, St. Louis, MO, USA) supplemented with 8% FBS (Gemini BioProducts) and 100 units/ml penicillin/0.1 mg/ml streptomycin (Sigma). Primary fibroblast-like synoviocytes isolated from a rheumatoid arthritis patient were cultured in complete synoviocyte medium with 100 units/ml penicillin/0.1 mg/ml streptomycin (Cell Applications, Inc., San Diego, CA) Cells were routinely tested for the presence of mycoplasma utilizing the Invivogen Mycostrip Detection Kit (San Diego, CA, USA) and only mycoplasma-free cultures were utilized.

### Cell-based luciferase assay

HepG2 40/6 cells were grown to confluency in 12 well plates and the medium replaced with 1 ml of αMEM supplemented with 5 mg/ml BSA, 100 units/ml penicillin/0.1 mg/ml streptomycin prior to treatment. All compounds were placed in DMSO and added to culture at a final concentration of 0.1%. After 4 h incubation, cells were washed with PBS, lysed using 100 µl of lysis buffer 25 mM Tris-phosphate pH 7.8,2 mM dithiothreitol, 2 mM 1,2-diaminocyclohexane-N,N,N′,N′-tetraacetic acid, 10% (v/v) glycerol, 1% (v/v) Triton X-100) and stored at −80°C until analysis. Luciferase activity was measured using a TD-20e luminometer (Turner Biosystems Inc.) and luciferase substrate (Promega, Madison, WI, USA), as described in the manufacturer’s instructions.

### RNA isolation and RT-qPCR

Either HepG2 or Hepa1 cells were seeded into 12 well plates and incubated overnight in maintenance media. The cells were washed with PBS followed by addition of αMEM supplemented with 5 mg/ml BSA or 1% Fetal Bovine Serum, 100 units/ml penicillin/0.1 mg/ml streptomycin prior to treatment. All compounds were placed in DMSO and added to culture at a final concentration of 0.1%. After 4 h incubation, cells were washed with PBS, then 1 ml of TRI Reagent (Sigma, MO, USA) was added to each well. RNA was isolated according to the manufacturer’s instructions. The purified total RNA was used to generate cDNA using the High Capacity cDNA Archive Kit following the manufacturer’s protocol (Applied Biosystems, MA, USA). Quantitative real-time PCR of mouse *Cyp1a1* or human *CYP1A1* was performed and normalized to β-actin (Table S4). The PCR was conducted using Perfecta SYBR Green (QuantaBio, MD, USA) in a Bio-Rad CFX96 real-time System (BioRad CA,USA).

### Cellular uptake of Trp metabolites

Six well cell culture plates were seeded with 100,000 HepG2 cells per well in maintenance media. To perform the assay, maintenance media was aspirated, HepG2 cells were washed with PBS, and Hanks Balanced Salt Solution (Gibco, Waltham, MA, USA) was added to cells. Cells were allowed to recover for 30 minutes before treatment with 100 μM (final concentration) of test compound. Plates were incubated for 90 min. At the end of the treatment exposure, cells were washed with PBS and snap frozen in liquid nitrogen prior to storage at -80°C.

### In vitro IAA conjugation

Renal and hepatic tissue were dissected from 8-12 week old C57BL6/J mice. Tissues were chilled on ice, minced and homogenized, using a glass Dounce homogenizer (Wheaton), in 4 volumes ice-cold homogenization buffer (150 mM KCl, 2 mM HEPES, pH8) supplemented with protease inhibitor cocktail (Roche). Crude homogenates were centrifuged (600 x *g*, 10 min, 4°C) to remove debris. Supernatants were recentrifuged (14, 300 x *g*, 20 min, 4°C) to pellet the intact mitochondrial fraction. Mitochondria were resuspended and permeabilized in 5 ml ice-cold homogenization buffer supplemented with 0.03% (v/v) Triton X-100. Samples were centrifuged (162, 000 x *g*, 30 min, 4°C). Mitochondrial supernatants were aliquoted and stored at -80°C until use.

1 ml conjugation reactions were initiated by addition of 5 μl vehicle (DMSO) or 100 mM IAA (500 μM final reaction concentration) to reaction mixes comprising 12.5 mM Tris-HCl, 2 mM HEPES, 5 mM glycine, pH8 supplemented with 340 μg renal or 500 μg hepatic mitochondrial extracts. Reactions were incubated at 37°C for 30 min with shaking. Reactions were quenched by addition of 1 ml ice-cold 100% methanol supplemented with 1 μM chlorpropamide (as internal extraction standard). Quenched samples were incubated at -20°C for 30 min to precipitate the protein fraction prior to centrifugation (16, 000 x *g*, 15 min, 4°C). 0.9 ml protein-free supernatants were transferred to fresh tubes and evaporated to dryness under vacuum. Dried samples were solubilized in 100 μl 3% methanol with sonication. Samples were centrifuged (16,000 x *g*, 15 min, 4°C) prior to transfer to LCMS autosampler vials and subsequent LCMS analysis.

### Microbial de/conjugation assays

Fecal pellets were aerobically collected from C57BL/6 mice, samples were immediately weighed and transferred to anaerobic conditions in a Coy vinyl anaerobic chamber (Grasslake, MI, USA) under 20% carbon dioxide, 20% hydrogen and 5% nitrogen. Samples were diluted (1:100 w/v) with pre-reduced brain heart infusion agar (Gibco) to generate initial culture conditions. Technical triplicates were prepared by treating cultures with 100 μM (final concentration) d5-IAA or d5-IAA-Gly (0.1% DMSO, final concentration), and anaerobically incubated at 37C for 1, 3, 9, and 24 hours. At each sampling timepoint, an equal volume of culture was added to 100% LC-MS grade methanol containing 1 μM chlorpropamide (internal standard) and stored at -20°C until all time points were complete. Following the final sampling, samples were incubated at 20°C for 30 min to precipitate protein, and centrifuged (16,000 x g, 15 min, 4°C). Supernatant was transferred to a fresh tube for concentration using an Eppendorf VacuFuge until sample reached complete dryness. Samples were resolubilized in 10% LC-MS grade acetonitrile in water prior to LC-MS analysis.

### Computational docking analysis

Molecular docking was conducted to evaluate the binding interactions of indirubin, IAA and its amino acid conjugates (IAA-Gly and IAA-Ser) within the PAS-B domain of the human AHR. AutoDock Vina 1.2.0 was used with a homology model derived from the cryo-EM structure of the AHR complex (PDB: 7ZUB)^4,70^. Ligand structures were retrieved from PubChem, energy-minimized in *The PyMOL Molecular Graphics System, Version 3.0, Schrödinger and DeLano, LLC* ^71^, and optimized for docking using AutoDock Tools.^72^ For each ligand, the docking grid (45.0 × 37.5 × 27.0 Å) was centered on the indirubin binding pocket, and binding energies along with dissociation constants (Kᴅ) were determined using AutoDock 4.2 parameters following established methods^15,73^. The highest affinity pose for each ligand was selected for further structural analysis, while an average Kᴅ was computed across nine docking replicates. Ligand-protein interactions were visualized in PyMOL 3.0 and PoseView.^70,74–77^

### Chemical antioxidant assay

The intrinsic chemical antioxidant potential of IPA, IAA and their respective glycine conjugates were quantified using the ORAC antioxidant (AOX-2) assay kit (Zenbio). Briefly, test compounds were solubilized in Dimethyl formamide (DMF) and subsequently serially diluted in AOX-2 assay buffer such that final indicated serum concentrations were achieved in a final DMF concentration of 0.03% (v/v). Kinetic fluorescence assays were performed following manufacturer’s instructions.

### Statistical analysis

Data sets were compared for statistical significance using one-way ANOVA with Tukey’s multiple comparison post-test or a two-way ANOVA with Sidak multiple comparison post-test, or *t*-test in GraphPad Prism v 9.4.1 (GraphPad Software, San Diego, CA, USA). Values of p<.05 were considered statistically significant (*: p<.05; **: p<.01; ***: p<.001; ****; p<.0001).

## Data availability

The authors declare that the data supporting the findings of this study reported in this manuscript are available along with the accompanying supplementary information files. The metabolomics data are available at the NIH Common Fund’s National Metabolomics Data Repository (NMDR) website, the Metabolomics Workbench, https://www.metabolomicsworkbench.org where it has been assigned Study ID ST002462. All other data can be accessed directly via its Project DOI: http://dx.doi.org/10.21228/M89H8W.

## Acknowledgements

The co-authors would like to acknowledge the Huck Institutes’ Metabolomics Core Facility (RRID:SCR_023864) for use of the Sciex ZenoTOF 7600 LC-MS. Graphical representations were created using BioRender. We thank Marcia H. Perdew for critically reviewing this manuscript.

## Author contributions

E.W.M., I.A.M., A.J.A., and G.H.P. conceptualized the experiments. E.W.M., F.C., I.A.M. conducted LC-MS studies. A.J.A. performed the AHR modeling studies. D.M.C., E.U.D., and E.W.M. performed the RT-PCR and cell culture experiments. K.G., D.D., and S.G.A were responsible for the organic synthesis and F.H. completed the NMR analysis. K.S.P., and P.M.K-E. provided the human serum samples. C.B.M., J.E.B. and A.D.P provided guidance on experimental design, data analysis and manuscript writing. E.W.M., I.A.M., and G.H.P. wrote the manuscript.

## Additional information

## Supplementary information

Online version contains supplementary material is available

