## supplement for "Interkingdom glycine conjugates of indole-3-carboxylates are Ah receptor ligands"

#### Table of Contents

Supplementary Tables…………………………………………………………………………2
Supplementary Figures……………………………………………………………………..5
Chemical Synthesis Methods…………………………………………………………..11
Mass Spectra…………...…………………………………………………………………..18
NMR Spectra……………………………………………………………………………..25

**Table S1. Limit of metabolite detection/quantification**

| Metabolite | LOD (nM; S/N>3) | LOQ (nM; S/N>10) |
|---|---|---|
| **IAA** | 60 | 250 |
| **IAA-Ala** | 150 | 300 |
| **IAA-Gln** | 60 | 150 |
| **IAA-Gly** | 60 | 150 |
| **IAA-Ser** | 150 | 300 |
| **IAA-Thr** | 150 | 300 |
| **IPA** | 30 | 60 |
| **IPA-Gly** | 20 | 60 |
| ***ICA** | 80 | 150 |
| ***ICA-Gly** | 80 | 150 |
| ****IACA** | 80 | 150 |
| ****IACA-Gly** | 80 | 150 |

*ICA= Indole-3-carboxylic acid

**IACA= indole-3-acrylic acid

**Table S2. Correlation statistics**

| | | | **Pearson** | | **Spearman** | | **Kendall** | |
|---|---|---|---|---|---|---|---|---|
| | | | r | p | rho | p | tau B | p |
| IAA | - | IPA | -0.115 | .480 | -0.065 | .689 | -0.036 | .755 |
| IAA | - | I3A-Gly | 1.000 | < .001 | 1.000 | < .001 | 1.000 | < .001 |

**Table S3. Computational docking analysis of indole-3-acetic acid and amino acid conjugates in the PAS-B domain of the human AHR.**

| **Substrate** | | **Autodock Vina** | |
|---|---|---|---|
| | | ***Structure-based Homology Model*** | |
| | | **Human AhR PAS B domain** | |
| | | **Binding Energy** [a] (kcal/mole) | **Dissociation Constant** [b] *[$K_D$] μM* |
| **Indirubin** | *Max* [c] | **-12.8** | ***0.00044*** |
| | *Avg* [d] | *-12.0 ± 0.40* | *0.0025 ± 0.0023* |
| **Indole-3-Acetic acid (IAA)** | *Max* | **-7.6** | ***2.6*** |
| | *Avg* | *-7.2 ± 0.26* | *5.34 ± 2.89* |
| **IAA-Glycine** | *Max* | ***-8.3*** | ***0.81*** |
| | *Avg* | *-8.0 ± 0.22* | *1.46 ± 0.53* |
| **IAA-Serine** | *Max* | ***-8.3*** | ***0.81*** |
| | *Avg* | *-8.0 ± 0.26* | *1.58 ± 0.75* |

[a] Substrate Binding Energies were derived computationally using Autodock Vina.

[b] Dissociation binding constants ($K_D$) (in the micromolar (*μ*M) scale), were derived computationally from Autodock Vina data using the Autodock 4.2 inhibitory constant conversion scale, based on the equation: $y = 0.5982\ln(x) - 12.304$.

[c] Max refers to the low-energy Vina docking solution (in kcal/mole), with the highest predicted binding affinity [$K_D$] for the human AhR PAS B domain (PDB: 7ZUB [1]).

[d] Avg refers to the average or cumulative dissociation constant and binding energy (± the standard deviation) for the Vina docking poses obtained for each individual substrate:model docking combination (N=9).

**Table S4. Quantitative qPCR primers**

| Gene | Forward Primer (5'->3') | Reverse Primer (5'->3') |
|---|---|---|
| Human *CYP1A1* | GATTGAGCACTGTCAGGAGAAGC | ATGAGGCTCCAGGAGATAGCAG |
| Human *β-ACTIN* | CACCATTGGCAATGAGCGGTTC | AGGTCTTTGCGGATGTCCACGT |
| Human *SERPINB2* | GCTGTTTGGTGAGAAGTCTGCG | CTGCACATTCTAGGAAGTCTACTG |
| Mouse *Cyp1a1* | CTCTTCCCTGGATGCCTTCAA | GGATGTGGCCCTTCTCAAATG |
| Mouse *β-Actin* | CATTGCTGACAGGATGCAGAAGG | TGCTGGAAGGTGGACAGTGAGG |

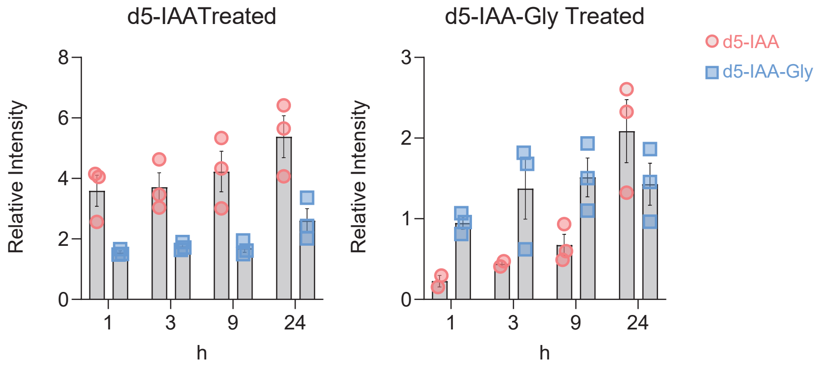


**Fig. S1. Conjugation of d5-IAA and deconjugation of d5-IAA-Gly by mouse gut microbiota.** Mouse feces were freshly collected in anerobic conditions, pooled, diluted in BHI broth, and anaerobically incubated at 37°C with 100 μM d5-IAA or d5-IAA-Gly. Samples were collected at 1, 3, 9, and 24 h to detect conjugation or deconjugation of the treatment compound using LC-MS. Deuterated treatment compounds were used to specifically assess microbial metabolism within the treatment window. Relative intensity describes the peak area of the analyte divided by the peak area of the internal standard (chlorpropamide).

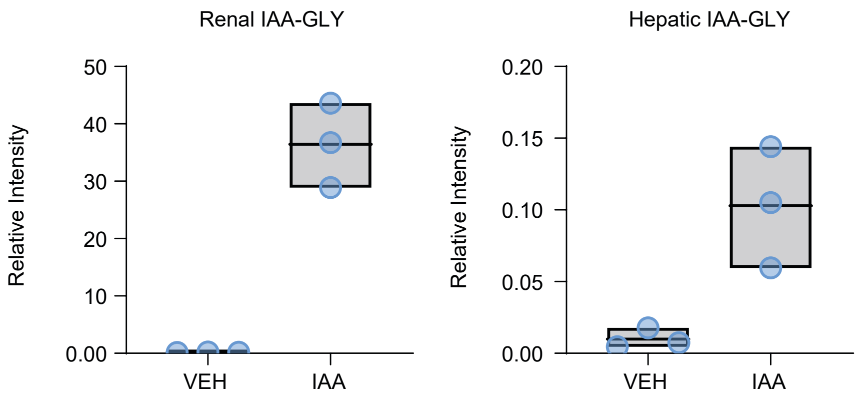


**Fig. S2. Mouse hepatic and renal mitochondrial extracts are permissive for glycine conjugation of IAA.** Mouse hepatic and renal mitochondrial extracts were incubated for 30 min at 37°C with vehicle or 500 mM IAA. Incubations were quenched with ice-cold 100% methanol. Samples were extracted, dried and solubilized in 3% methanol for subsequent LCMS analysis. Data represented as relative abundance of IAA-Gly normalized to mg protein (mean ± SEM, n=3).

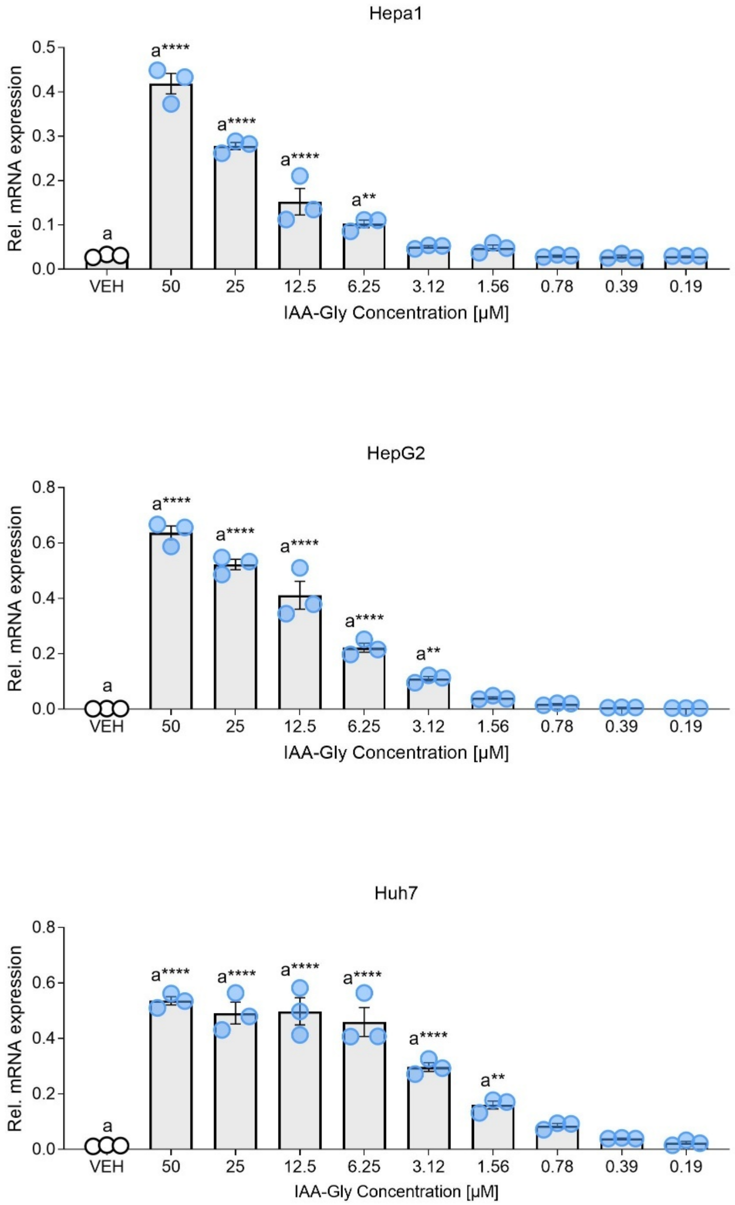


**Fig. S3. Dose-response analysis of IAA-Gly across multiple cell lines.** Cells were treated for 4 h with VEH or IAA-Gly, at the doses indicated. Quantitative PCR analysis for relative mRNA expression was performed. Data were normalized to the housekeeping gene beta-actin. Data are represented at ± SEM, n=3, and statistical analysis was performed using one-way ANOVA. *: $p < 0.05$, **: $p < 0.01$, ***: $p < 0.001$, ****: $p < 0.0001$.

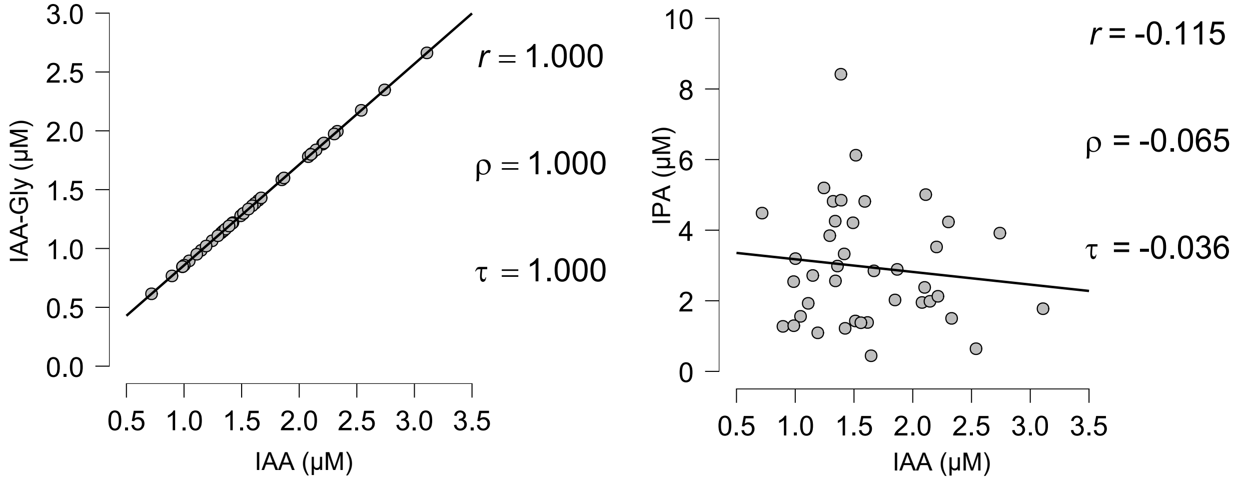


**Fig. S4. Human serum concentrations of IAA and IAA-Gly are correlated, while IAA and IPA are not.** Pearson's r, Spearman's rho and Kendall's tau-b values indicate a highly significant ($p < 0.001$) correlation between metabolites.

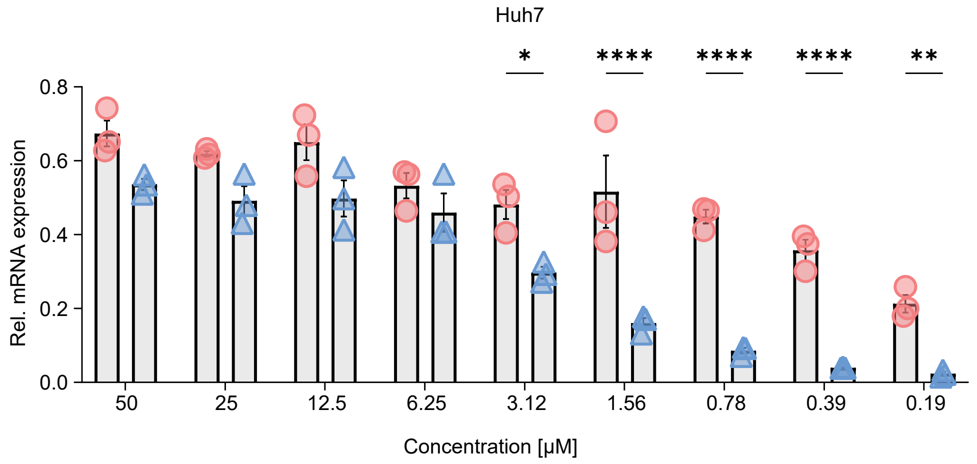


**Fig. S5. *CYP1A1* expression in Huh7 cells is dependent on dose.** Cells were treated for 4h IAA or IAA-Gly, at the doses indicated. Quantitative PCR analysis for relative *CYP1A1* mRNA expression was performed. Data were normalized to the housekeeping gene beta-actin and the vehicle baseline subtracted. Data are represented at ± SEM, n=3, and statistical analysis was performed using 2-way ANOVA followed by Sidak multiple-comparison test. *: $p < 0.05$, **: $p < 0.01$, ***: $p < 0.001$, ****: $p < 0.0001$

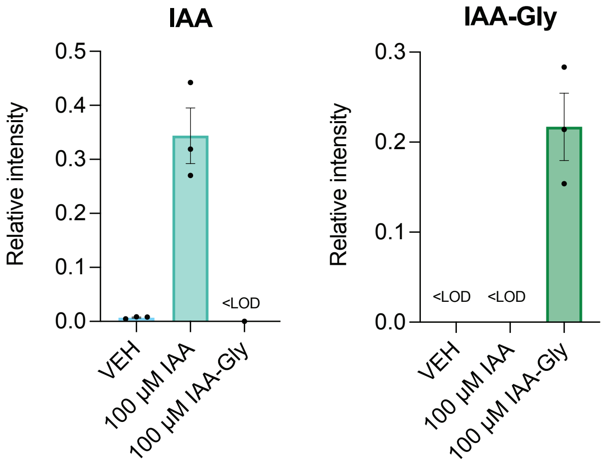


**Fig. S6. IAA is not conjugated with glycine and IAA-Gly is not deconjugated in HepG2 cells.** HepG2 cells in 6-well plates were exposed to either 100 μM IAA or IAA-Gly for 24 h, washed with PBS, and the cells scraped into 50% methanol. Relative IAA and IAA-Gly levels were determined by LC-MS.

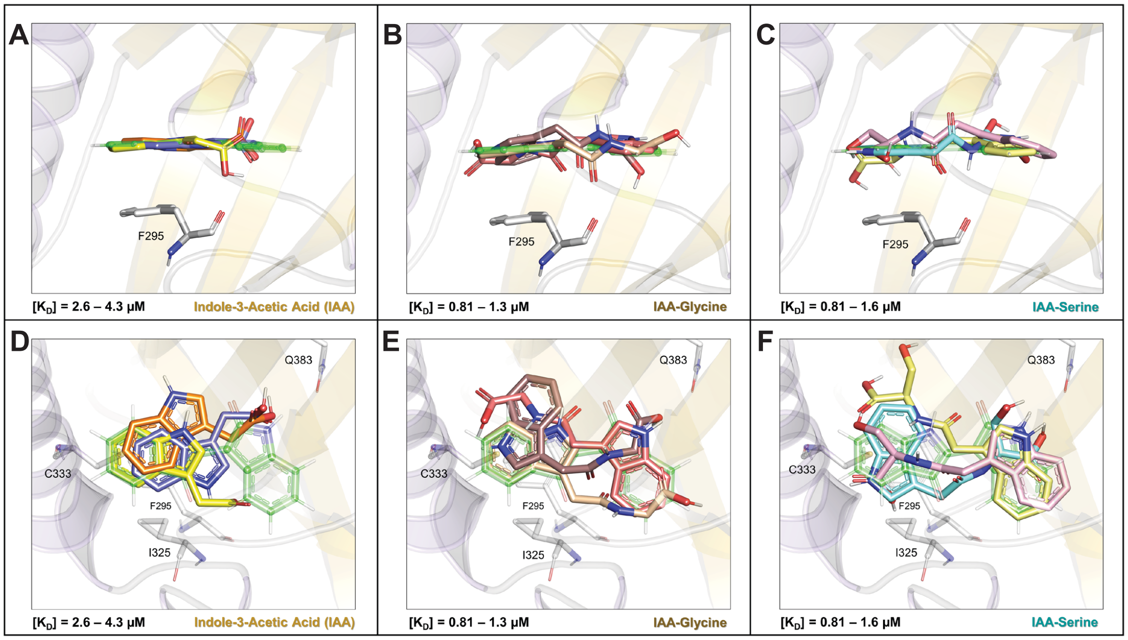


**Fig. S7. Exploring the structural basis for AHR agonism through superposition with indirubin in the PAS B domain. A.** To further highlight differences in the structural recognition of these related ligands by the AHR, we superimposed three high affinity docking results for IAA (yellow stick; [$K_D$] = 2.6 μM; orange stick; [$K_D$] = 3.6 μM; dark blue stick; [$K_D$] = 4.3 μM) onto the coordinates of indirubin, to highlight how this substrate can consistently dock the PAS B domain in a highly planar configuration across a range of binding energies. **B.** IAA-glycine showed greater heterogeneity with respect to planar binding across multiple docking results (tan stick; [$K_D$] = 0.81 μM; brown stick; [$K_D$] = 0.81 μM; salmon stick; [$K_D$] = 1.30 μM), however highly planar binding configurations were noted for both high (tan stick) and moderate (salmon stick) affinity results. **C.** In contrast, IAA-serine displayed its most kinked docking conformations, with respect to indirubin, when bound to the PAS-B domain at high affinity (pink stick; [$K_D$] = 0.81 μM), only displaying more planar results typical of a good AHR agonists at lower binding affinity (cyan stick; [KD $K_D$] = 1.13 μM; light yellow stick; [$K_D$] = 1.6 μM). Collectively, these docking results suggest that ligand planarity, particularly in configurations resembling indirubin, is a critical determinant of AHR agonism. Departures from planarity in the docking poses, as seen in IAA-serine, correlate with reduced receptor activation despite high affinity binding. **D.** To further highlight differences in ligand binding among the three AHR agonists, we also assessed superposition trends in relation to indirubin's bi-indole structure.  Three high affinity docking results for IAA (yellow stick; [$K_D$] = 2.6 μM; orange stick; [$K_D$] = 3.6 μM; dark blue stick; [$K_D$] = 4.3 μM) are shown superimposed on the coordinates of indirubin, highlighting how this ligand can consistently dock the PAS B domain in a planar configuration, but only the high-affinity result replicates the prototype indole ring positioning relative to F295. **E.** Multiple docking results for IAA-glycine (tan stick; [$K_D$] = 0.81 μM; brown stick; [$K_D$] = 0.81 μM; salmon stick; [$K_D$] = 1.30 μM) are also shown with respect to indirubin, demonstrating the ligand's ability to bind in two prototypical orientations, with its indole ring closely superimposing on the bi-indole structure of indirubin in both the high (tan stick) and moderate (salmon stick) affinity results. **F.** In contrast, IAA-serine which displays a kinked, docking conformation, with respect to indirubin when bound at high affinity (pink stick; [$K_D$] = 0.81 μM), only displays a planar configuration at lower binding affinity (cyan stick; [$K_D$] = 1.13 μM; light yellow stick; [$K_D$] = 1.6 μM). However, none of the planar binding configurations of IAA-serine effectively position the indole ring relative to indirubin's bi-indole structure. Since IAA-serine is a poor AHR ligand, this suggests that ligand recognition is primarily driven by hydrophobic contacts with residues I325, C333, and F295 in the primary binding site (Fα/Gβ site) near the pocket's opening, rather than by interactions with H291 and L353 deeper in the secondary pocket near the Cα helix. These docking results collectively suggest that effective AHR ligands must adopt a planar configuration that properly engages the hydrophobic cleft in the primary binding site, which seems highly evolved to recognize specific modified indole structures.

#### 1. Synthesis of Indole-3 acetic acid-Serine conjugate (IAA-Serine Conjugate)

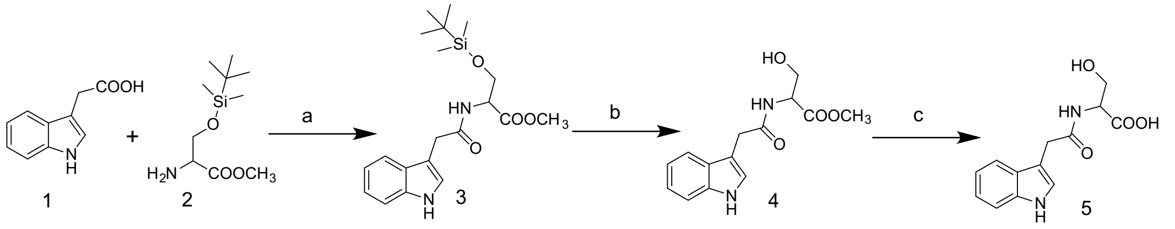


Reagents and Conditions: a) TBTU (1 eq), DIPEA (2.5 eq), DMF, RT, 24 hrs.; b) TBAF (3 eq), THF, RT c) NaOH, MeOH, RT

**Scheme 1. Synthesis of indole-3-acetic acid-serine conjugate**

Indol-3-acetic acid **1** (500 mg, 2.85 mmol), O-(*t*-butyldimethylsilyl)-L-serine methyl ester **2** (1.1 eq, 731 mg, 3.13 mmol), TBTU (1 eq, 915 mg), and DIPEA (2.5 eq) were stirred in anhydrous DMF (20 mL) overnight at room temperature. The reaction was monitored on TLC to complete the reaction, and the mixture was poured into water, extracted with ethyl acetate, washed with 10% sodium bicarbonate, 5% HCl, and with water, dried, and concentrated in a rotavapor to obtain **3**. The crude product was dissolved in THF and stirred with tetrabutylammonium fluoride for 6-8 hours at RT. The reaction mixture was extracted with ethyl acetate, washed with water, dried, concentrated, and purified by silica gel column chromatography using hexane/ethylacetate (3:1) as eluent to achieve **4**. Product **4** was hydrolyzed with 1N NaOH in methanol, neutralized with acetic acid, and concentrated. The product was dissolved again in methanol and filtered. The filtrate was concentrated and dried in a high vacuum pump to obtain IAA-serine conjugate **5** (360 mg, 48%) as a white solid. The structure of indole-3 acetic acid-serine was confirmed by nuclear magnetic resonance (NMR) and mass spectra analysis.

#### 2. Synthesis of Indole-3 acetic acid-Alanine conjugate (IAA-Alanine Conjugate)

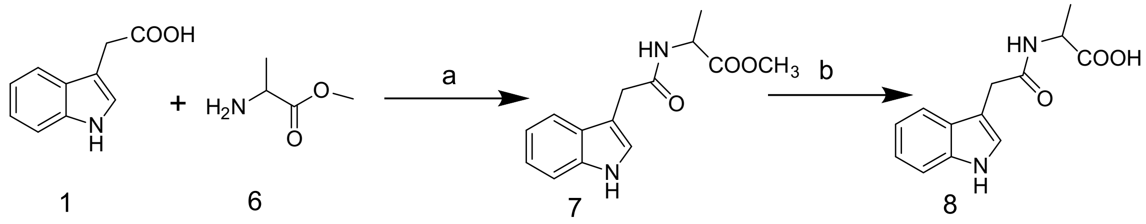


Reagents and Conditions: a) TBTU (1 eq), DIPEA (2.5 eq), DMF, RT, 24 hrs.; b) NaOH, MeOH-THF, RT

**Scheme 2. Synthesis of indole-3-alanine conjugate**

Indol-3-acetic acid **1** (530 mg, 3.02 mmol) was stirred in anhydrous DMF (15 mL) at RT with L-alanine methyl ester hydrochloride **6** (1.1 eq, 464 mg, 3.32 mmol), TBTU (1 eq, 970 mg), and DIPEA (2.5 eq). The completion of the reaction was monitored on TLC, and the mixture was poured into water, extracted with ethyl acetate, washed with 10% sodium bicarbonate, 5% HCl, water, dried, and concentrated in a rotavapor to obtain **7**. Product **7** was hydrolyzed with 1N NaOH in methanol-THF, neutralized with HCl, and concentrated. The crude product was extracted with ethyl acetate, washed with water, dried, concentrated, and purified by silica gel column using $CH_2Cl_2$/MeOH (95:5) to yield IAA-alanine conjugate **8** (410 mg, 55%) as a white solid. Nuclear magnetic resonance (NMR) and mass spectra analysis confirmed the structure of IAA-alanine conjugate.

##### 3. Synthesis of Indole-3 acetic acid-Threonine conjugate (IAA-Threonine Conjugate)

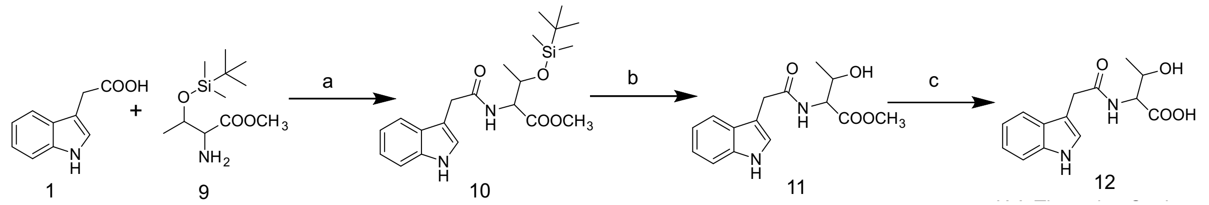


Reagents and Conditions: a) TBTU (1 eq), DIPEA (2.5 eq), DMF, RT, 24 hrs.; b) TBAF (3 eq), THF, RT c) LiOH, MeOH-THF, RT

**Scheme 3. Synthesis of indole-3-acetic acid-threonine conjugate**

To the mixture of indol-3-acetic acid **1** (520 mg, 2.96 mmol) in anhydrous DMF (20 mL) was added methyl 2-amino-3-((tert-butyldimethylsilyl)oxy)butanoate **9** (1.1 eq, 806 mg, 3.26 mmol), TBTU (1 eq, 950 mg), and DIPEA (2.5 eq) and stirred overnight at room temperature. The completion of the reaction was monitored on TLC, and the mixture was poured into water, extracted with ethyl acetate, washed with 10% sodium bicarbonate, 5% HCl, water, dried, and concentrated in a rotavapor to obtain **10**. Product **10** was dissolved in THF and stirred with tetrabutylammonium fluoride for 7-8 hours at RT and extracted with ethyl acetate, washed with water, dried, concentrated, and purified by silica gel column chromatography using hexane/ethylacetate (3:1) as eluent to give **11**. Product **11** was hydrolyzed with 1N LiOH in MeOH-THF, neutralized with 2%HCl, and concentrated. The product was purified by silica gel column using $CH_2Cl_2$/MeOH (5:1) to obtain **12** (335 mg, yield 39%) as a white solid. The structure of indole-3 acetic acid-threonine conjugate was confirmed by nuclear magnetic resonance (NMR) and mass spectra analysis.

**4. Synthesis of Indole-3 acetic acid-Glutamine conjugate (IAA-Glutamine Conjugate)**

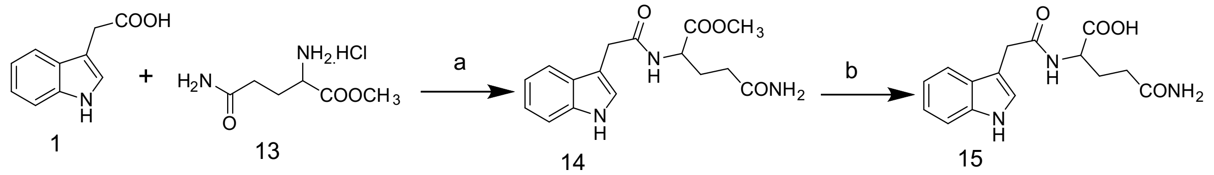


Reagents and Conditions: a) TBTU (1 eq), DIPEA (2.5 eq), DMF, RT, 24 hrs.; b) NaOH, MeOH-THF, RT

**Scheme 4. Synthesis of indole-3-acetic acid-glutamine conjugate**

Indol-3-acetic acid **1** (480 mg, 2.73 mmol), L-glutamine methyl ester hydrochloride **13** (1.1 eq, 590 mg, 3.0 mmol), TBTU (1 eq, 876 mg), and DIPEA (2.5 eq) were added to anhydrous DMF (15 mL) and stirred overnight at room temperature. After completion of the reaction, the mixture was poured into water, extracted with ethyl acetate, washed with 10% sodium bicarbonate and 5% HCl, and with water, dried and concentrated in a rotavapor to obtain **14**. It was hydrolyzed with 1N NaOH in MeOH-THF, neutralized with 2%HCl, and concentrated. The product was dissolved in a minimum quantity of methanol and filtered. Ethyl acetate was added dropwise to the filtrate, and the precipitate was filtered and dried to obtain a compound of **15** (270 mg, yield 32%) as a white solid. The structure of IAA-glutamine conjugate was confirmed by nuclear magnetic resonance (NMR) and mass spectra analysis.

#### 5. Synthesis of Indole-3-acetic acid-Glycine conjugate (IAA-Glycine conjugate)

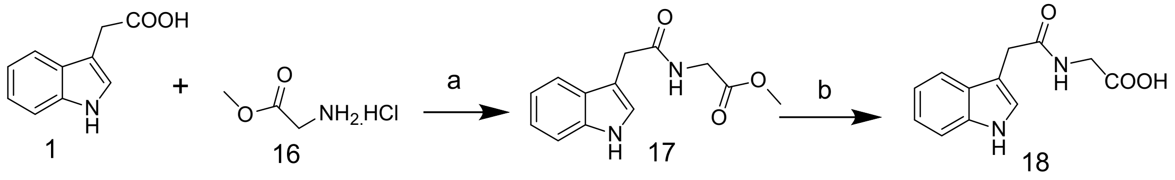


Reagents and Conditions: a) TBTU (1 eq), DIPEA (2.5 eq), DMF, 24 hrs.; b) LiOH, THF-MeOH

**Scheme 5. Synthesis of indole-3-acetic acid-glycine conjugate**

Indol-3-acetic acid **1** (120 mg, 0.68 mmol), glycine methyl ester hydrochloride **16** (1.1 eq, 94 mg, 0.75 mmol), TBTU (1 eq, 218 mg), and DIPEA (2.5 eq) were dissolved in anhydrous DMF (8 mL), and the reaction mixture was stirred overnight at room temperature. The reaction was monitored on TLC to complete the reaction, and the mixture was poured into water, extracted with ethyl acetate, washed with water, dried, and concentrated in a rotavapour. The crude product **17** was dissolved in a methanol-THF mixture and stirred with 1M LiOH for ~3 hours to hydrolyze the ester group into carboxylic acid and acidified with 1M HCl. The methanol-THF was removed by rotavaporation, extracted with ethyl acetate, washed with water, dried, and concentrated. Finally, the crude product was purified by silica gel column using EtOAc: MeOH (2:1) as eluents, and the product was dried in a high vacuum pump to obtain Indole-3-acetic acid-glycine **18** (55 mg, yield 45%) as a white solid. The structure of indole-3 acetic acid-glycine was confirmed by nuclear magnetic resonance (NMR) and mass spectra analysis.

d5-indole-3-acetic acid-glycine conjugate was synthesized utilizing the same procedure except for the use of d5-indole-3-acetic acid (Cayman Chemical Co., Ann Arbor, MI, USA).

**6. Synthesis of indole-3-propionic acid-Glycine conjugate (IPA-Glycine conjugate)**

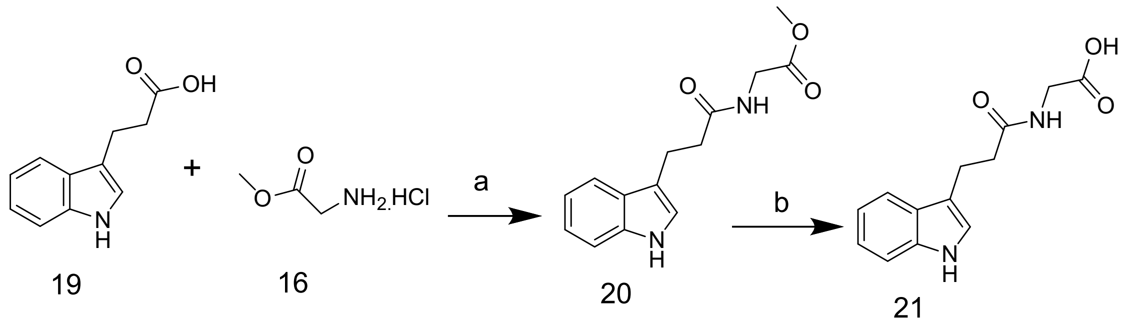


Reagents and Conditions: a) TBTU (1 eq), DIPEA (2.5 eq), DMF, 24 hrs.; b) LiOH, THF-MeOH

**Scheme 6. Synthesis of indole-3-propionic acid-glycine conjugate**

3-(Indol-3-yl) propionic acid **19** (378 mg, 2.0 mmol), glycine methyl ester hydrochloride **16** (1.1 eq, 276 mg, 2.2 mmol), TBTU (1 eq, 642 mg), and DIPEA (2.5 eq) were dissolved in anhydrous DMF (8 mL), and the reaction mixture was stirred overnight at room temperature. The reaction was monitored on TLC to complete the reaction, and the mixture was poured into water, extracted with ethyl acetate, washed with water, dried, and concentrated in a rotavapour. The crude product **20** was dissolved in a methanol-THF mixture and stirred with 1M LiOH for ~3 hours to hydrolyze the ester group into carboxylic acid and acidified with 1M HCl. The methanol-THF was removed by rotavaporation, extracted with ethyl acetate, washed with water, dried, and concentrated. Finally, the crude product was purified by silica gel column using EtOAc: MeOH (2:1) as eluents, and the product was dried in a high vacuum pump to obtain Indole-3-propionic acid-glycine **21** (255 mg, yield 67%) as a white solid. The structure of indole-3-propionic acid-glycine was confirmed by nuclear magnetic resonance (NMR) and mass spectra analysis.

##### 7. Synthesis of Indole-3-carboxylic acid-Glycine conjugate

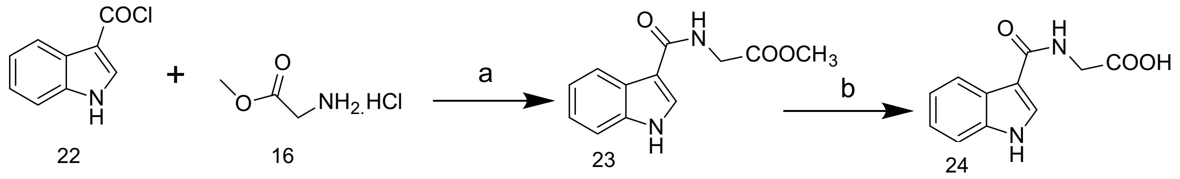


Reagents and Conditions: a) TEA (2.5 eq), DMF, 8 hrs.; b) NaOH, THF-MeOH

**Scheme 7. Synthesis of indole-3-carboxylic acid-glycine conjugate**

Glycine methyl ester hydrochloride **16** (1.1 eq, 385 mg, 3.06 mmol), and TEA (2.2 eq, 6.73 mmol, 681 mg) were dissolved in anhydrous DMF (10 mL) and cooled on ice with stirring. 1H-Indole-3-carbonyl chloride **22** (500 mg, 2.79 mmol) was dissolved in methylene chloride and slowly dropped into the reaction mixture and stirred at room temperature for 6 h. The reaction was monitored on TLC to complete the reaction, and the mixture was poured into water, extracted with ethyl acetate, washed with water, dried, and concentrated in a rotavapor. The crude product 22 was dissolved in a methanol-THF mixture and stirred with 1M NaOH for ~5 hours to hydrolyze the ester group into carboxylic acid and acidified with 1M HCl. The methanol-THF was removed by rotavaporation, extracted with ethyl acetate, washed with water, dried, and concentrated. Finally, the crude product was purified by a silica gel column using 5% methanol in methylene chloride as eluent, and the product was dried in a high vacuum pump to obtain indole-3-carboxylic acid-glycine conjugate **24** (280 mg, yield 56%) as a white solid. The structure of indole-3-carboxylic acid-glycine was confirmed by nuclear magnetic resonance (NMR) and mass spectra analysis.

Spectrum from 240812_TrpMetabolites_POS_184.wiff2 (sam...MSMS (CID) of 247.0 (50 - 270) from 7.063 to 7.090 min

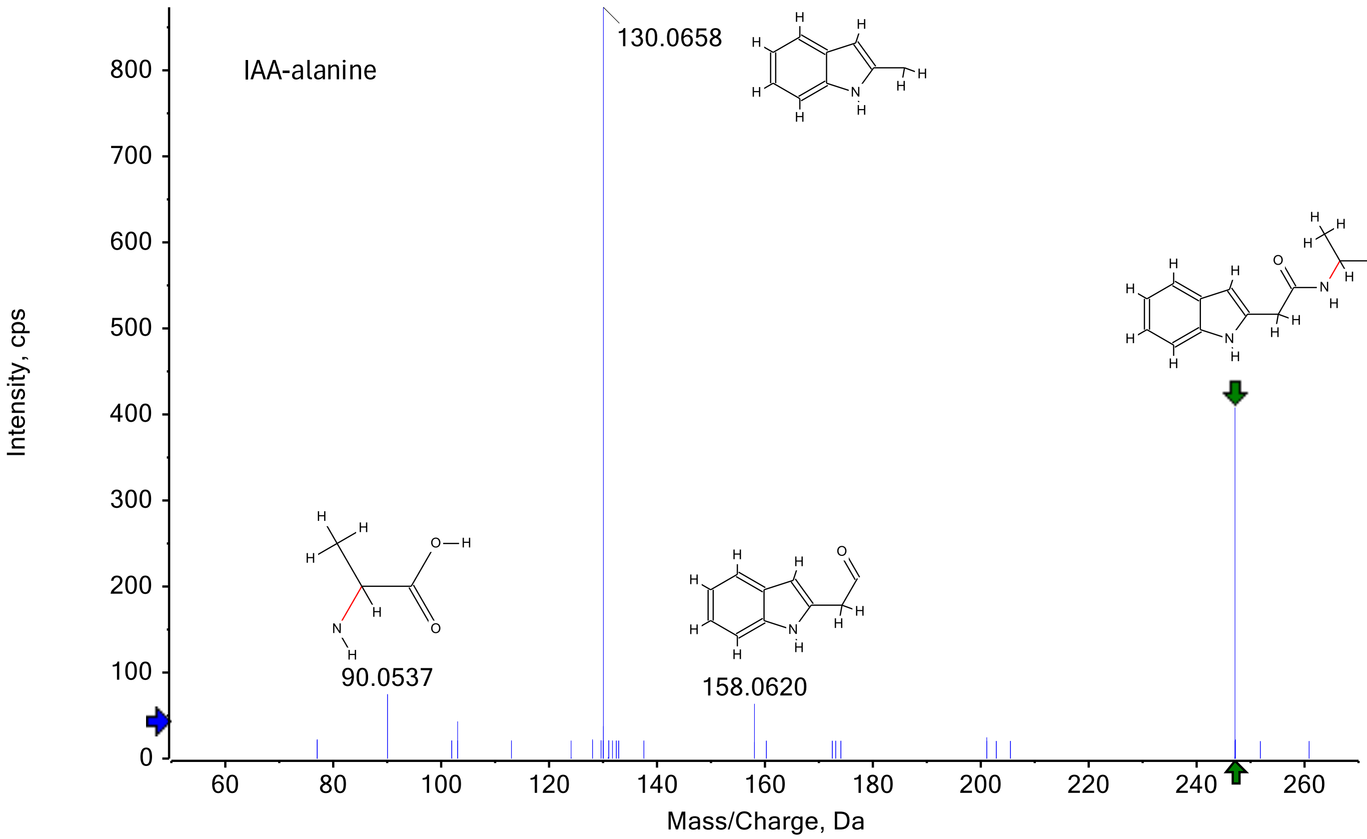

### IAA-glycine

Spectrum from Ethan_Trp_250131_45.wiff (sample 1) - InG 9, Experiment 23, +TOF MS^2 (CID) of 233.1 (50 - 250) from 5.843 to 5.994 min Precursor: 233.1 Da, CE: 45.0

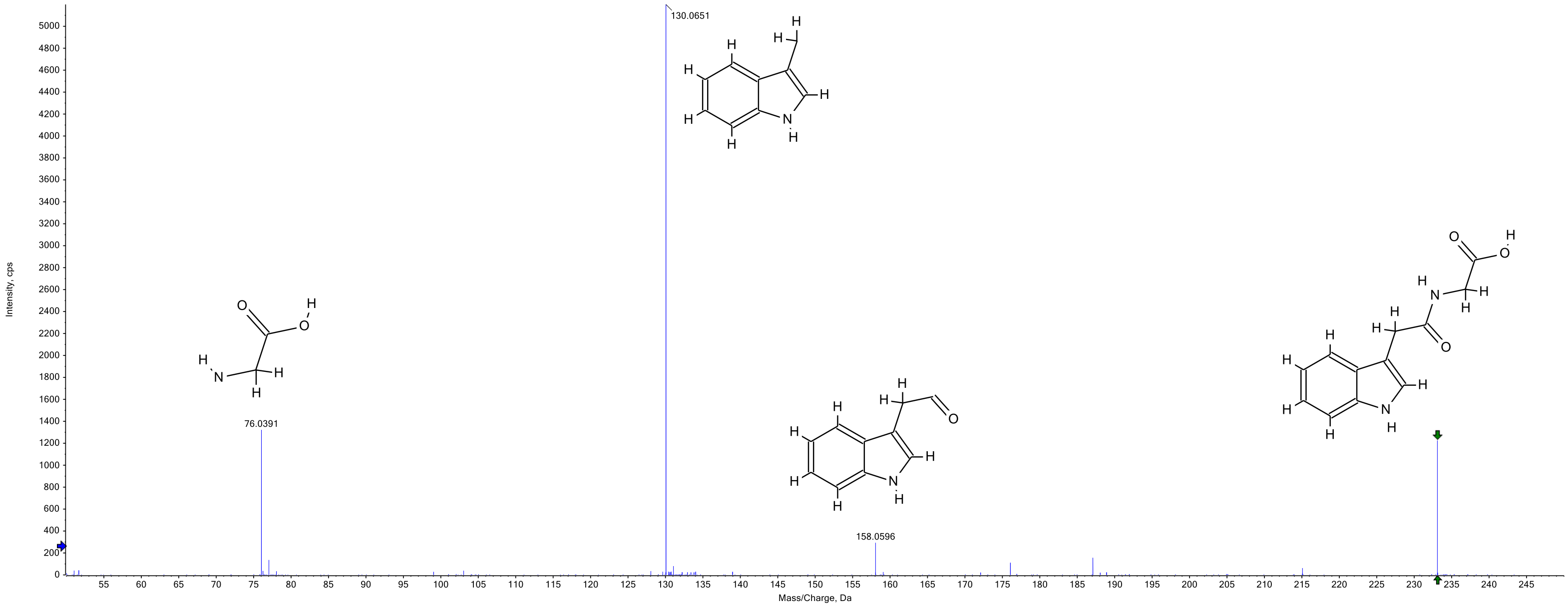

Spectrum from 240812_TrpMetabolites_POS_184.wiff2 (sampl...F MSMS (CID) of 305.1 (50 - 200) from 6.210 to 6.411 min

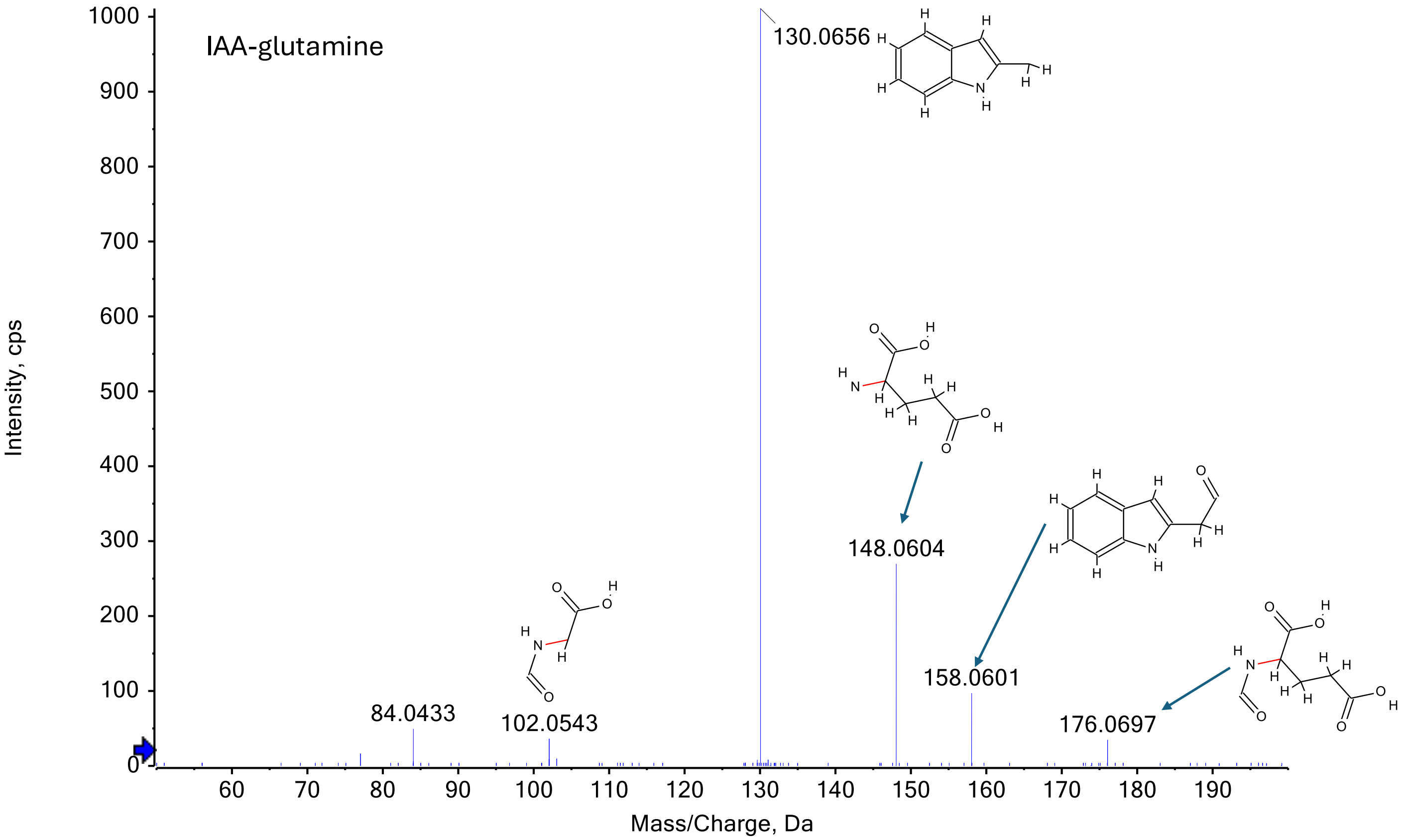

IAA-serine

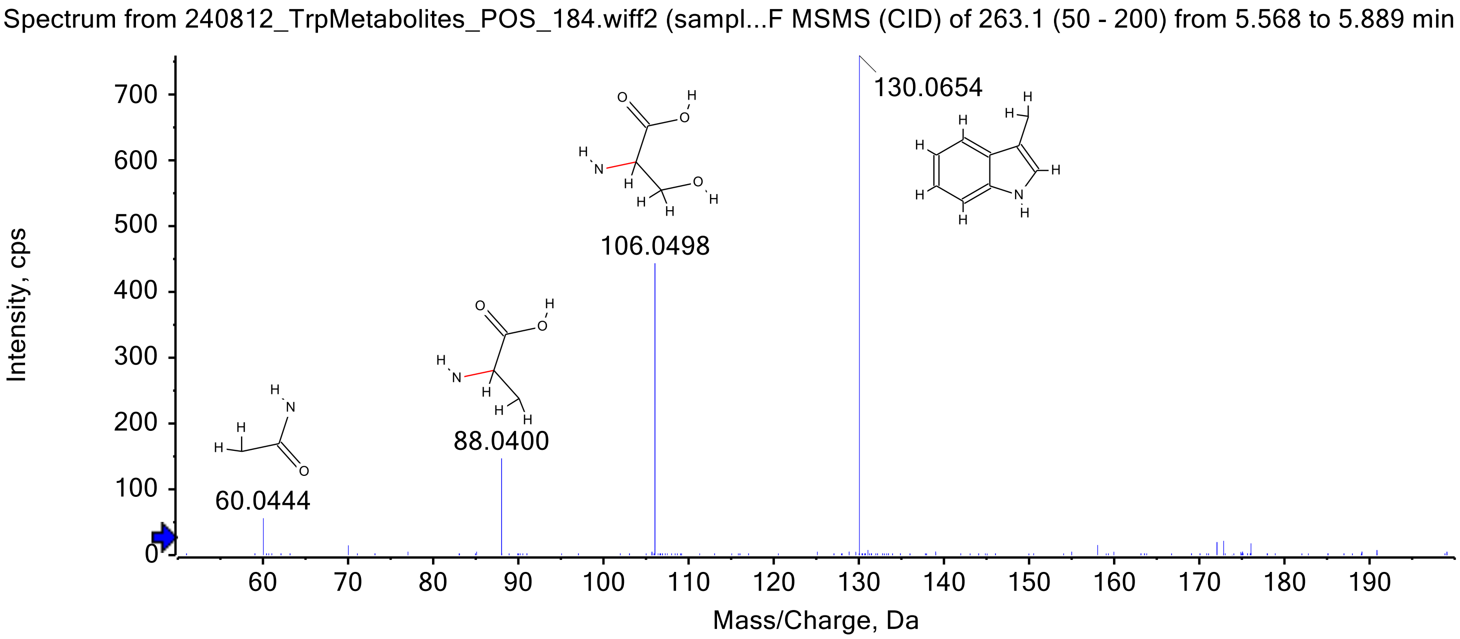
Spectrum from 240812_TrpMetabolites_POS_184.wiff2 (sampl...F MSMS (CID) of 263.1 (50 - 200) from 5.568 to 5.889 min
Intensity, cps
Mass/Charge, Da
130.0654
106.0498
88.0400
60.0444

### IAA-threonine

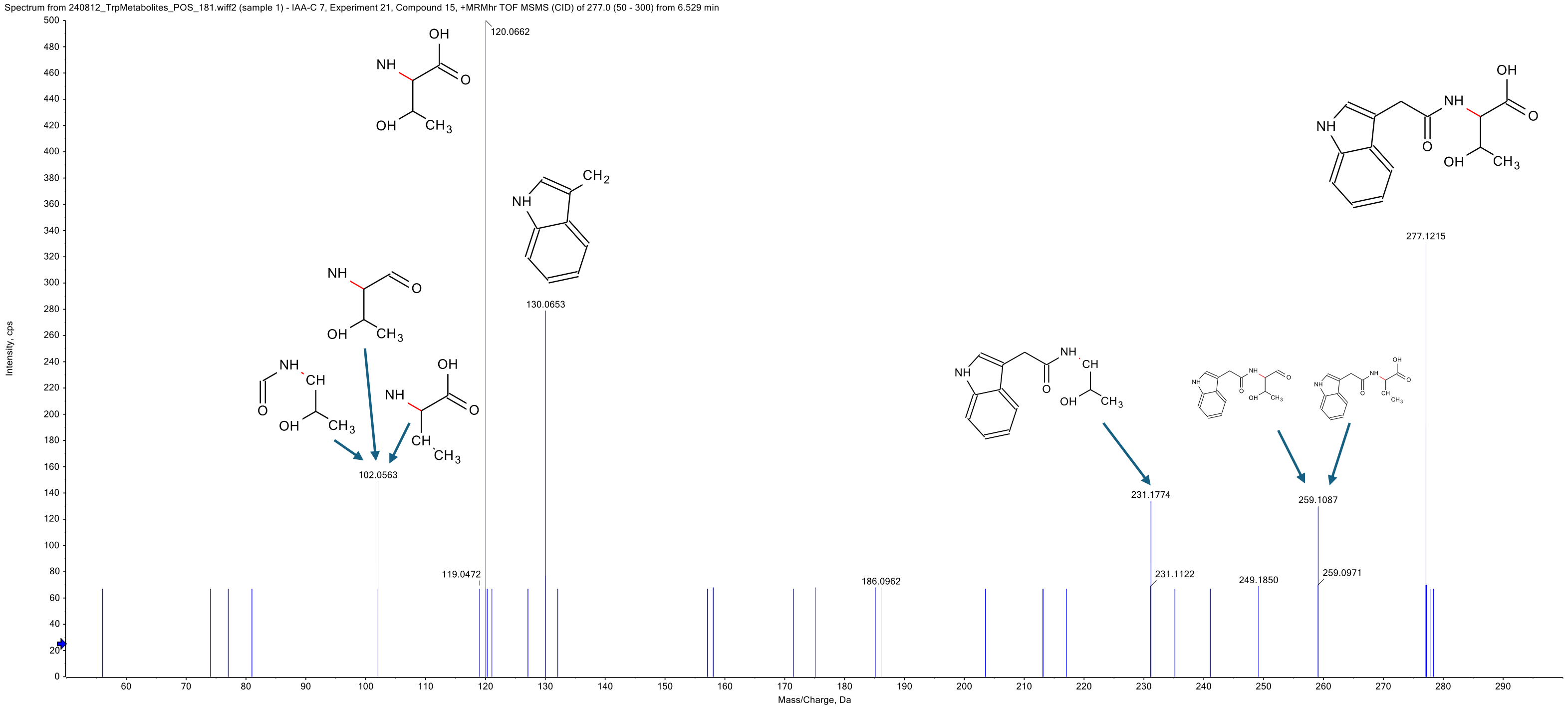

### IPA-glycine

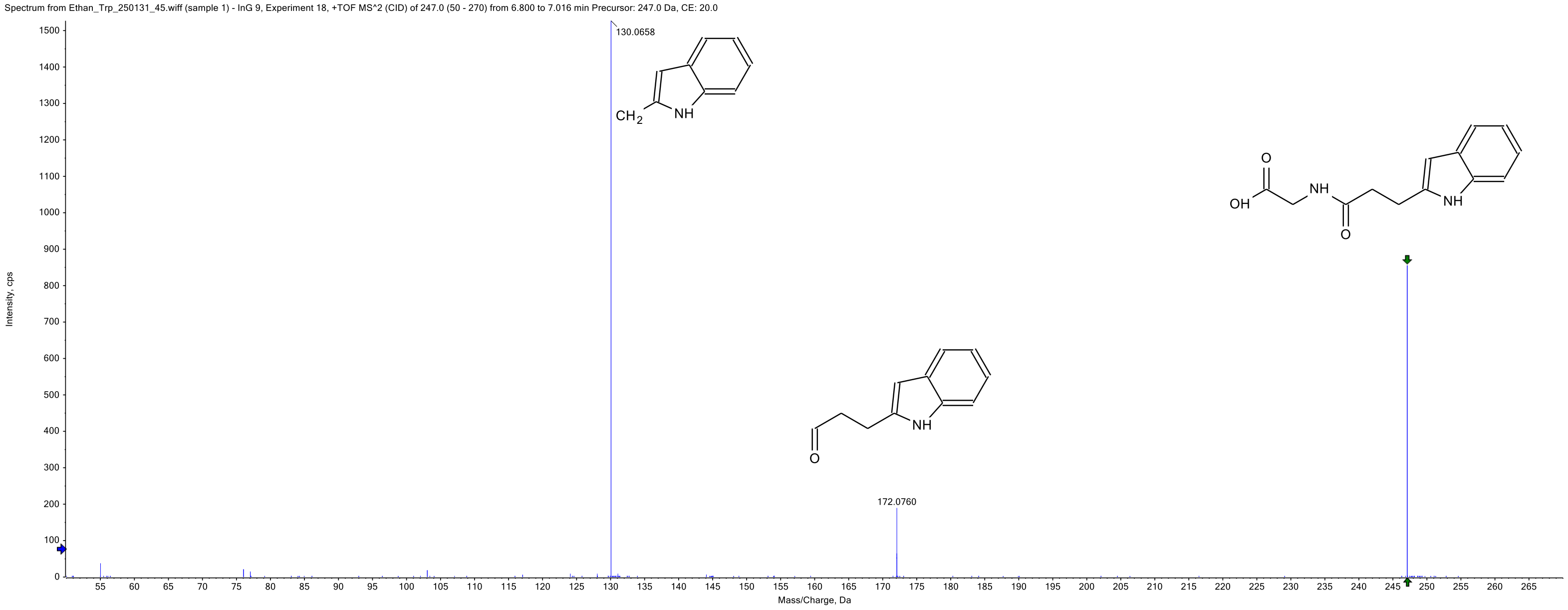

Spectrum from Ethan_Trp_250131_45.wiff (sample 1) - InG 9, Experiment 18, +TOF MS^2 (CID) of 247.0 (50 - 270) from 6.800 to 7.016 min Precursor: 247.0 Da, CE: 20.0
130.0658
CH2
NH
172.0760
O
NH
OH
NH
O
O
NH
Intensity, cps
Mass/Charge, Da

#### ICA-glycine

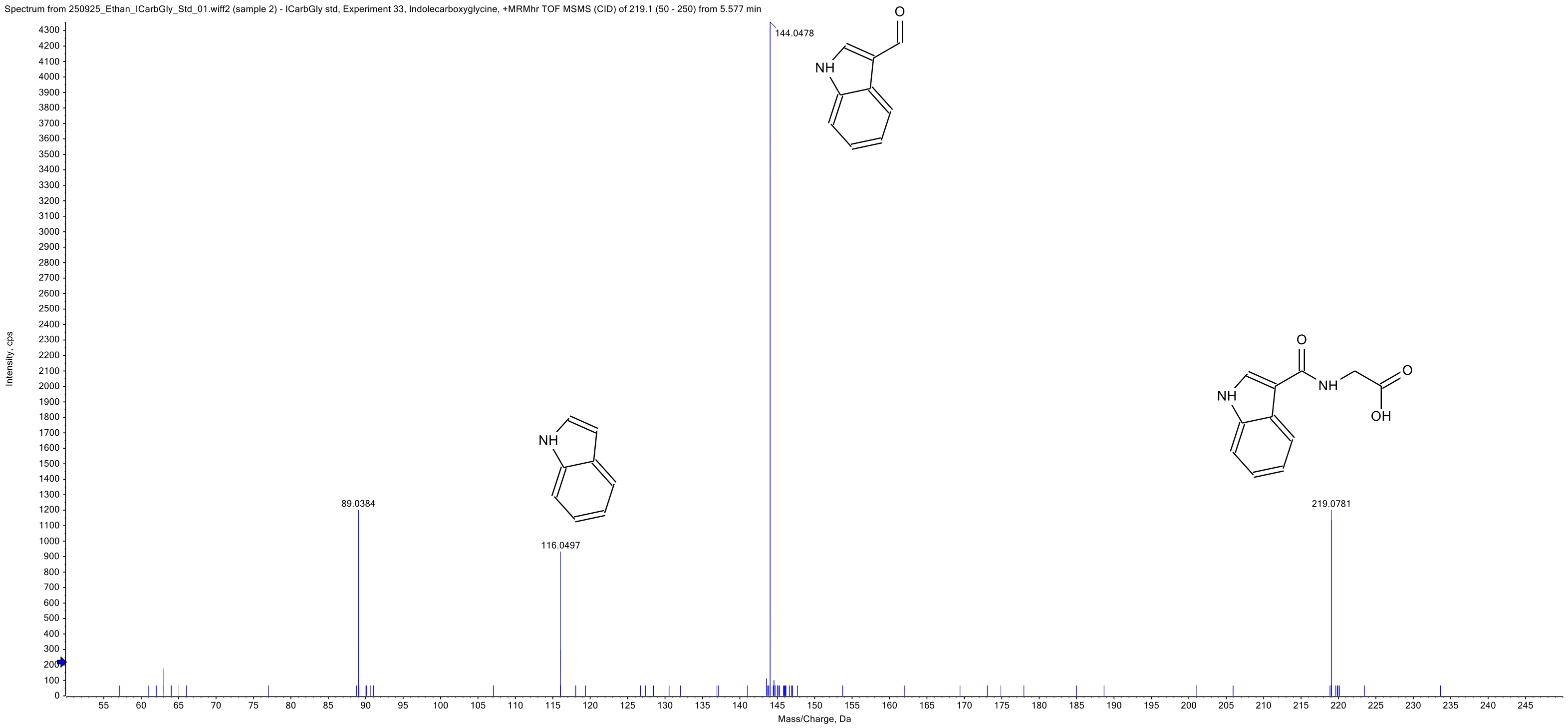
Spectrum from 250925_Ethan_ICarbGly_Std_01.wiff2 (sample 2) - ICarbGly std, Experiment 33, Indolecarboxyglycine, +MRMhr TOF MSMS (CID) of 219.1 (50 - 250) from 5.577 min
144.0478
89.0384
116.0497
219.0781
Intensity, cps
Mass/Charge, Da
NH
O
OH

ICA-glycine

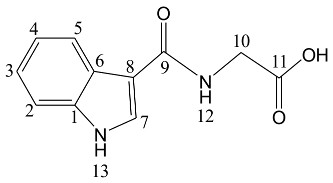


IPA-glycine

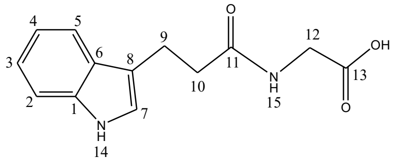


IAA-glycine

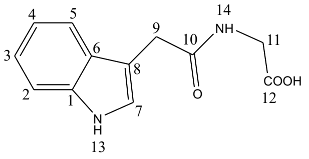


IAA-serine

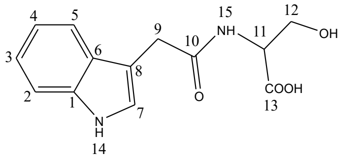


IAA-alanine

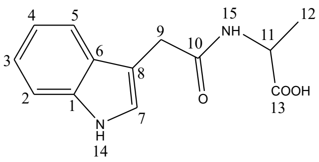


IAA-glutamine

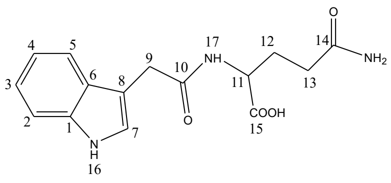


IAA-threonine

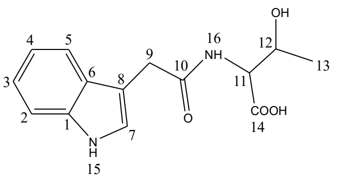

#### The summary of the chemical shift (ppm) of $^1H$ and $^{13}C$ of the ICA-, IPA-, and IAA-combined compounds

| | ICA-glycine | | IPA-glycine | | IAA-glycine | | IAA-serine | | IAA-alanine | | IAA-glutamine | | IAA-threonine | |
|---|---|---|---|---|---|---|---|---|---|---|---|---|---|---|
| Atom | $^1H$(ppm) | $^{13}C$(ppm) | $^1H$(ppm) | $^{13}C$(ppm) | $^1H$(ppm) | $^{13}C$(ppm) | $^1H$(ppm) | $^{13}C$(ppm) | $^1H$(ppm) | $^{13}C$(ppm) | $^1H$(ppm) | $^{13}C$(ppm) | $^1H$(ppm) | $^{13}C$(ppm) |
| 1 | - | 136.22 | - | 136.16 | - | 136.08 | - | 136.17 | - | 135.98 | - | 136.11 | - | 136.18 |
| 2 | 7.43 | 111.68 | 7.32 | 111.08 | 7.33 | 110.96 | 7.34 | 111.02 | 7.33 | 111.01 | 7.33 | 111.05 | 7.33 | 110.98 |
| 3 | 7.15 | 126.06 | 7.05 | 120.70 | 7.06 | 120.61 | 7.06 | 120.71 | 7.06 | 120.64 | 7.05 | 120.64 | 7.06 | 120.62 |
| 4 | 7.10 | 120.30 | 6.97 | 117.97 | 6.96 | 118.01 | 6.96 | 118.06 | 6.96 | 117.97 | 6.95 | 118.03 | 6.95 | 117.96 |
| 5 | 8.11 | 120.70 | 7.52 | 118.10 | 7.54 | 118.40 | 7.51 | 118.34 | 7.55 | 118.52 | 7.51 | 118.35 | 7.52 | 118.35 |
| 6 | - | 121.78 | - | 126.93 | - | 127.24 | - | 127.26 | - | 127.26 | - | 127.18 | - | 127.23 |
| 7 | 8.03 | 128.00 | 7.11 | 121.98 | 7.22 | 123.59 | 7.23 | 123.80 | 7.19 | 123.41 | 7.21 | 123.67 | 7.23 | 123.68 |
| 8 | - | 110.08 | - | 113.76 | - | 108.62 | - | 108.66 | - | 108.79 | - | 108.83 | - | 108.88 |
| 9 | - | 164.70 | 2.91 | 20.67 | 3.55 | 32.16 | 3.55 | 32.59 | 3.54 | 32.02 | 3.53 | 32.6 | 3.58 | 32.44 |
| 10 | 3.92 | 40.45 | 2.50 | 35.74 | - | 170.69 | - | 170.34 | - | 170.4 | - | 169.59 | - | 170.18 |
| 11 | - | 172.10 | - | 172.33 | 3.62 | 41.63 | 3.72 | 54.33 | 4.20 | 47.31 | 3.99 | 52.76 | 4.07 | 56.75 |
| 12 | 8.21 | - | 3.74 | 40.46 | - | 171.33 | 3.54, 3.26 | 62.52 | 1.27 | 17.08 | 1.84, 1.67 | 28.56 | 3.93 | 65.89 |
| 13 | 11.57 | - | - | 171.44 | 10.89 | - | - | 173.07 | - | 174.29 | 2.12, 2.04 | 32.65 | 0.87 | 18.26 |
| 14 | - | - | 10.75 | - | 7.8 | - | 10.95 | - | 10.85 | - | - | 176.66 | - | 172.60 |
| 15 | - | - | 8.20 | - | - | - | 7.29 | - | 8.25 | - | - | 174.14 | 10.91 | - |
| 16 | - | - | - | - | - | - | - | - | - | - | 10.96 | - | 7.36 | - |
| 17 | - | - | - | - | - | - | - | - | - | - | 7.45 | - | - | - |

Assignment reference: (DMSO-d6, $^1H$ of the residual protons: δ 2.50; $^{13}C$: δ 39.5)
$^1H$-NMR (600 MHz, DMSO-d6, 25 ºC) and $^{13}C$-NMR (150 MHz, DMSO-d6, 25 ºC)

ICA-glycine

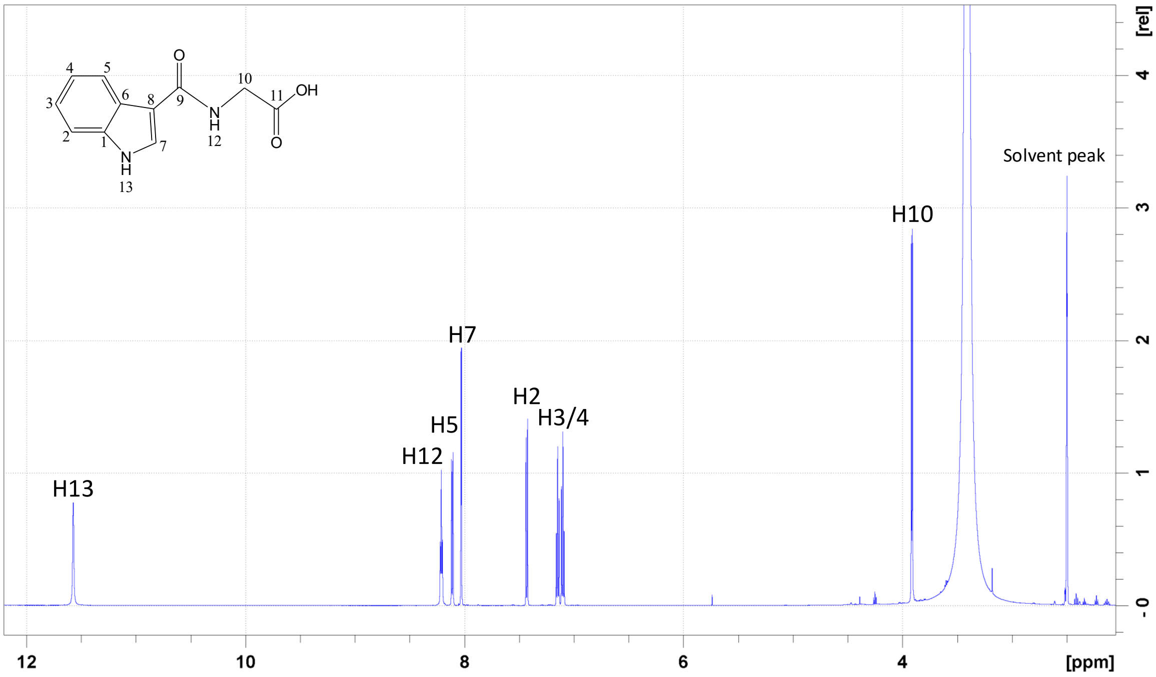


1D $^{1}H$ spectrum of ICA-glycine with assignments (in DMSO-d6)

ICA-glycine

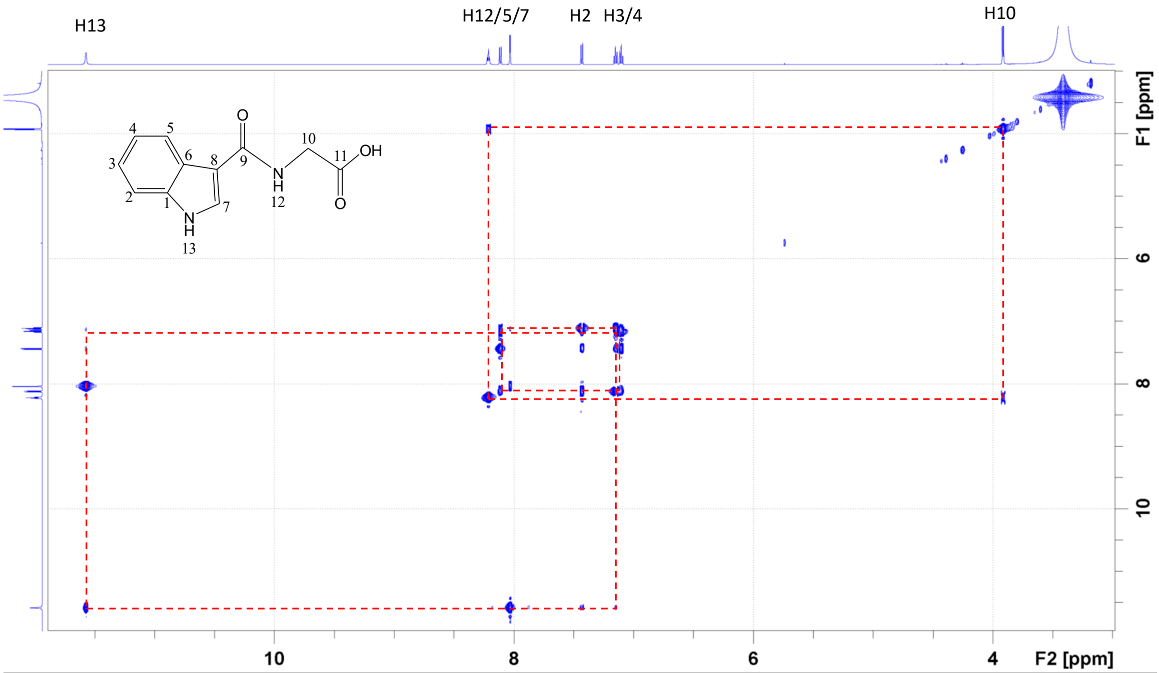


$^1$H-$^1$H TOCSY spectrum of ICA-glycine with assignments (in DMSO-d6)

ICA-glycine

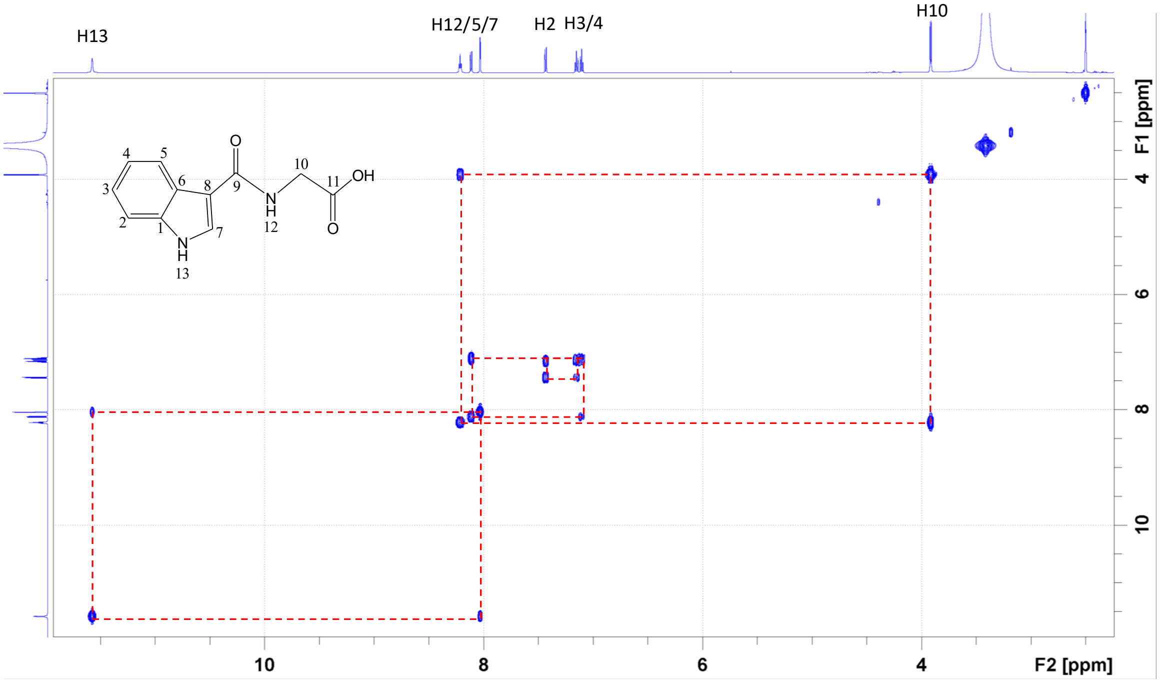


$^{1}H$-$^{1}H$ COSY spectrum of ICA-glycine with assignments (in DMSO-d6)

ICA-glycine

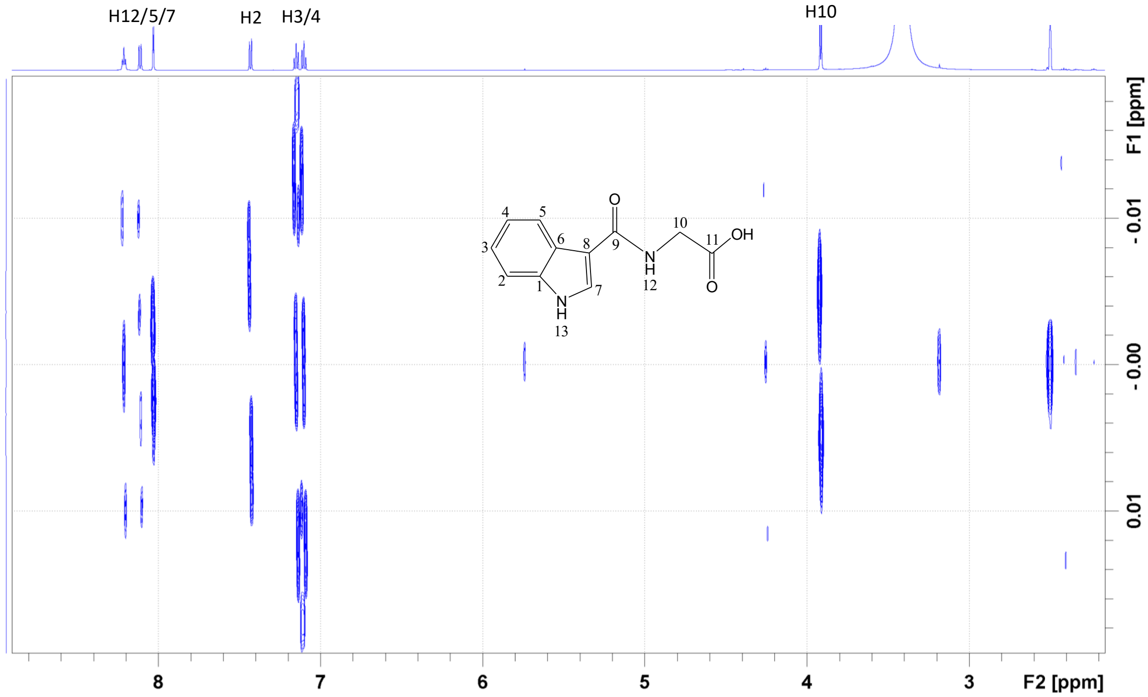


$^1H$-$^1H$ JREs spectrum of ICA-glycine showing J-coupling patterns (in DMSO-d6)

ICA-glycine

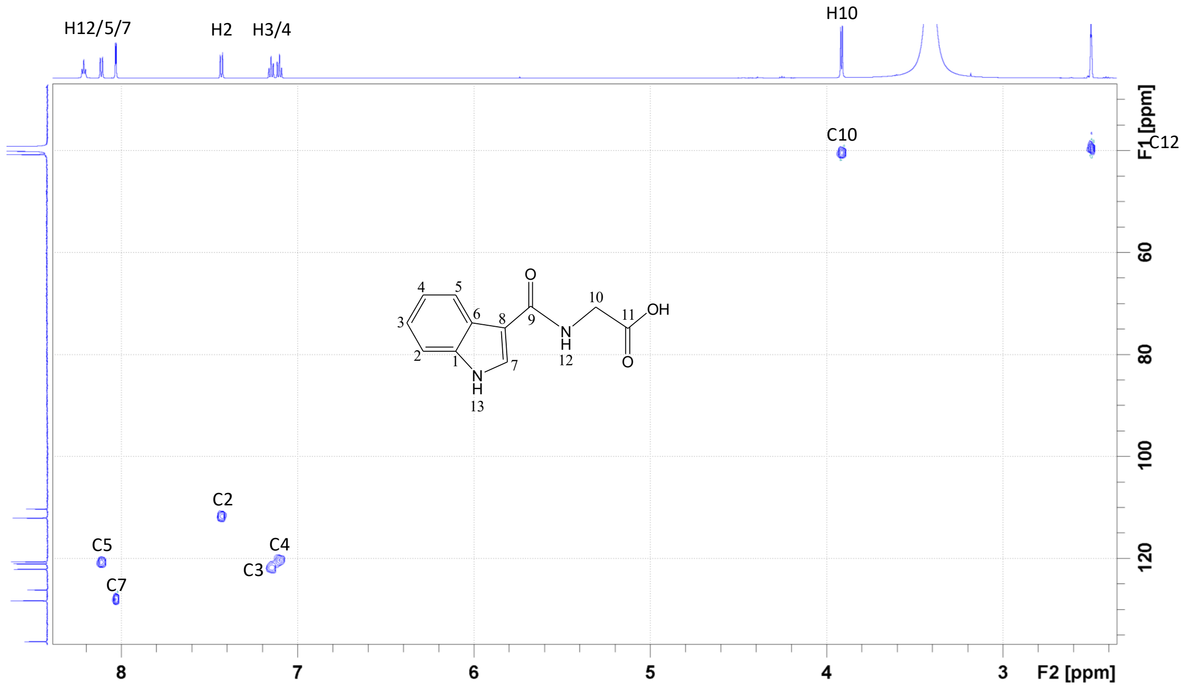


$^{1}H$-$^{13}C$ HSQC spectrum of ICA-glycine with assignments (in DMSO-d6)

ICA-glycine

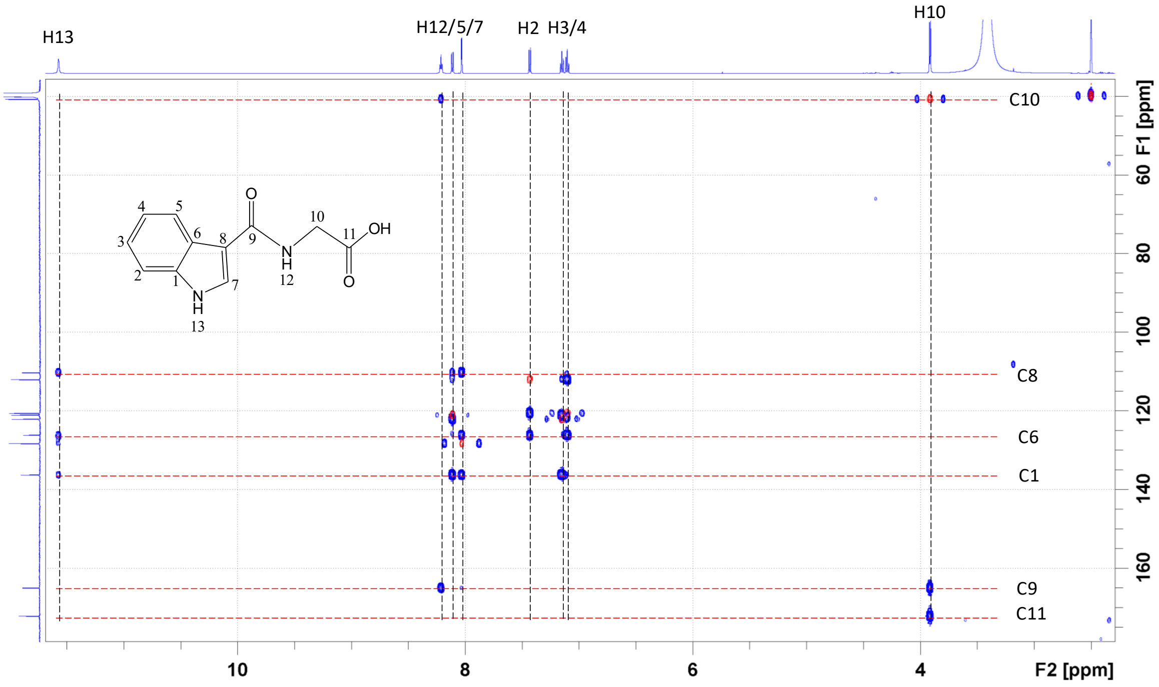


$^{1}H$-$^{13}C$ HMBC (blue) and HSQC (red) spectrum of ICA-glycine and signal assignments (in DMSO-d6)

IPA-glycine

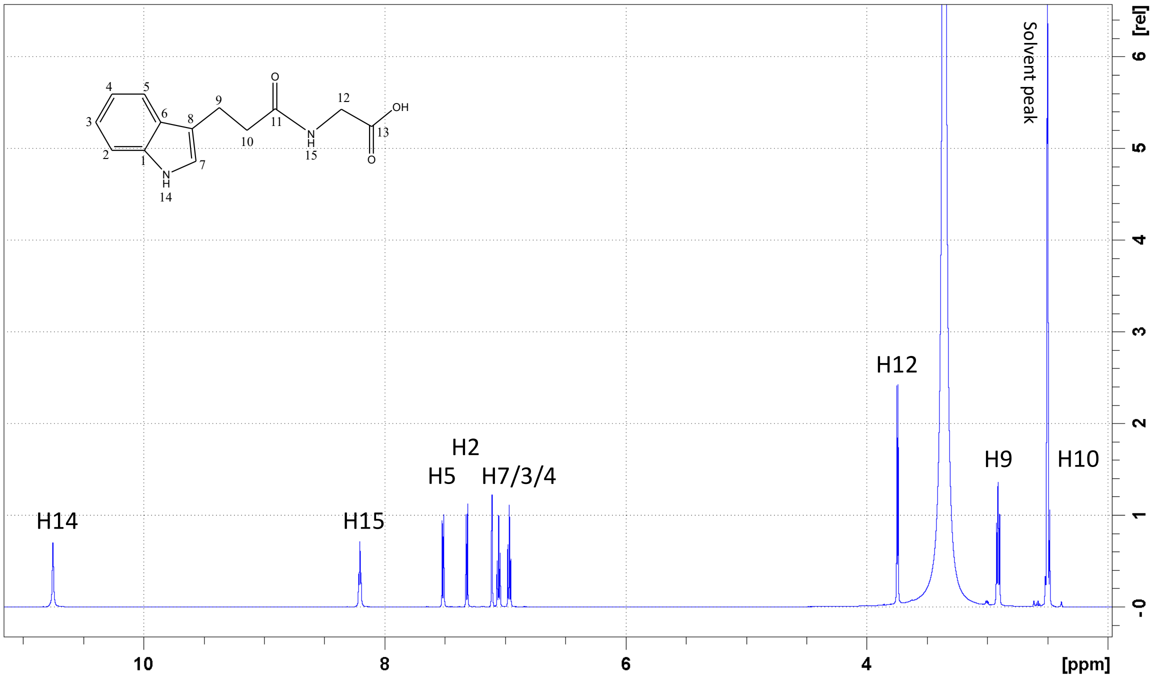


1D ¹H spectrum of IPA-glycine with assignments (in DMSO-d6)

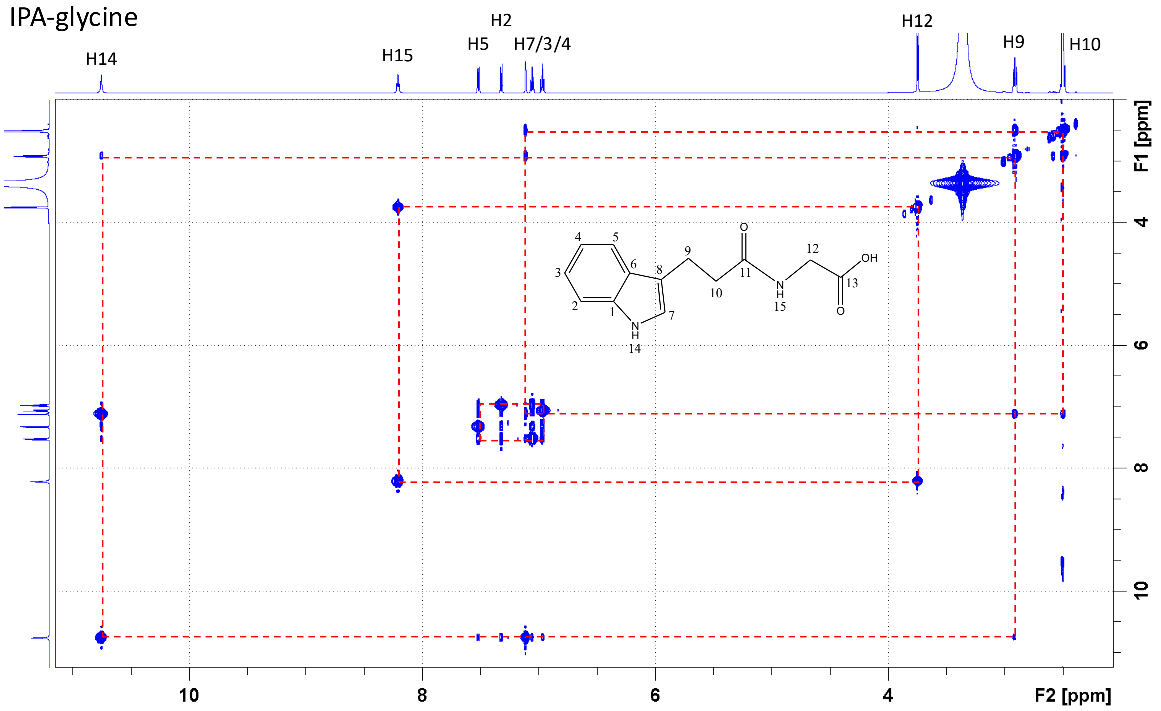


$^1$H-$^1$H TOCSY spectrum of IPA-glycine with assignments (in DMSO-d6)

IPA-glycine

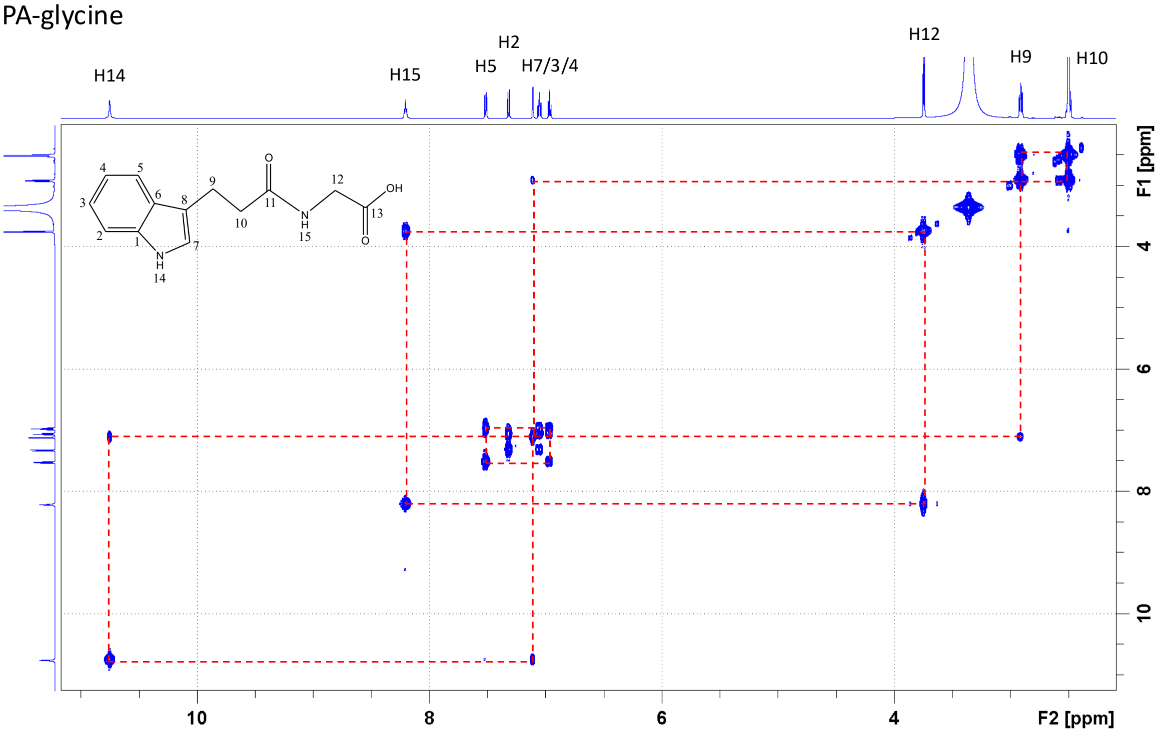


$^1H$-$^1H$ COSY spectrum of IPA-glycine with assignments (in DMSO-d6)

IPA-glycine

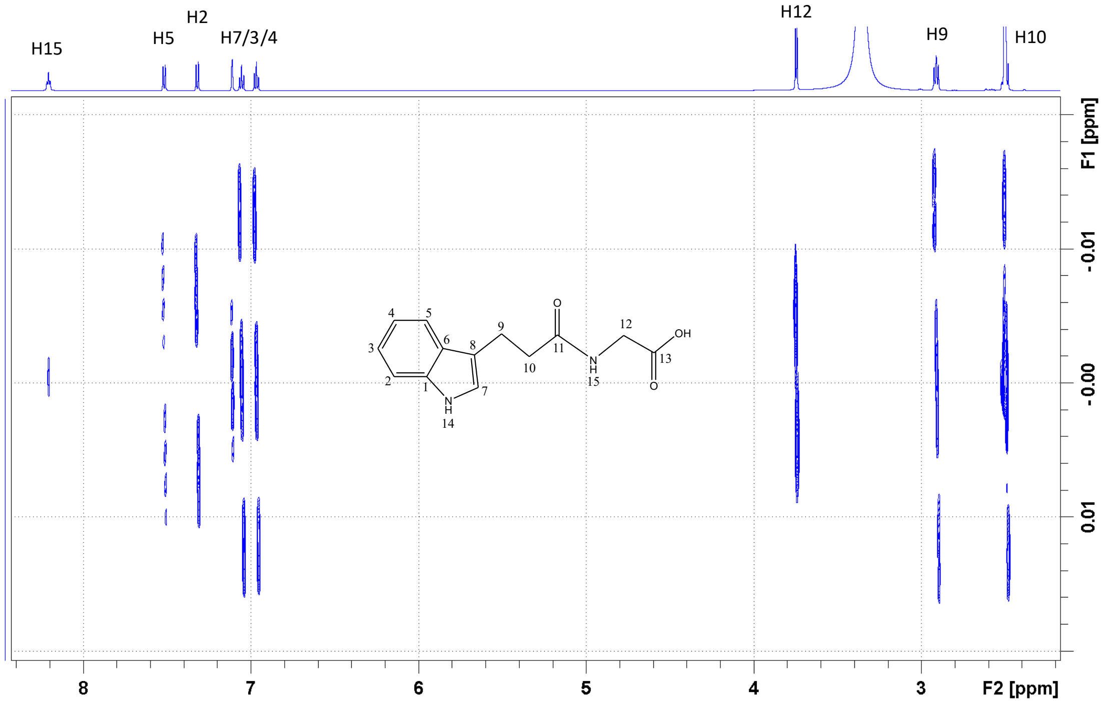


$^{1}H$-$^{1}H$ JREs spectrum of IPA-glycine showing J-coupling patterns (in DMSO-d6)

IPA-glycine

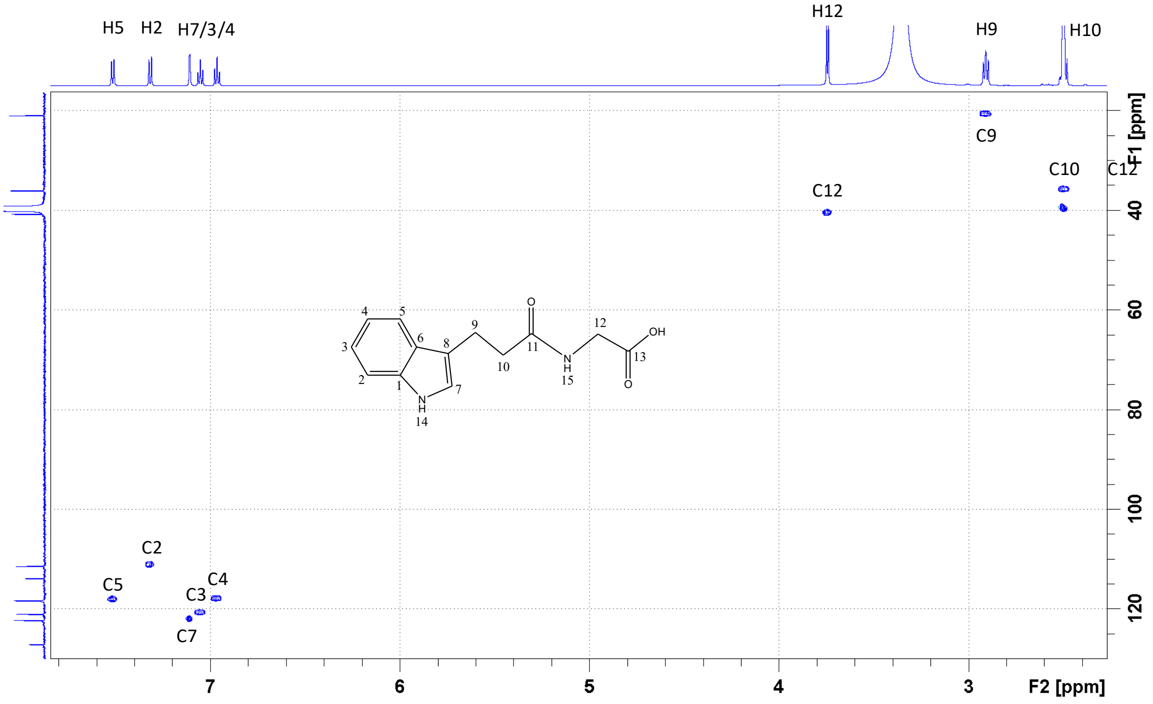


$^{1}H$-$^{13}C$ HSQC spectrum of IPA-glycine with assignments (in DMSO-d6)

IPA-glycine

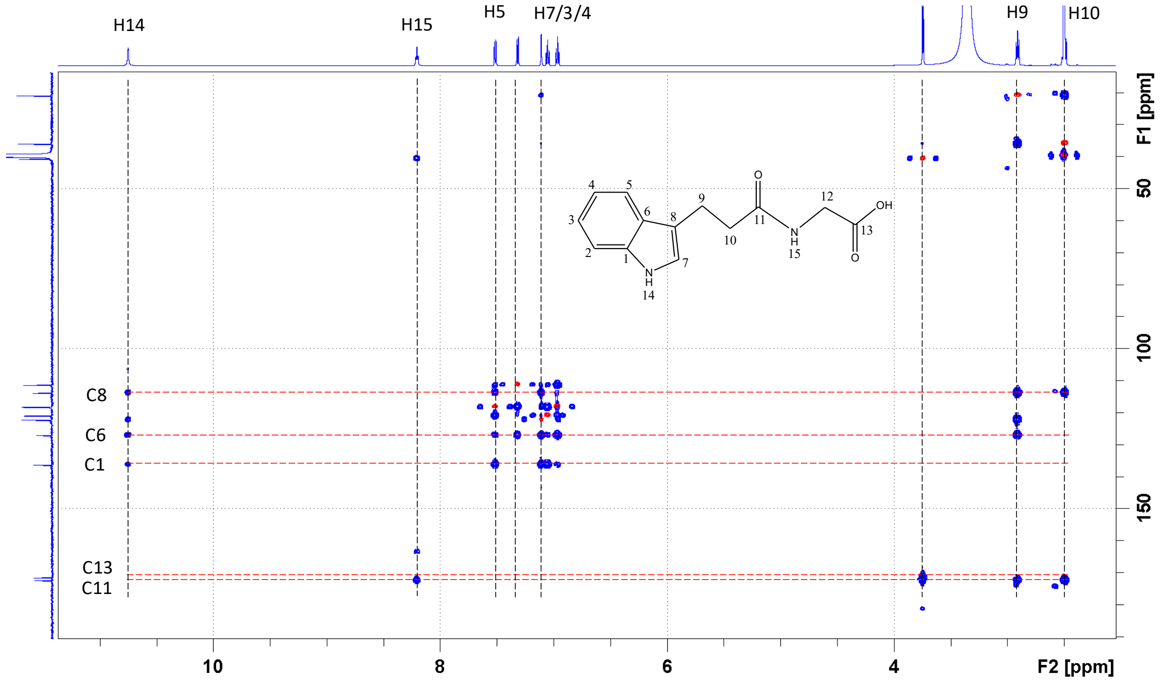


$^{1}H$-$^{13}C$ HMBC (blue) and HSQC (red) spectrum of IPA-glycine and signal assignments (in DMSO-d6)

IAA-glycine

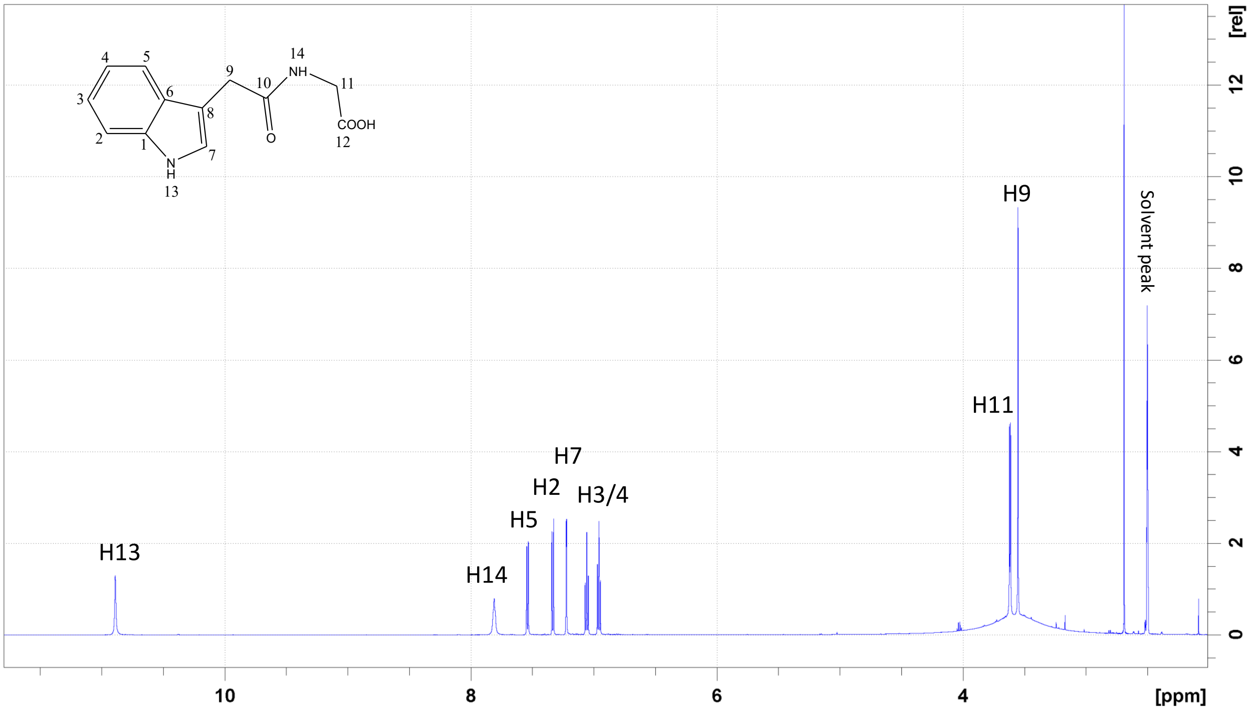


1D $^{1}H$ spectrum of IAA-glycine with assignments (in DMSO-d6)

IAA-glycine

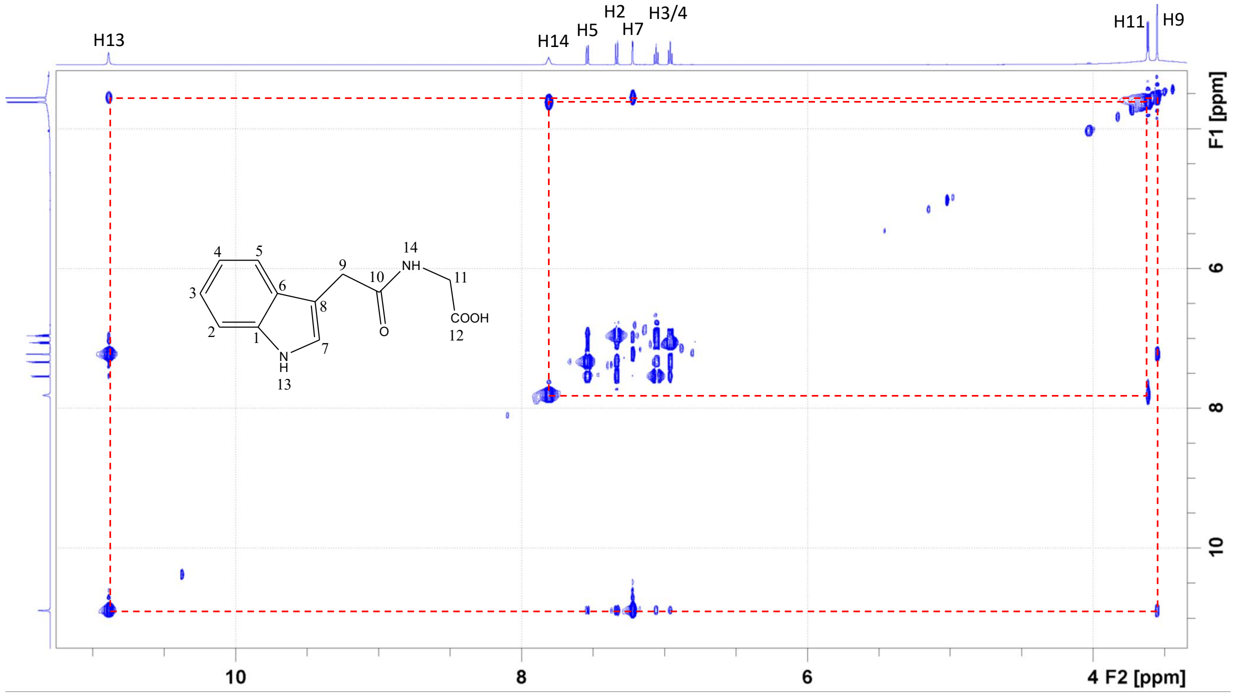


$^1H$-$^1H$ TOCSY spectrum of IAA-glycine with signal assignments (in DMSO-d6)

IAA-glycine

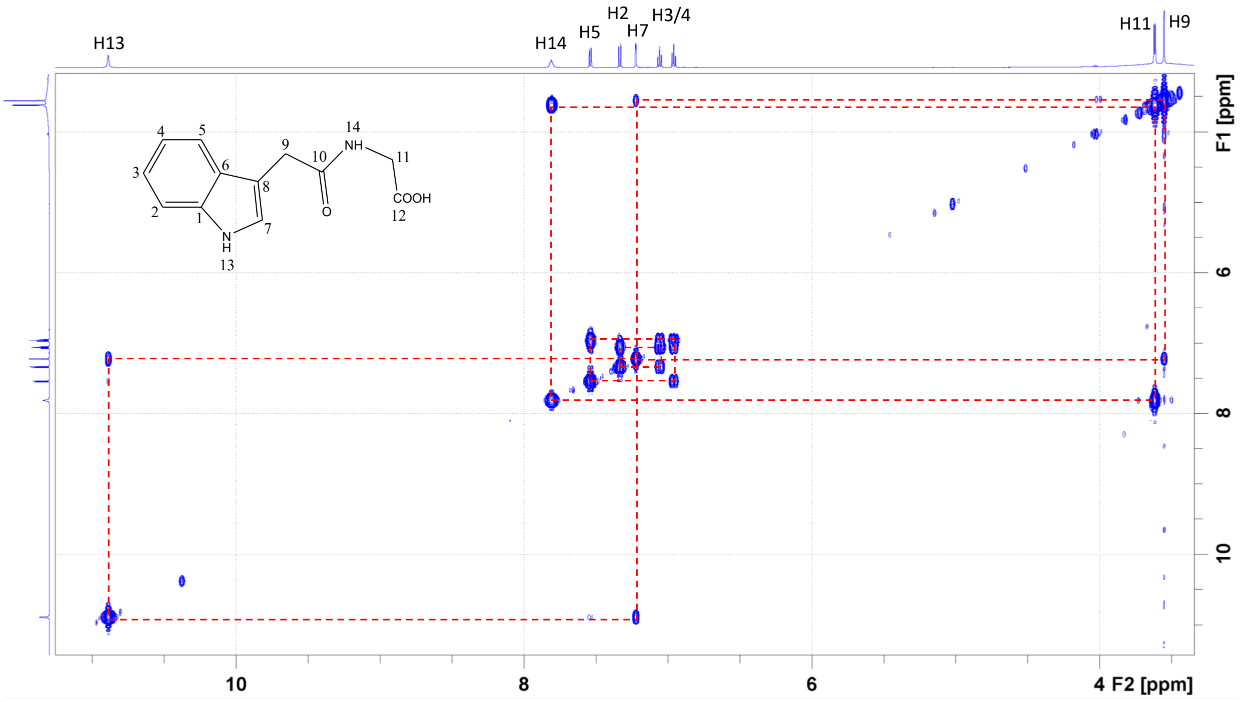


$^1H$-$^1H$ COSY spectrum of IAA-glycine with signal assignments (in DMSO-d6)

IAA-glycine

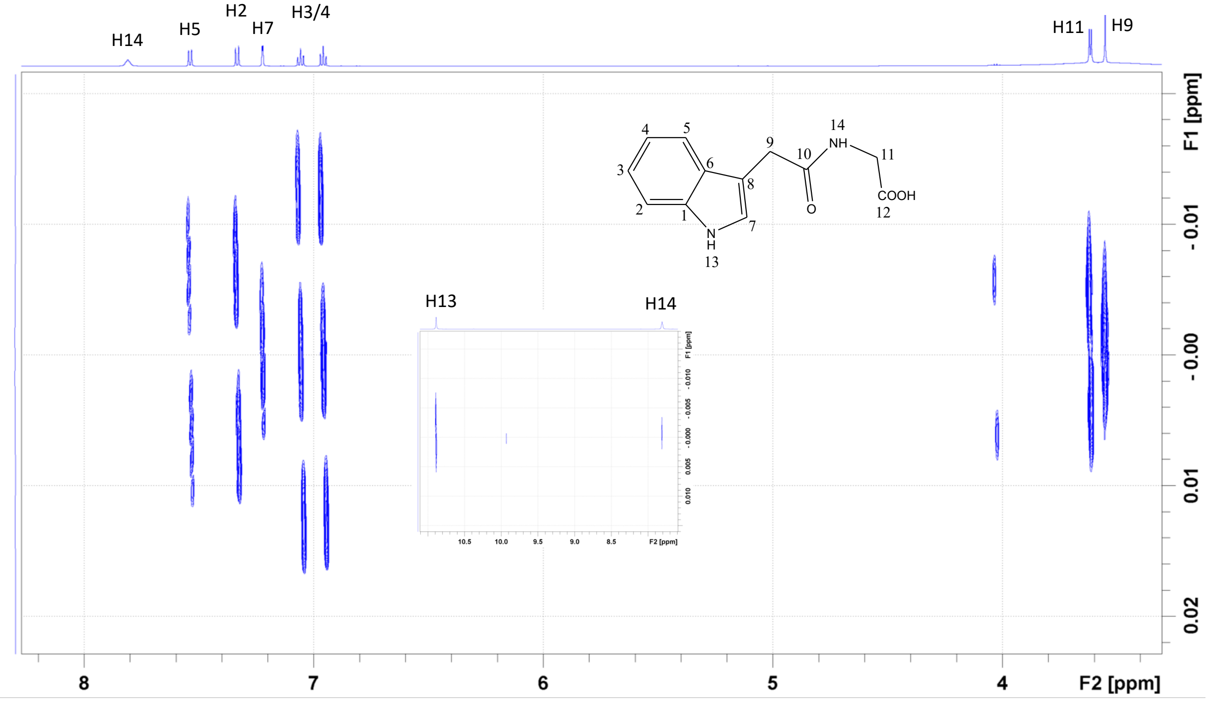


$^{1}H$-$^{1}H$ JREs spectrum of IAA-glycine showing J-coupling patterns (in DMSO-d6)

IAA-glycine

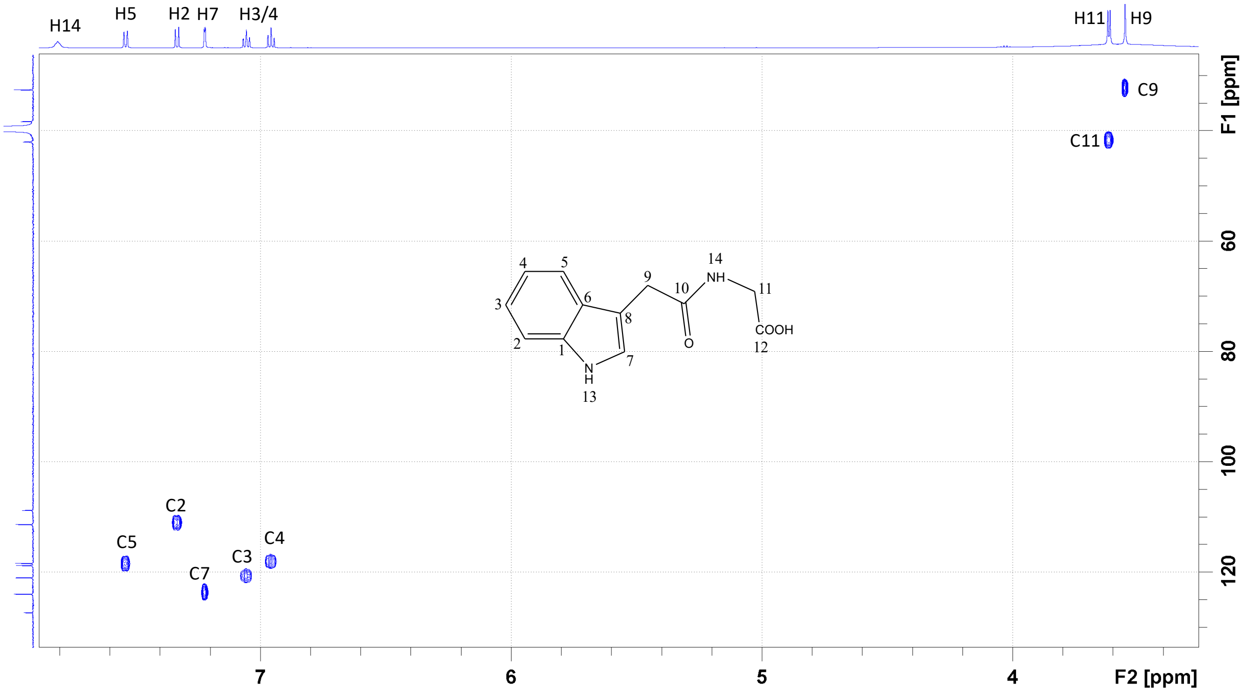


$^{1}H$-$^{13}C$ HSQC spectrum of IAA-glycine and signal assignments (in DMSO-d6)

IAA-glycine

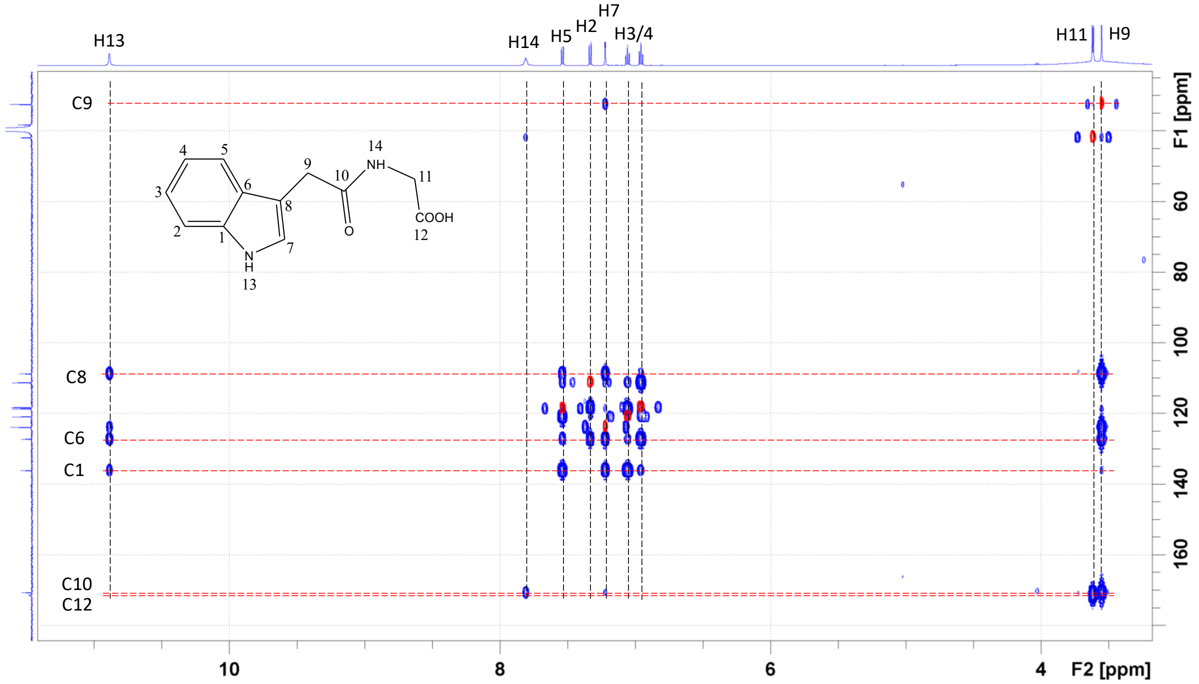


$^{1}H$-$^{13}C$ HMBC (blue) and HSQC (red) spectrum of IAA-glycine and signal assignments (in DMSO-d6)

IAA-serine

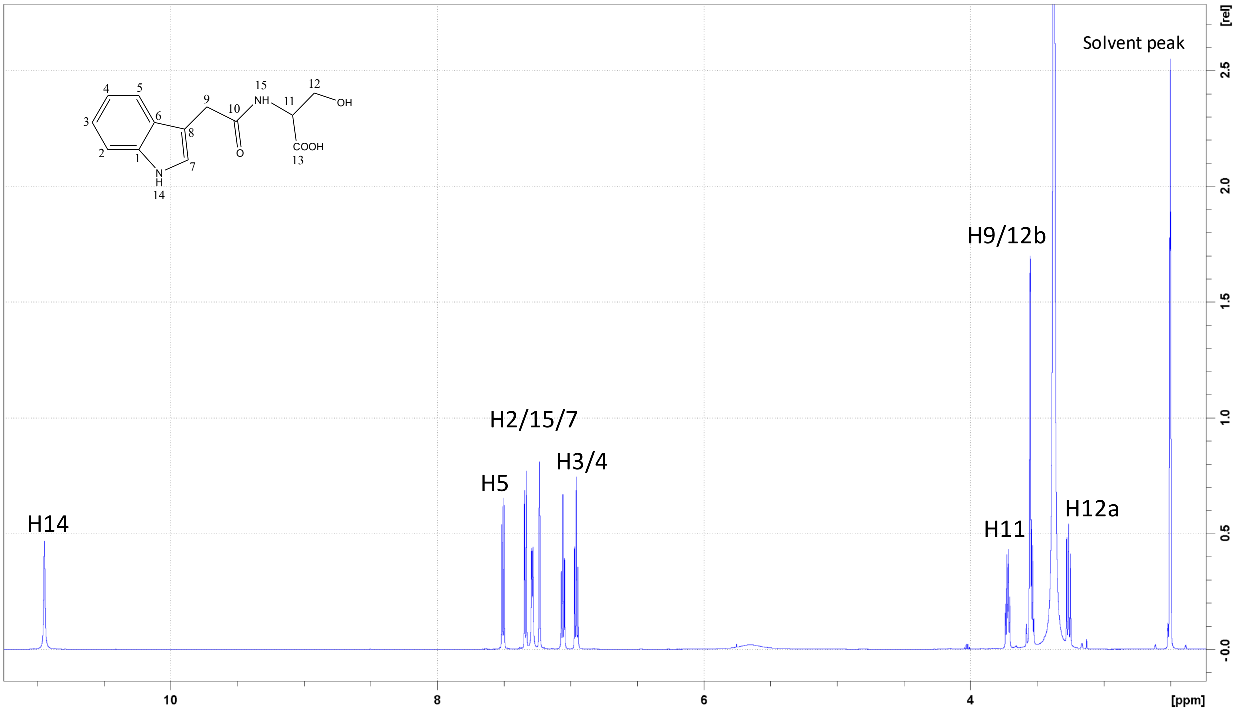


1D $^{1}H$ spectrum of IAA-serine with assignments (in DMSO-d6)

IAA-serine

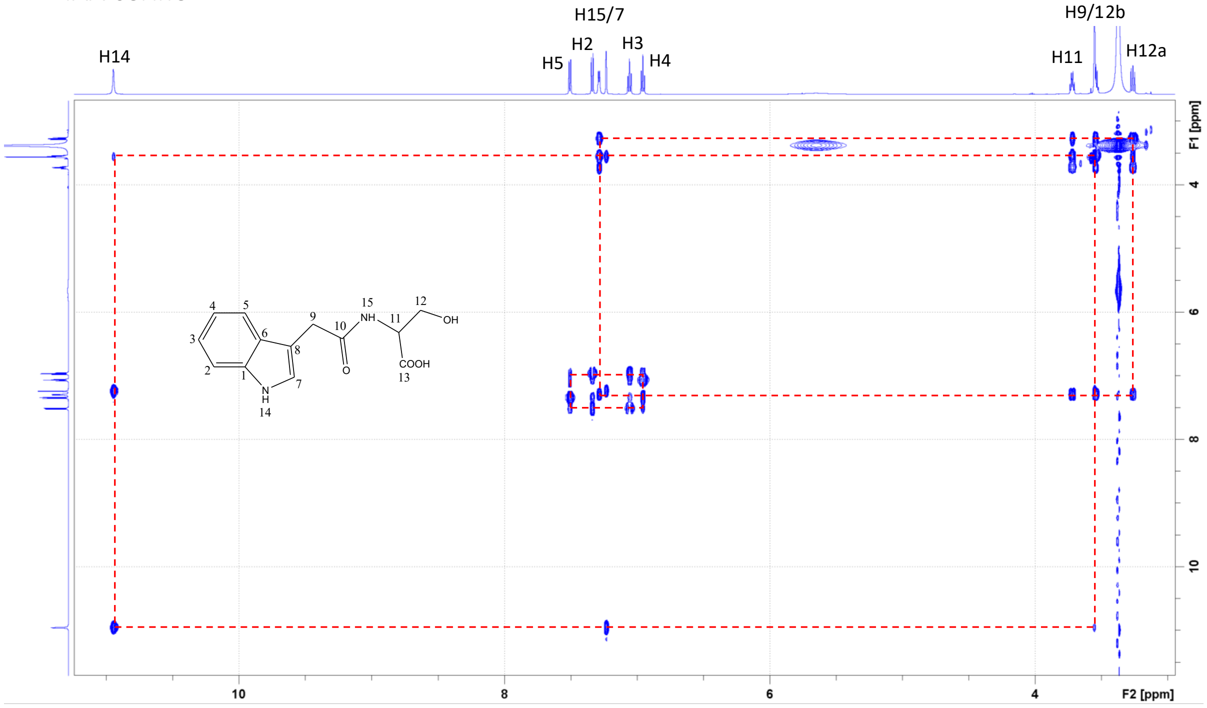


$^1$H-$^1$H TOCSY spectrum of IAA-serine with signal assignments (in DMSO-d6)

IAA-serine

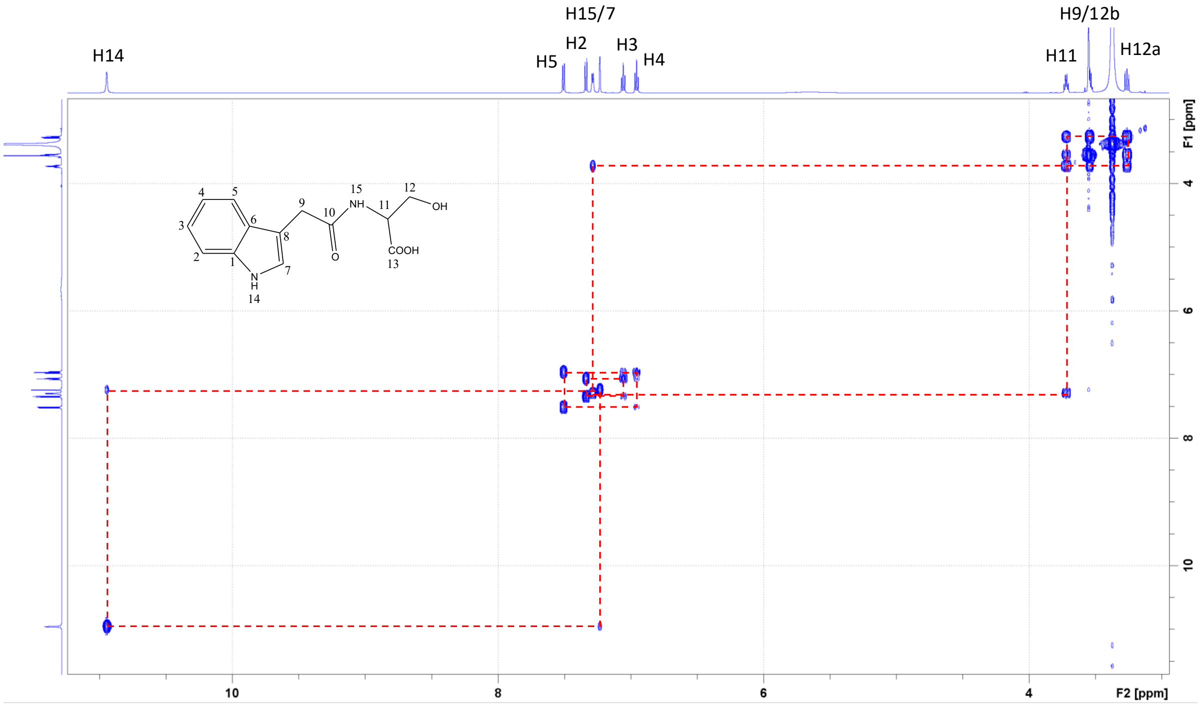


$^{1}H$-$^{1}H$ COSY spectrum of IAA-serine with signal assignments (in DMSO-d6)

IAA-serine

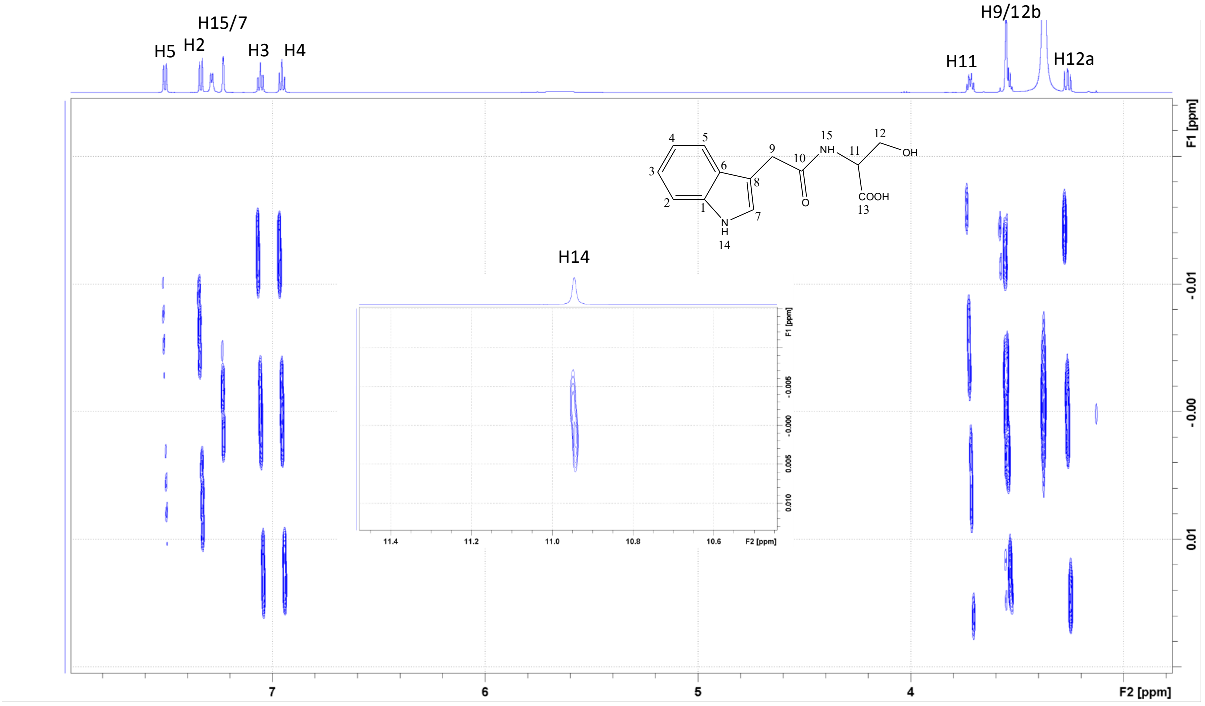


$^1$H-$^1$H JREs spectrum of IAA-serine showing J-coupling patterns (in DMSO-d6)

IAA-serine

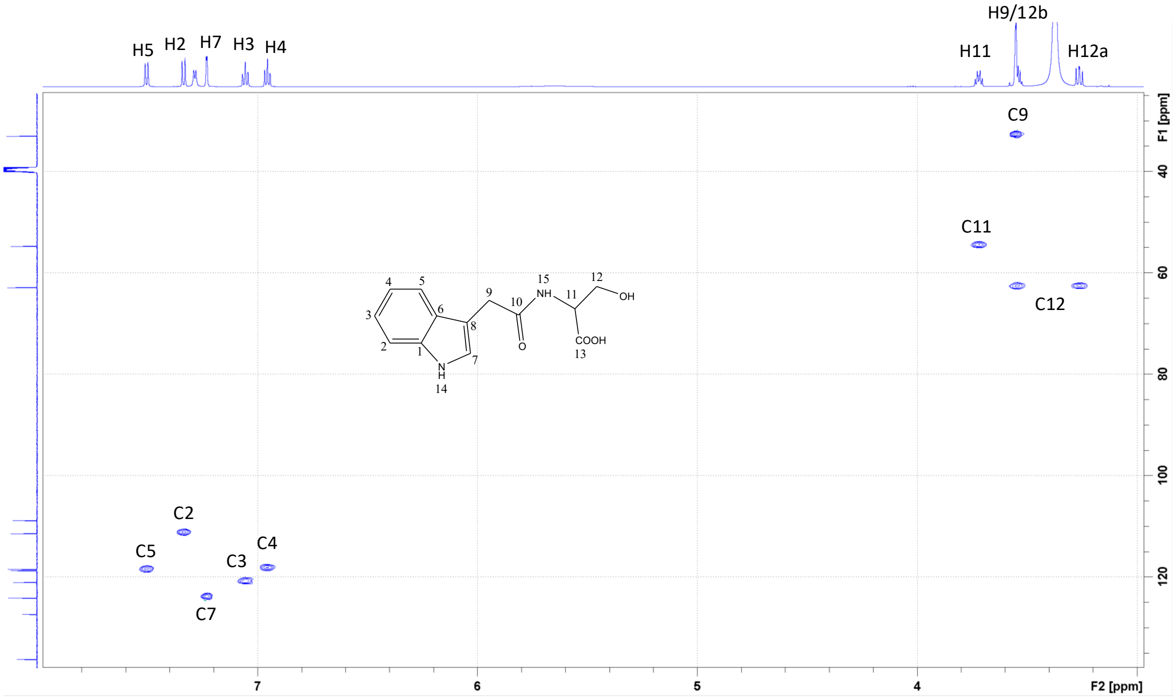


$^{1}H$-$^{13}C$ HSQC spectrum of IAA-serine and signal assignments (in DMSO-d6)

IAA-serine

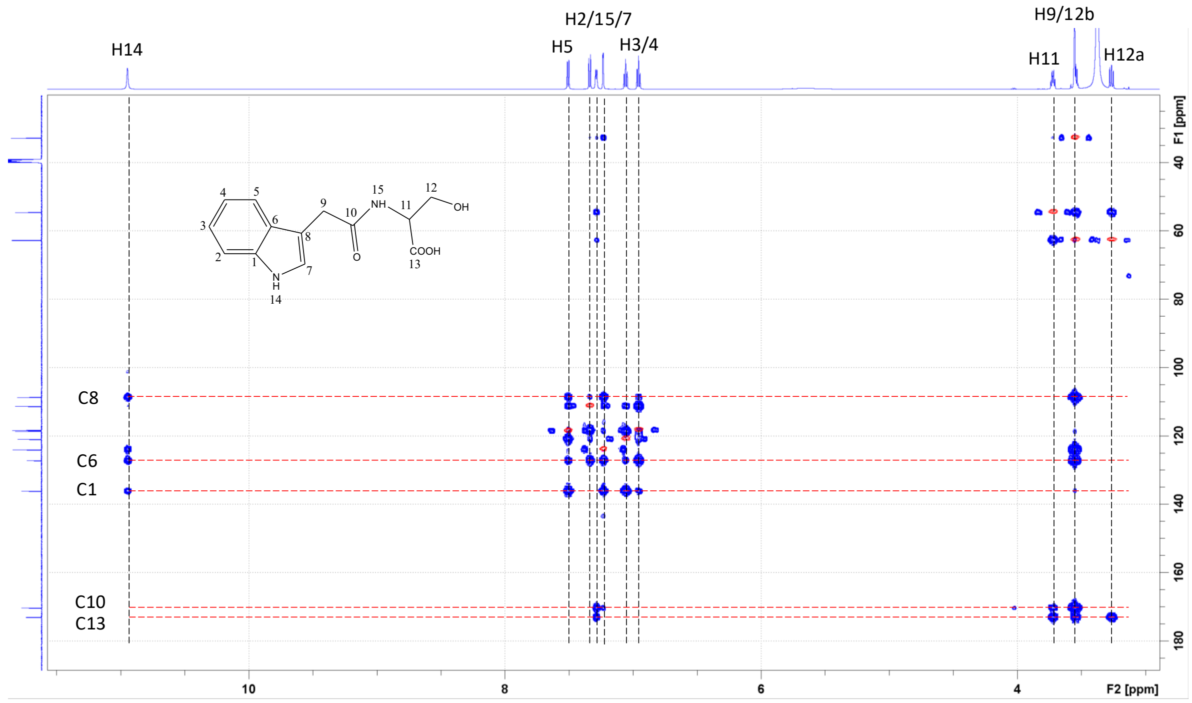


$^1H$-$^{13}C$ HMBC (blue) and HSQC (red) spectrum of IAA-serine and signal assignments (in DMSO-d6)

IAA-alanine

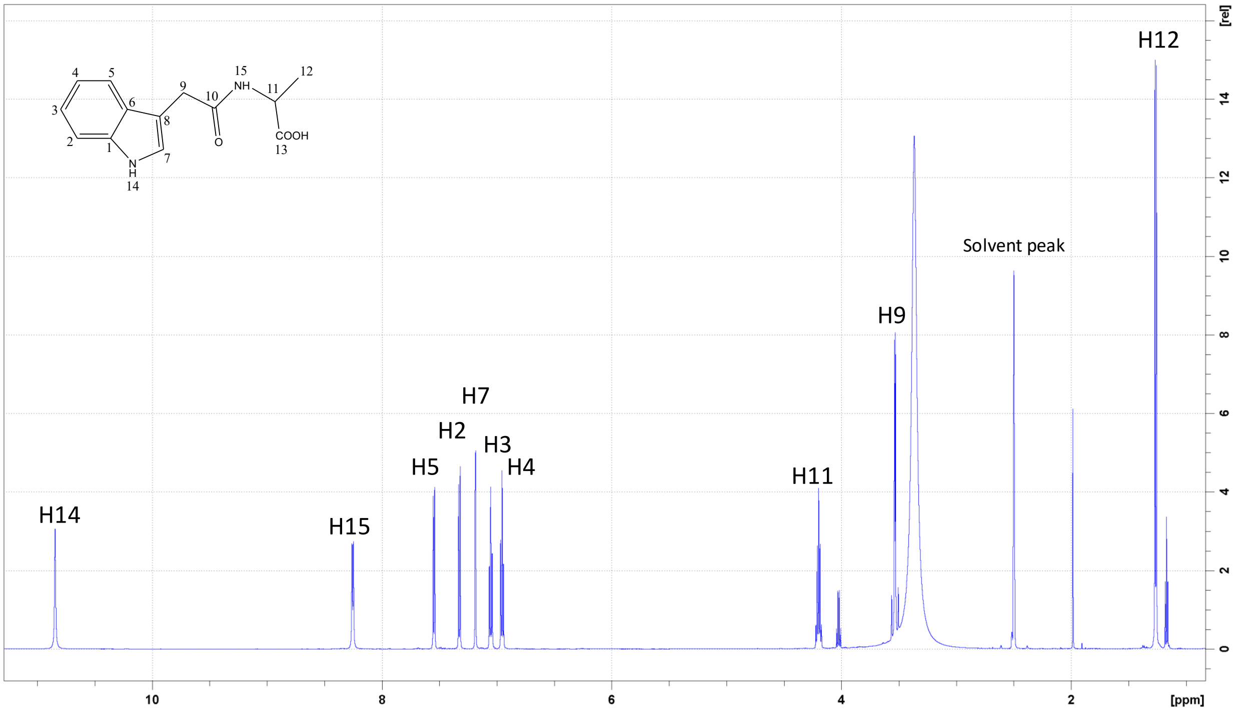


1D $^{1}H$ spectrum of IAA-alanine with assignments (in DMSO-d6)

IAA-alanine

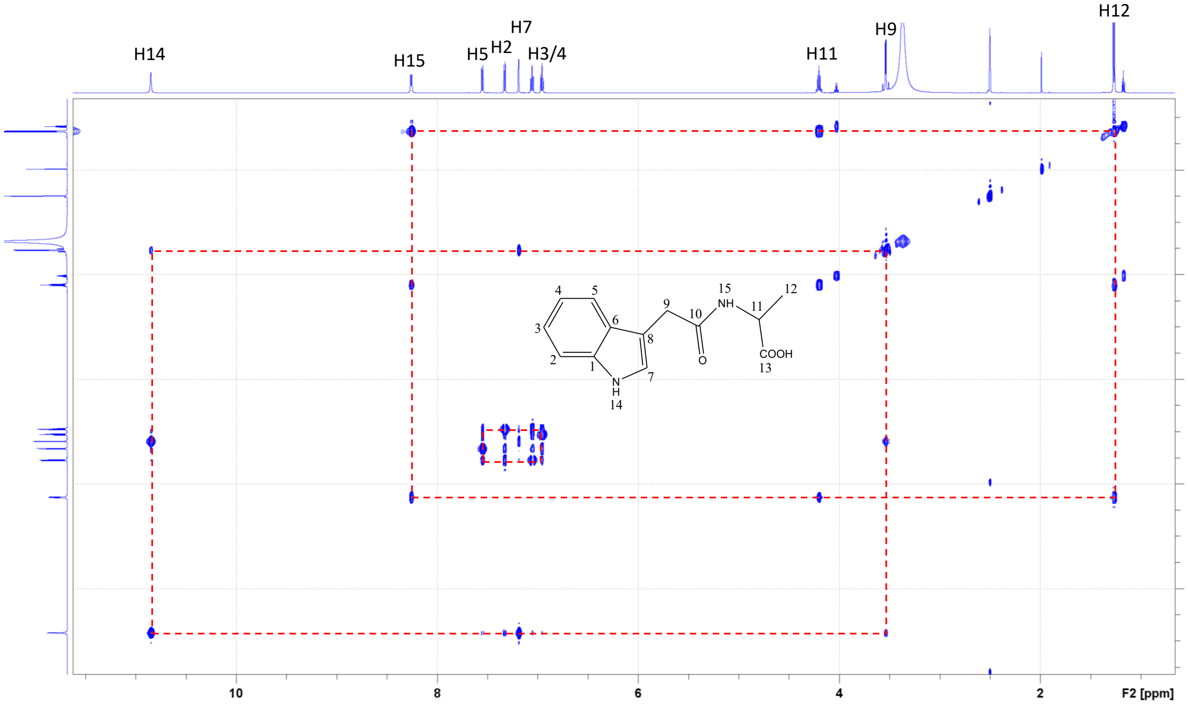


$^1H$-$^1H$ TOCSY spectrum of IAA-alanine with assignments (in DMSO-d6)

IAA-alanine

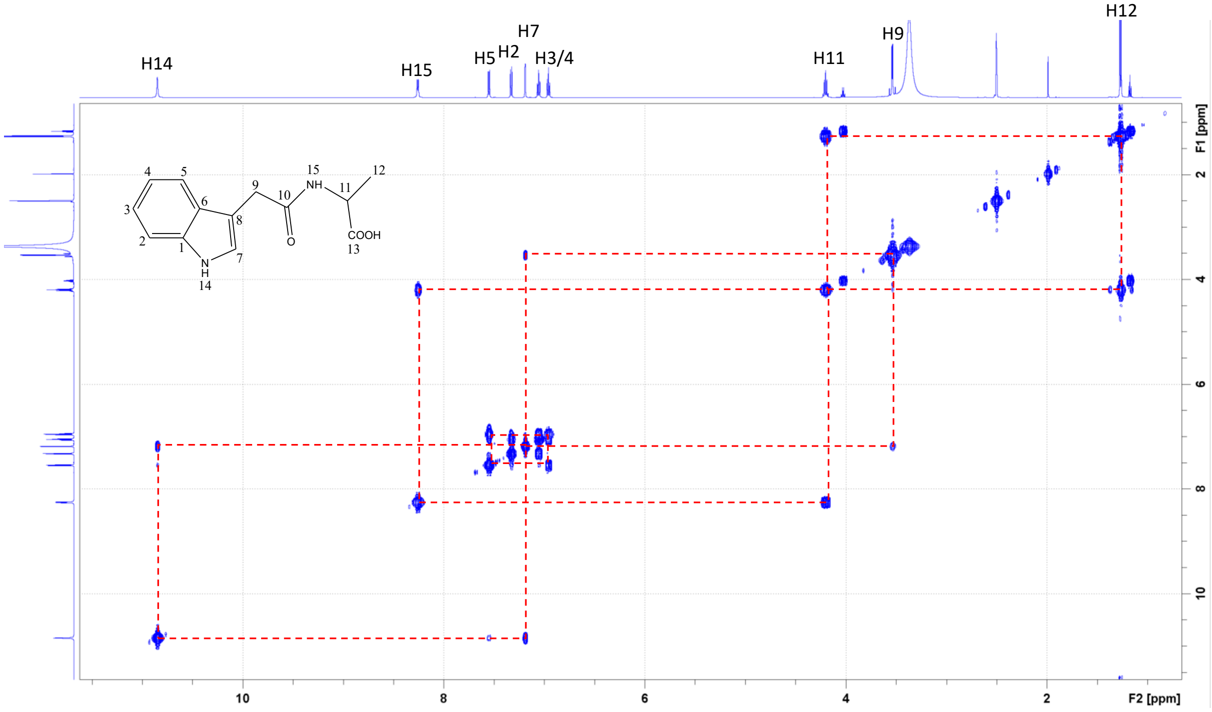


$^1$H-$^1$H COSY spectrum of IAA-alanine with assignments (in DMSO-d6)

IAA-alanine

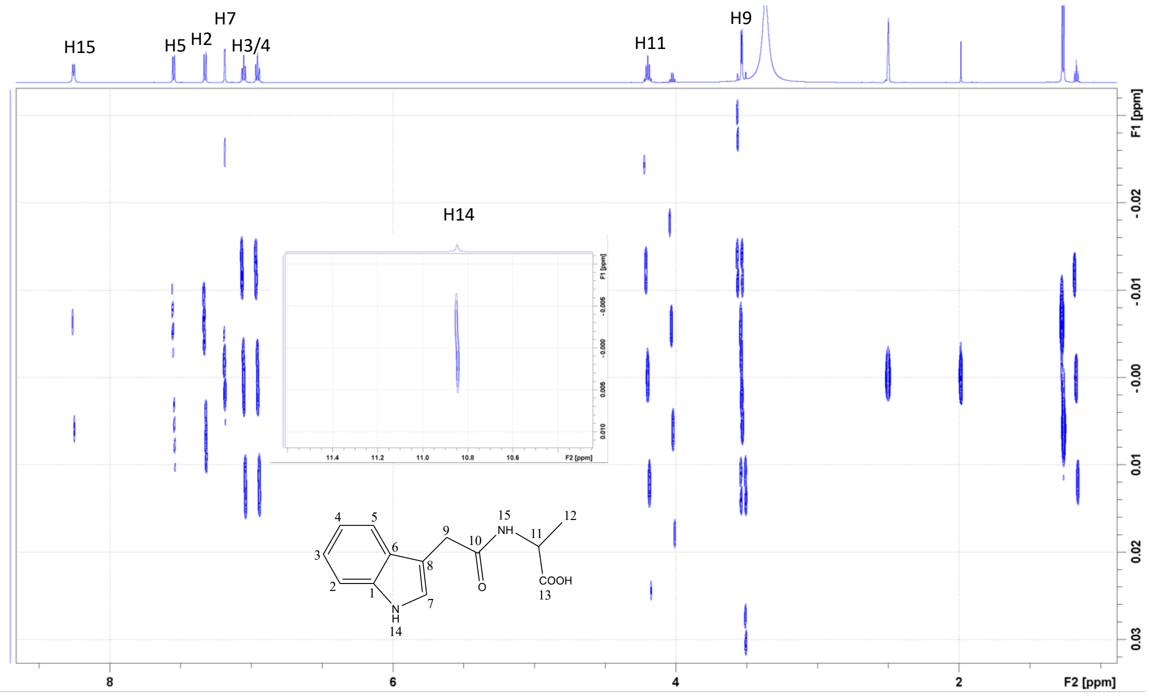


$^1$H-$^1$H JREs spectrum of IAA-alanine showing J-coupling patterns (in DMSO-d6)

IAA-alanine

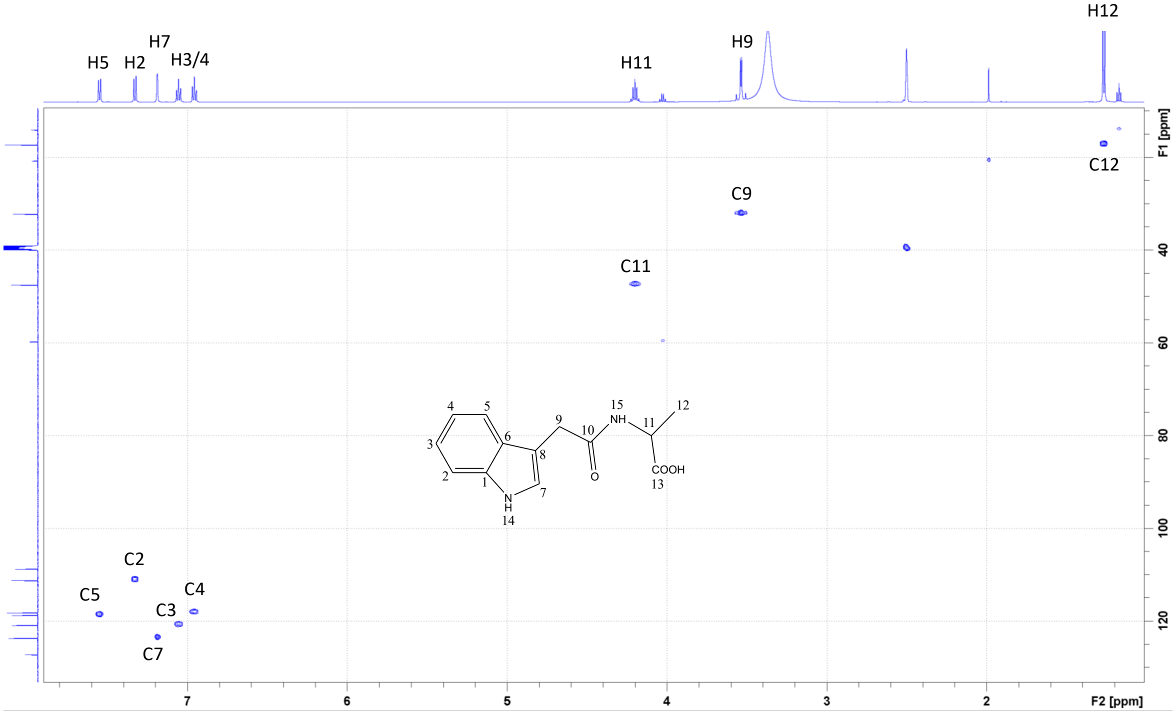


$^1H$-$^{13}C$ HSQC spectrum of IAA-alanine with assignments (in DMSO-d6)

IAA-alanine

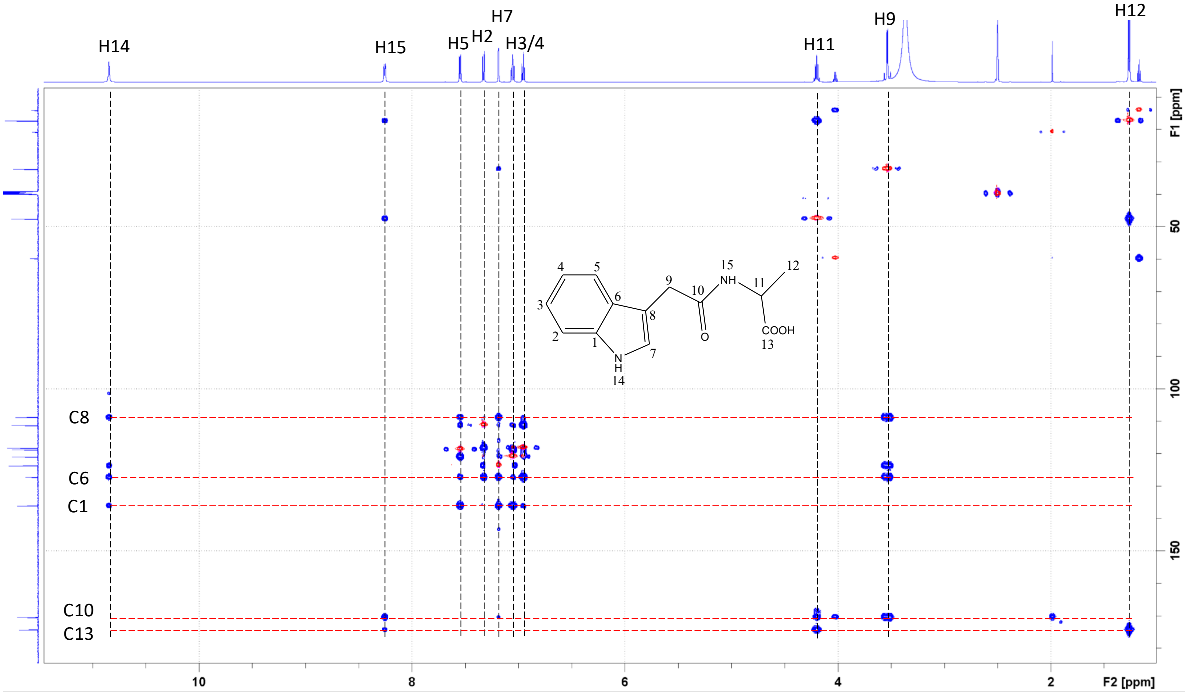


$^{1}H$-$^{13}C$ HMBC (blue) and HSQC (red) spectrum of IAA-alanine and signal assignments (in DMSO-d6)

IAA-glutamine

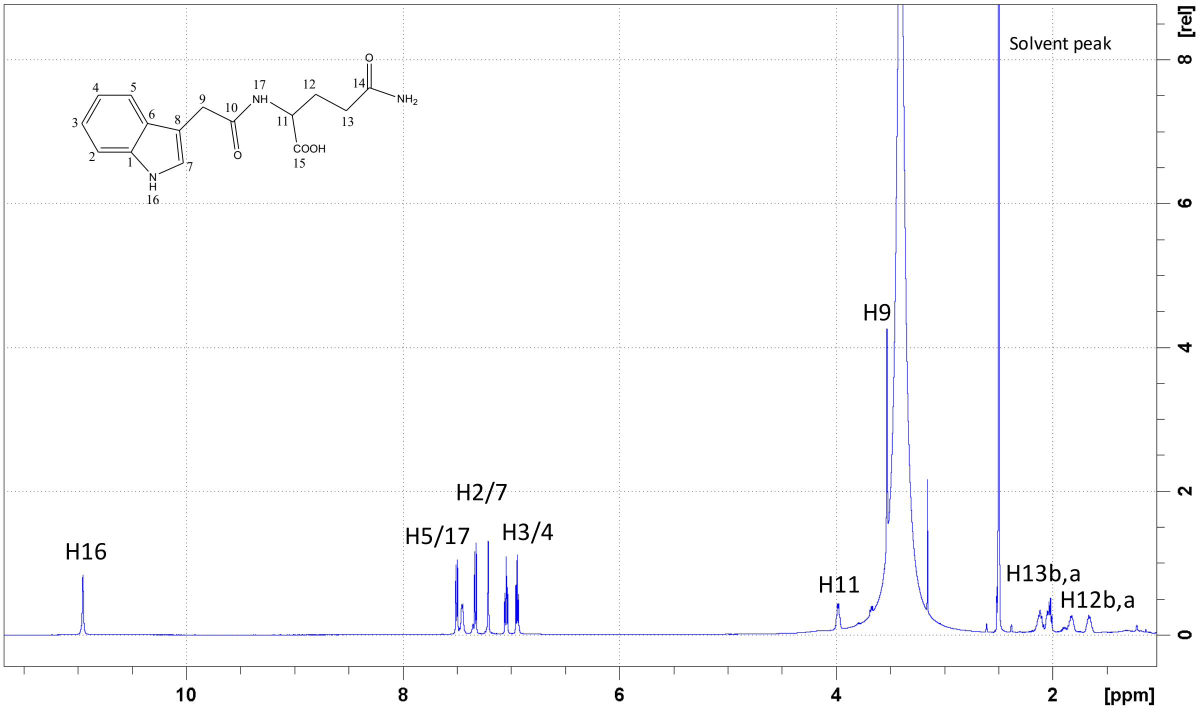


1D $^{1}H$ spectrum of IAA-glutamine with assignments (in DMSO-d6)

IAA-glutamine

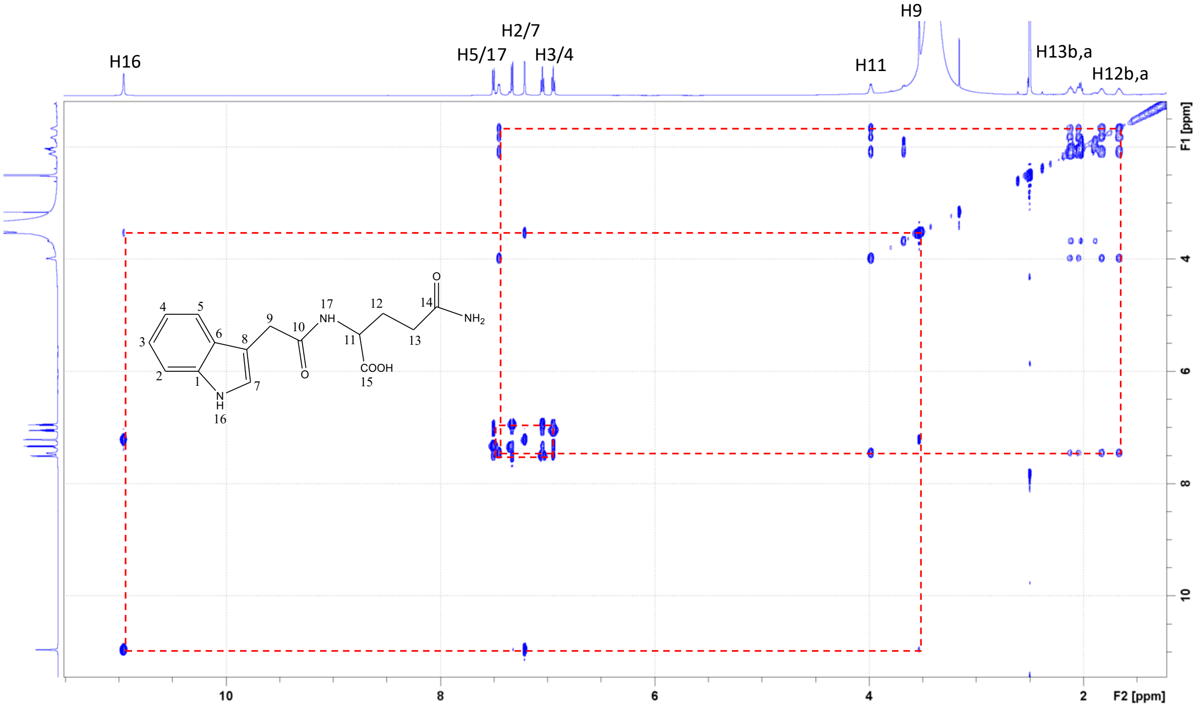


$^1$H-$^1$H TOCSY spectrum of IAA-glutamine with assignments (in DMSO-d6)

IAA-glutamine

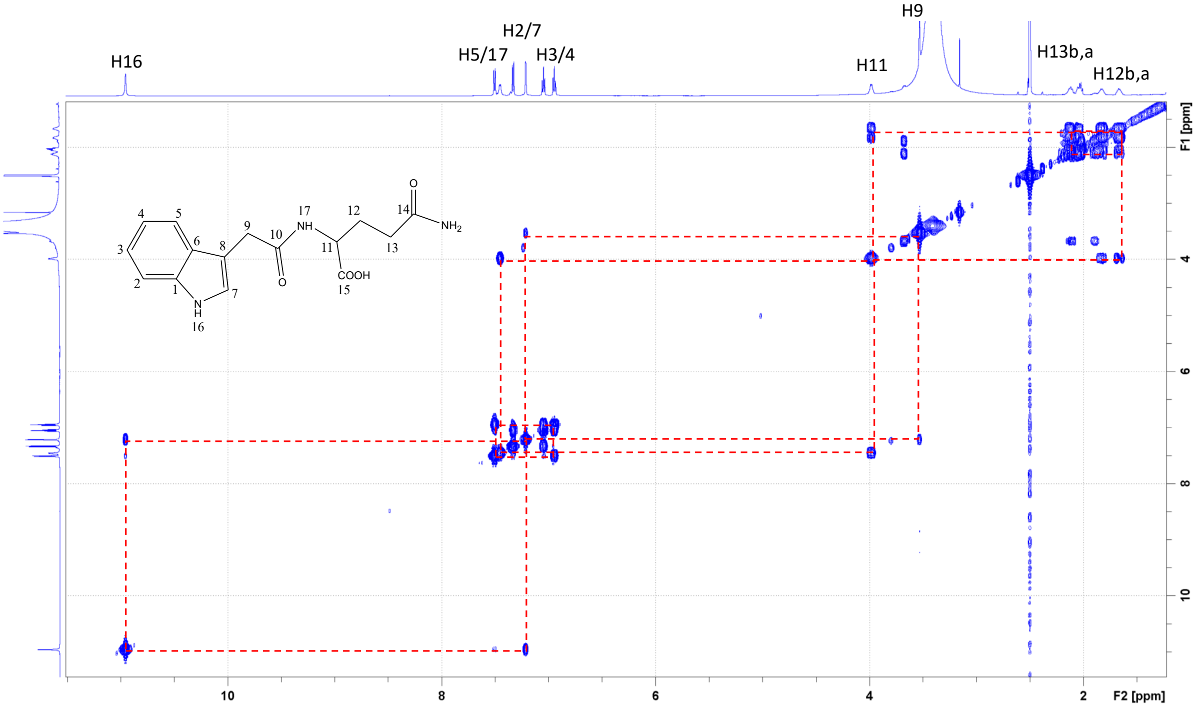


$^1H$-$^1H$ COSY spectrum of IAA-glutamine with assignments (in DMSO-d6)

IAA-glutamine

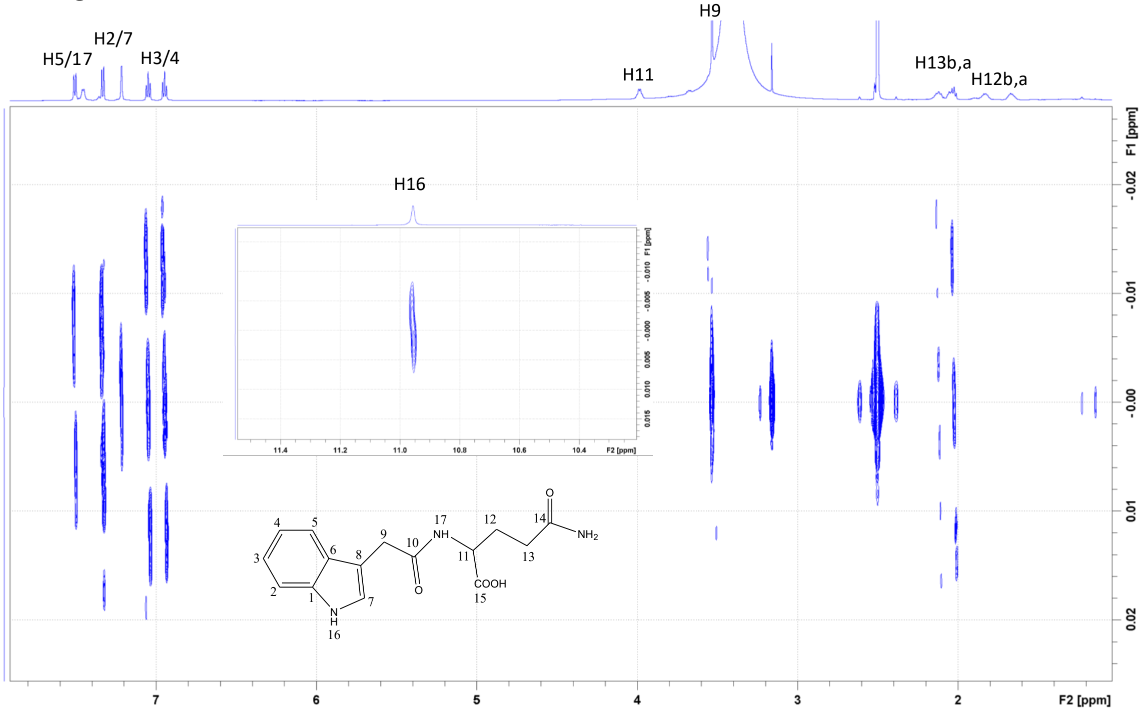


$^1H$-$^1H$ JREs spectrum of IAA-glutamine showing J-coupling patterns (in DMSO-d6)

IAA-glutamine

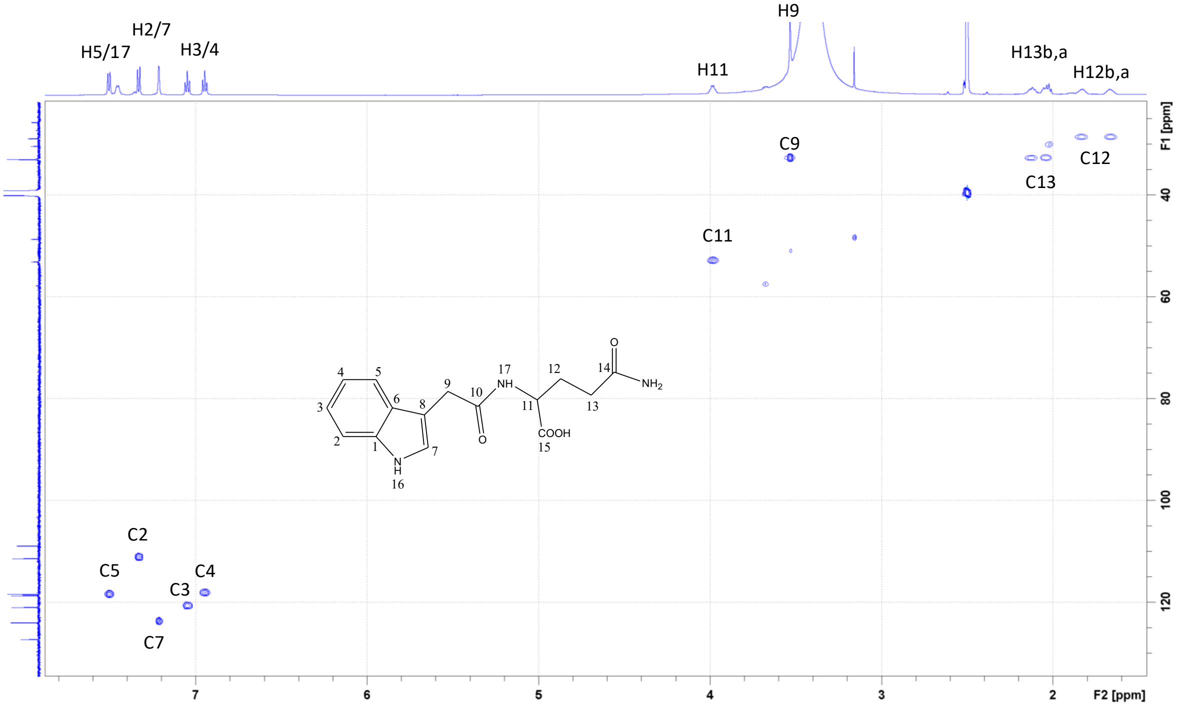


$^1H$-$^{13}C$ HSQC spectrum of IAA-glutamine with assignments (in DMSO-d6)

IAA-glutamine

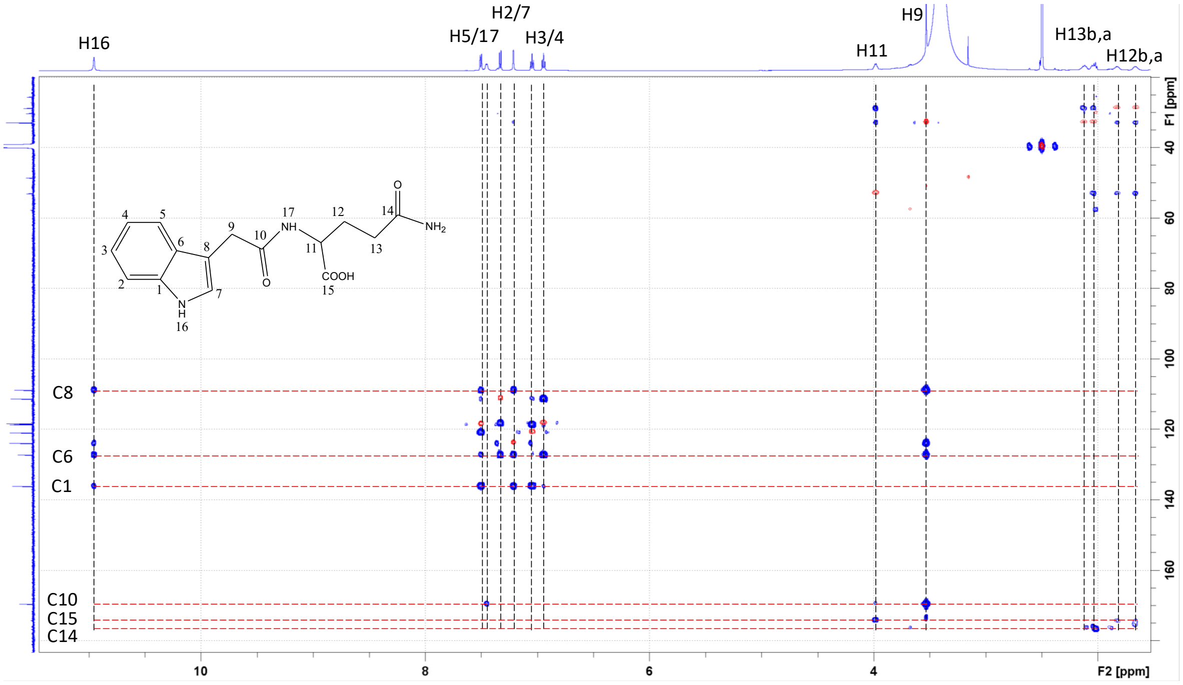


$^{1}H$-$^{13}C$ HMBC (blue) and HSQC (red) spectrum of IAA-glutamine and signal assignments (in DMSO-d6)

IAA-threonine

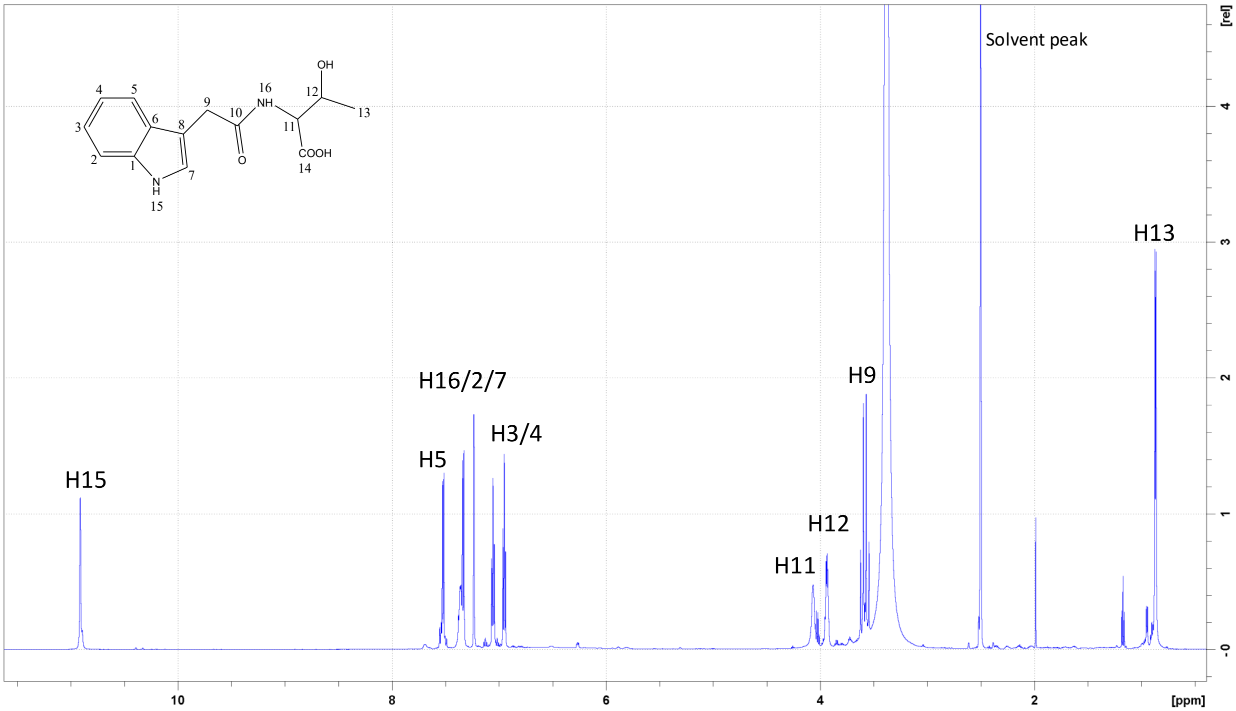


1D $^{1}H$ spectrum of IAA-threonine with assignments (in DMSO-d6)

IAA-threonine

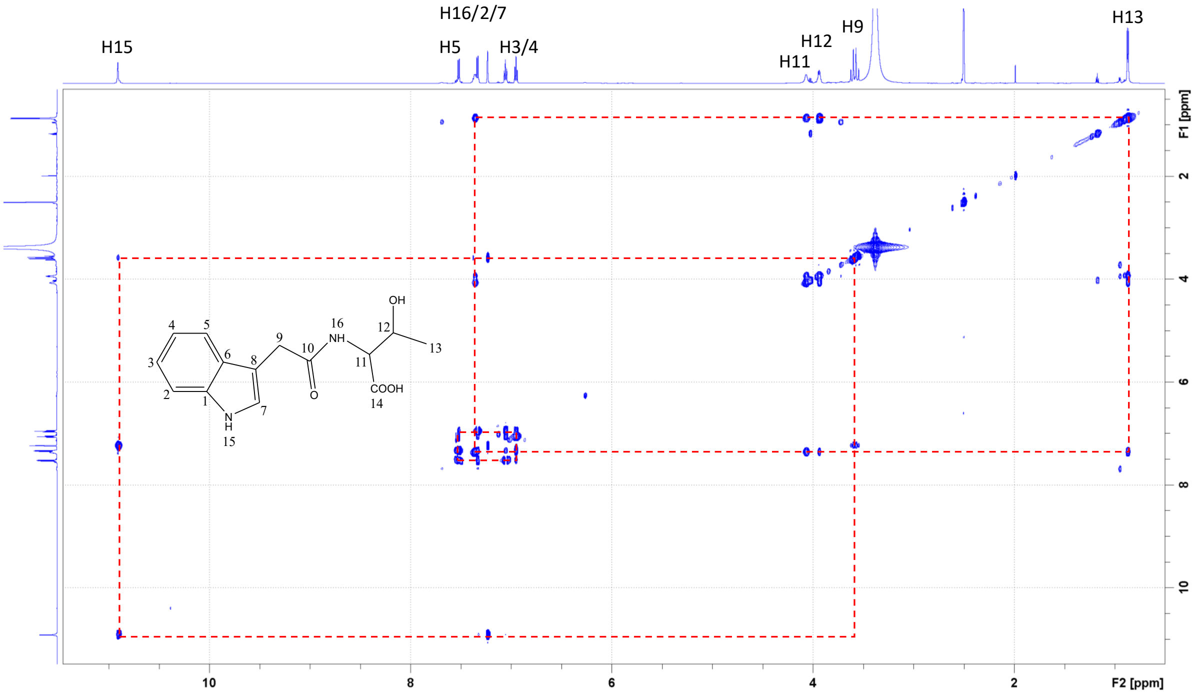

$^{1}H$-$^{1}H$ TOCSY spectrum of IAA-threonine with assignments (in DMSO-d6)

IAA-threonine

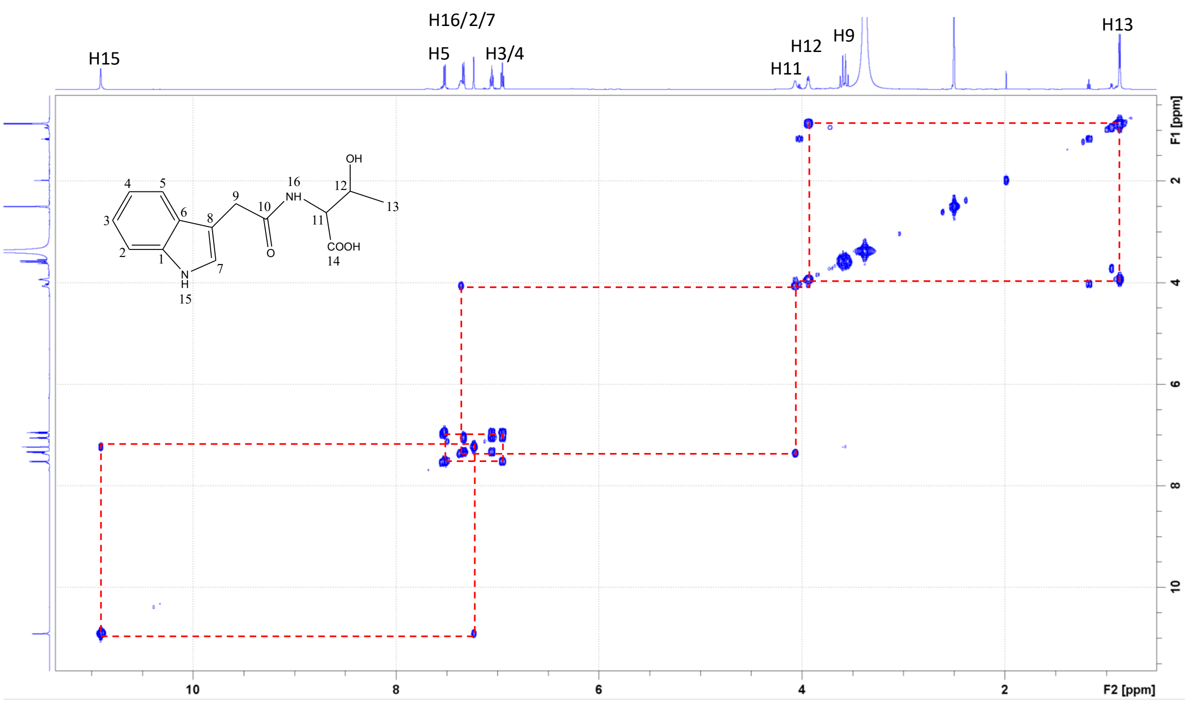


[1]H-[1]H COSY spectrum of IAA-threonine with assignments (in DMSO-d6)

#### IAA-threonine

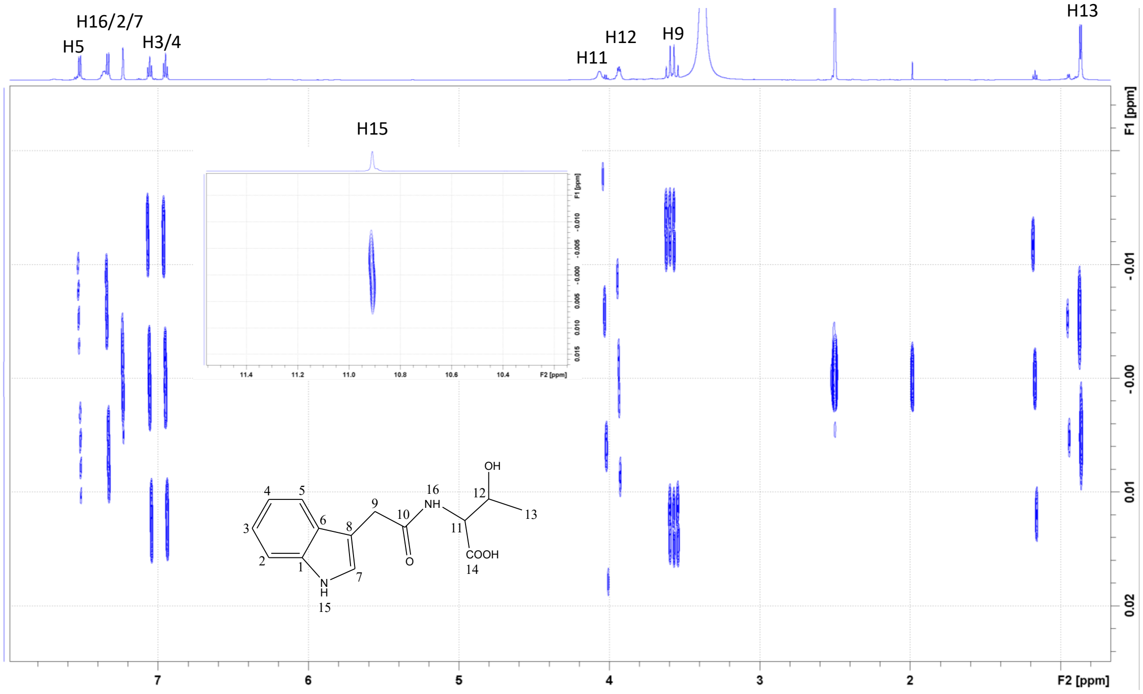


$^1$H-$^1$H JREs spectrum of IAA-threonine showing J-coupling patterns (in DMSO-d6)

IAA-threonine

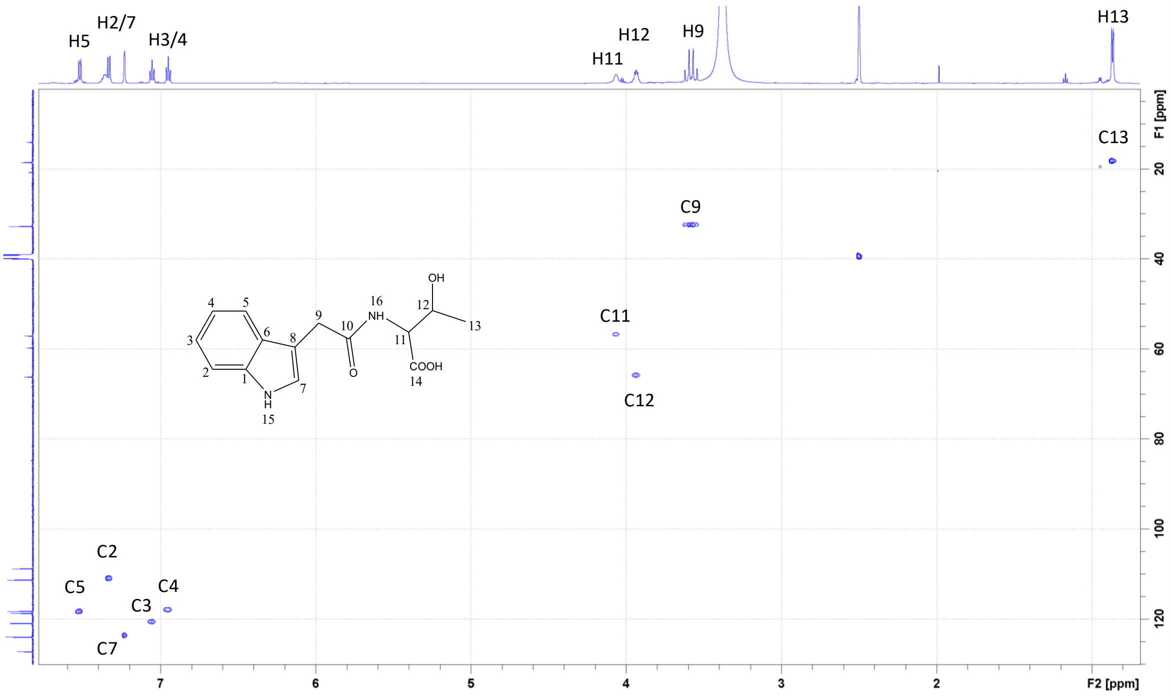


$^{1}H$-$^{13}C$ HSQC spectrum of IAA-threonine with assignments (in DMSO-d6)

IAA-threonine

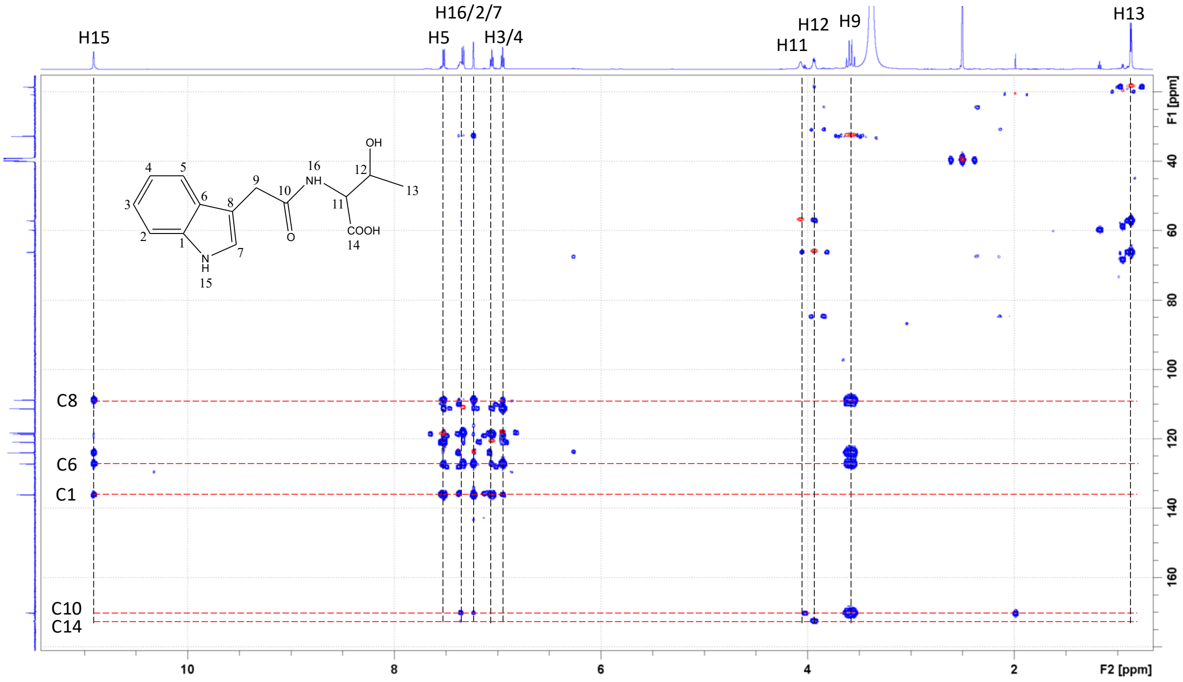


$^{1}H$-$^{13}C$ HMBC (blue) and HSQC (red) spectrum of IAA-threonine and signal assignments (in DMSO-d6)
